# Altered thymic niche synergistically drives the massive proliferation of malignant thymocytes

**DOI:** 10.1101/2024.06.25.600590

**Authors:** Erika Tsingos, Advaita M. Dick, Baubak Bajoghli

## Abstract

The discovery of genetic alterations in patient samples over the last decades has reinforced a cell-autonomous view of proliferative expansion during T-cell acute lymphoblastic leukemia (T-ALL) development in the thymus. However, the potential contribution of non-cell-autonomous factors, particularly the impact of thymic epithelial cells (TECs) within the thymic niche during the initiation phase, remains unexplored. In this study, we combine a cell-based computational model of the thymus with complementary *in vivo* experiments in medaka (*Oryzias latipes*) to systematically analyze the impact of 12 cell-autonomous and non-autonomous factors, individually and in combination, on the proliferation of normal and malignant thymocytes carrying interleukin-7 receptor (IL7R) gain-of-function mutations or elevated IL7R levels, as observed in T-ALL patients. By simulating over 1500 scenarios, we show that while a dense TEC network favored the proliferation of normal thymocytes, it inhibited the proliferation of malignant lineages, which achieved their maximal proliferative capacity when TECs were sparsely distributed. Our *in silico* model further predicts that specific mutations could accelerate proliferative expansion within a few days. This prediction was experimentally validated, revealing the rapid onset of thymic lymphoma and systemic infiltration of malignant T-cells within just 8 days of embryonic development. These findings demonstrate that synergistic interaction between oncogenic alterations and modifications in the thymic niche can significantly accelerate disease progression. Our results also suggest that negative feedback from the proliferative state suppresses thymocyte differentiation. Overall, this multidisciplinary work reveals the critical role of TEC-thymocyte interactions in both the initiation and progression of T-ALL, highlighting the importance of the thymic microenvironment in early leukemogenesis.

## INTRODUCTION

T-cell acute lymphoblastic leukemia (T-ALL) is a malignancy characterized by the proliferation of immature T cells, known as thymocytes, and is subclassified according to the stage of thymic maturation (Terwilliger and Abdul-Hay 2017; De Keersmaecker et al. 2013). Representing approximately 10-15% of pediatric cases and 25% of adult cases of all acute lymphoblastic leukemia (Vadillo et al. 2018), T-ALL originates in the thymus. This organ is the primary site where early T cell progenitors (ETPs) originating from the hematopoietic tissue differentiate into mature, immunocompetent T cells. Thymocytes follow a well-orchestrated migratory path within distinct thymic niches, receiving essential extrinsic signals for their proliferation, differentiation, and selection (Takahama 2006). Thymic epithelial cells (TECs) play a central role in this process, acting as the main cellular component of these niches, and influencing thymocyte migration and differentiation by expressing crucial growth factors, ligands (such as DLL4), or releasing chemokines (e.g. CCL25) and cytokines such as interleukin-7 (IL- 7) into the thymic niche (Yui and Rothenberg 2014; Koch et al. 2008; Zamisch et al. 2005).

The etiology of T-ALL is a topic of extensive research. Isolation and genome sequencing of malignant thymocytes from patients have uncovered mutations in over 100 putative driver genes (Liu et al. 2017; Girardi et al. 2017). Notably, over 50% of T-ALL patients harbor gain-of-function mutations in the NOTCH1 gene, which encodes the receptor for DLL4, while around 10% of patients exhibit similar mutations in the *IL-7 receptor* (*IL7R*) gene (Liu et al. 2017). The leukemogenic potential of constitutive activation of these receptors has been demonstrated across various vertebrate models (González-García et al. 2009; Silva et al. 2021; Shochat et al. 2011; Zenatti et al. 2011; Oliveira et al. 2022; Chen et al. 2007; Blackburn et al. 2012), emphasizing the evolutionary conservation of thymic developmental mechanisms that become misrouted in T-ALL (Bajoghli et al. 2019). These studies have contributed valuable insights into the impact of genetic alterations and oncogenic pathways on T-ALL development. However, the gene-centric view provided only limited understanding of the initiation and progression of the disease. In recent years, the growing evidence of the role of an abnormal tumor microenvironment in carcinogenesis strongly supports a more integrative view that considers the convergence of cellular genetics and the surrounding malignant niche as crucial elements for disease progression (Vadillo et al. 2018). In the context of T-ALL, however, the impact of the thymic niche or the role of TECs has not been studied.

Computational and mathematical models are powerful tools for integrating complex biological processes and can be used to test new hypotheses in health and disease (Ji et al. 2017; Metzcar et al. 2019; King et al. 2021). Several such models have been developed to study population dynamics during T cell development (Robert et al. 2021; Aghaallaei et al. 2021; Efroni, Harel, and Cohen 2007; Thomas-Vaslin et al. 2008; Vibert and Thomas-Vaslin 2017; Souza-e-Silva et al. 2009). Our recently developed cell-based computational model, for example, simulates individual thymocytes using agents within a spatially-resolved "virtual thymus" to explore cell-level behaviors and their effects on thymocyte population dynamics. This model was originally developed to investigate the impact of thymic niche signals and intrathymic cell localization on the αβ/γδ T cell sublineage outcomes (Aghaallaei et al. 2021).

In this study, we developed new computational tools to enhance the virtual thymus model, enabling us to identify both cell-autonomous and non-autonomous factors in thymocytes and TECs that could drive the proliferative expansion of a lineage derived from a single progenitor cell (hereafter referred to as a clone) within the thymus. We conducted an unbiased systematic analysis of various parameters, including TEC shape and architecture, IL-7 signaling, cell cycle, duration of the proliferative phase, and cell migration. This analysis facilitated a detailed comparison of proliferative expansion between wildtype (WT) and lesioned clones with alterations in the IL7R and NOTCH1 receptors. Through simulations of over 1500 scenarios, we identified factors that promote substantial expansion of malignant clones. In particular, modifications in the TEC network and niche emerged as a previously uncharacterized factor that could synergistically accelerate clonal expansion. Subsequent *in vivo* experiments using medaka fish (*Oryzias latipes*) confirmed the outcomes of our *in silico* analysis, solidifying the predictive accuracy of our virtual model, and revealing the impact of TECs on the initiation and progression of T-ALL.

## RESULTS AND DISCUSSION

### The virtual thymus model enables reconstructing clonal dynamics of medaka embryonic T cell development

The details of the virtual thymus model implementation and parameter estimation have been extensively described previously (see Supplementary Material of Aghaallaei et al. 2021), and in the following we will only delineate its main features. The model considers both spatial and temporal aspects of early thymus development at multiple scales: Cells are physically represented as one or multiple spheres that mechanically interact with adhesive and repulsive forces to capture cell crowding; each cell has an independent internal state that changes dynamically based on subcellular signaling modeled with ordinary differential equations or with phenomenological rules; and at the tissue level a partial differential equation is used to solve the diffusion of extracellular molecules such as cytokines. To reduce computational cost, the computational representation focuses on a 5 µm deep slice of the lower half of the radially symmetric organ, approximately 1/10 of the total volume of a medaka embryonic thymus (Figure 1 - Supplement 1A, B). Consistent with observations in vertebrates, including in medaka (Bajoghli et al. 2009; 2015; Aghaallaei et al. 2021; 2022), ETPs within our virtual thymus mechanically and biochemically interact with TECs, proliferate and differentiate in response to signals from the thymic niche, and subsequently diverge into two distinct T cell sublineages before selection and exit from the thymus (Figure 1A; see the Appendix and Aghaallaei et al. 2021 for more technical details). In this work, we define cells of a lineage derived from a single founder ETP as a “clone”, and we do not consider differences in immunological clonotypes that emerge after recombination of the T cell receptor.

**Figure 1.**
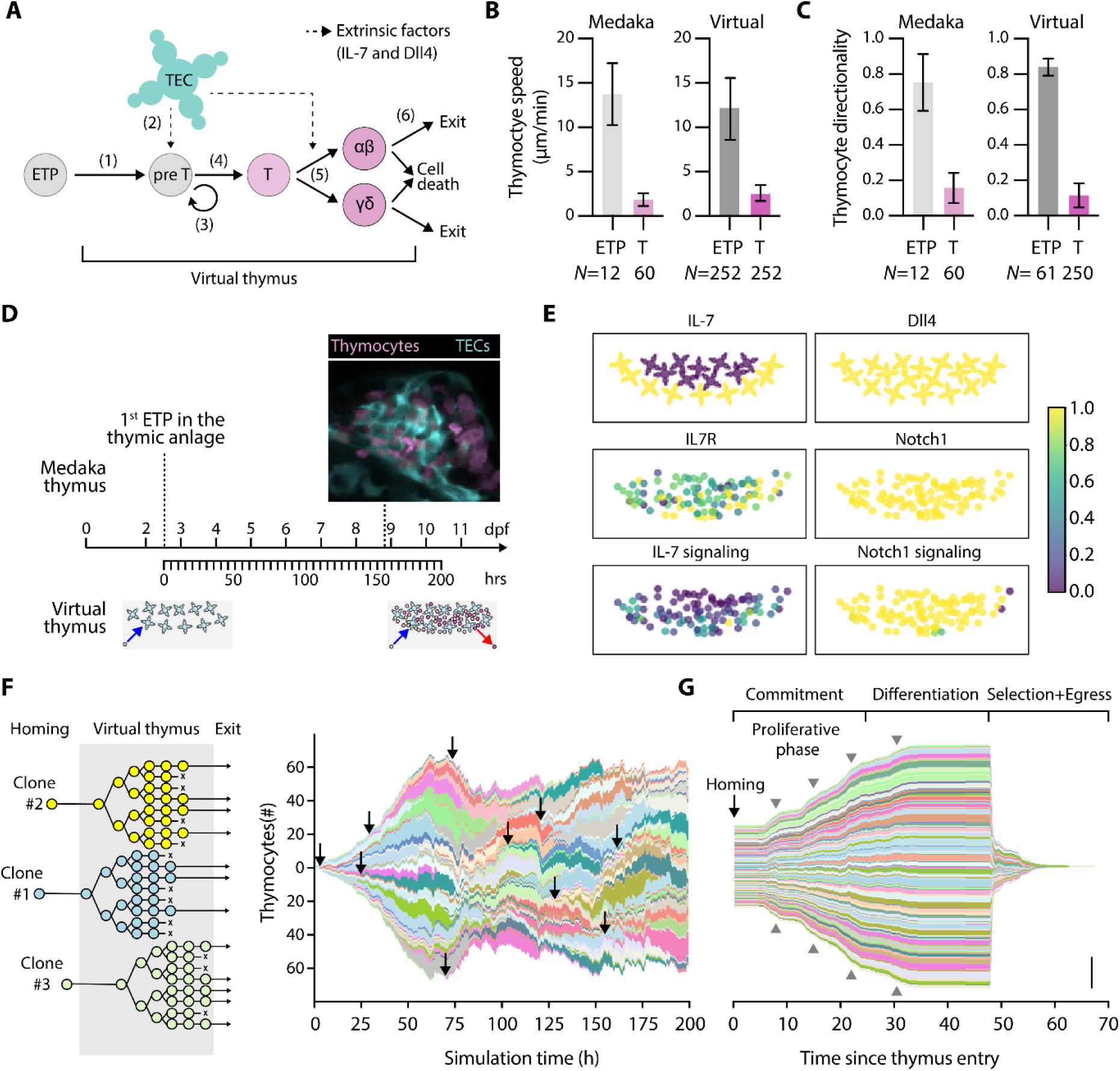
The virtual thymus model enables spatiotemporally resolved clonal analysis. (A) Schematic overview highlighting 6 key steps during T-cell development, which are integrated into the virtual thymus model: (1) ETPs enter the thymus niche; (2) they receive Dll4 and IL-7 signals from the TECs. Triggered by these signals, (3) thymocytes undergo proliferation and (4) differentiation into two distinct T-cell sublineages (5). Finally, thymocytes undergo selection, either leaving the organ or undergoing cell death (6). (B-C) Experimentally measured (“Medaka”) and computationally modeled (“Virtual”) cell speed (B) and cell directionality (C) for ETPs and thymocytes. Bars are mean values, and error bars standard deviation. (D) Timeline of medaka embryonic development (top) compared to simulated time (bottom). As indicated by the cartoon at the bottom, the first ETP enters the thymus roughly at 2.5 dpf (t = 0 h in the simulation). At roughly 60 h (roughly 5 dpf), the thymus reaches its homeostatic cell population, with both cell entry and exit. (E) Representative snapshots of the simulation at homeostasis showing spatial expression patterns and signaling activity of components of IL7R and Notch1 signaling pathways in TECs (top two panels) and thymocytes (bottom 4 panels). Data are shown in arbitrary units normalized to a scale from 0 to 1. (F) Left: Cartoon explaining the asynchronous arrival (homing) of new cells into the organ, their clonal expansion, and asynchronous cell death or exit. Right: Visualization of clonal diversity in a typical simulation; each color shade uniquely indicates a clonal lineage. The height of each colored patch indicates the number of cells in that clone, the length indicates simulation time. For illustrative purposes, a black arrow indicates the homing of a sample of lineages. (G) Data from the right panel in (F) was normalized to time of homing. This visualization highlights developmental phases of thymocyte lineages. Gray arrowheads indicate rounds of cell division. Scale bar 50 cells. Abbreviations: ETP = early thymic progenitor, T = Thymocyte, *N* = number of replicates (biological or computational).

In the virtual thymus model, factors governing thymocyte motility, including cell speed (Figure 1B) and directionality (Figure 1C) were fine- tuned based on quantitative noninvasive imaging of the thymus using multiple medaka transgenic reporter lines (Bajoghli et al. 2009; 2015; Aghaallaei et al. 2021). Based on these experimental observations, we modeled entry into the organ by creating new cells at the bottom of the simulated domain (ventral to the organ), these cells move into the thymus and, some time after differentiation, either leave the organ by exiting from the ventral side as a naïve T cell or undergo cell death by gradually shrinking and eventually being removed from the simulation (Figure 1 – Supplement Movie 1-2). The model accounts for the time taken for a single progenitor to enter and exit the thymus, which is estimated at approximately 3 days in zebrafish (Hess and Boehm 2012) and medaka (Bajoghli et al. 2015). For time calibration, each simulation step was set to represent 15 seconds. With a total of 48,000 steps per simulation, the entire simulation spans 720,000 seconds, equivalent to about 8.3 days (Figure 1D). This time interval was chosen to recapitulate thymic development in medaka from 2.5 to 11 days post-fertilization (dpf), because at 2.5 dpf the first ETP enters the embryonic thymus (Bajoghli et al. 2019). Correspondingly, each simulation starts with entering of the first ETP into the virtual thymus and ends with a homeostatic cell population of roughly 100 cells (mean ± standard deviation: 98±7; Figure 1 – Supplement Movie 1).

At 11 dpf, the medaka thymus contains 877±146 (N=7) thymocytes. Considering that the virtual thymus represents 1/10 of the medaka embryonic thymus, the cell population in the virtual thymus model reproduces the temporal dynamics in medaka within the margin of error.

Each cell’s internal state and decision to proliferate or differentiate depends on intrinsic and extrinsic factors that are integrated by signaling pathways. The virtual thymus includes Delta-like 4 (Dll4), the ligand of Notch1 receptor, and interleukin-7 (IL-7) cytokine as two cell-extrinsic factors provided by the TECs (Figure 1E). Consistent with medaka WT embryonic thymus (Aghaallaei et al. 2021; 2022), all virtual TECs in this model express the Dll4 (Figure 1E, top-right panel), with a subset spatially releasing IL-7 into the environment (Figure 1E, top-left panel), creating a short-ranged IL-7 cytokine gradient in the thymic cortex niche (Aghaallaei et al. 2021). In terms of cell-intrinsic factors, all ETPs uniformly express the Notch1 receptor (Figure 1E, middle-right panel), and each cell lineage expresses the IL-7 receptor (IL7R) at a constant level chosen randomly between 0 and 1 (Figure 1E, middle-left panel). In each simulation, new ETPs continually enter the thymic niche at regular intervals. Following engagement with TECs and receiving Notch1 and IL-7 signals (Figure 1E, bottom panels), they proliferate and differentiate (step 3 in Figure 1A). The cell cycle was modeled with an average duration of 7 hours and was subdivided into G1, S, G2, and M phases. In line with *in vivo* observations (Ruijtenberg and van den Heuvel 2016), virtual cells require external pro-proliferative signals during the G1 phase to commit to S through M. Once differentiated, cells can no longer enter the cell cycle. The dynamics of pro-proliferative signals and the proliferation stop after differentiation allow clones *in silico* to undergo up to four rounds of cell division (Aghaallaei et al. 2021). Thymic selection is modeled as a probabilistic event and is not regulated by signaling dynamics.

In this work, we implemented a code that enables us to meticulously trace the unique identity of each cell upon entering the virtual thymus and subsequently track the identity of its descendants (Figure 1F, left panel; Figure 1 – Supplement 1 C). This tool identified various waves of clones because of the constant influx of new ETPs and negative selection and efflux of fully differentiated T cells (Figure 1F, arrows in the right panel, Figure 1 – Supplement Movie 1). We note a slight variability in the clone sizes, which could arise from three processes. First, random variations in the expression level of IL7R impact IL-7 signal transduction and, consequently, proliferation (Aghaallaei et al. 2021). Second, due to random cell motility and cell crowding effects, certain cells could coincidentally remain in close contact with IL-7-secreting TECs, maximizing their exposure to pro-proliferative signals such as IL-7 and Notch ligand, and thus increasing their chances of entering the cell cycle. Third, each cell determines the duration of its next cell cycle by drawing an Erlang-distributed random variable with a mean of 7 hours and a standard deviation of ≅0.99 hours.

After normalizing all clones based on their time of entry into the thymus, the pattern of developmental phases became clear across all clones (Figure 1G). In this visualization, the now near-synchronous rounds of cell division across clones can be seen as bumps in the graph indicating clonal expansion (Figure 1G, gray arrowheads). During the commitment phase, which has a minimum duration of 24h but can be extended due to insufficient Notch signaling, thymocytes were competent for proliferation, leading to an increase in the total cell number per clone. In the differentiation phase, which has a fixed duration of 24h, clones reached a maximum size. Note that differentiation prevents further entry into the cell cycle but permits cells that passed G1 and therefore committed to S through M phases to complete their division. Finally, clones underwent selection and exit from the virtual thymus, thus leaving the simulation. Together, the enhanced virtual thymus model facilitated the investigation of the heterogeneity and temporal dynamics of individual clones originating from a founder progenitor cell.

### TEC architecture promotes thymocyte proliferation by modulating IL-7 availability

*In vivo*, individual TECs exhibit a distinctive star-shaped morphology, and protrusions of neighboring TECs contact each other forming a three-dimensional network (Figure 2A, left panel). Using confocal imaging, we estimated the relative area occupied by thymocytes (53±3%; N=4) and TECs (47±3%; N=4). A similar morphology and tissue density was replicated in our *in silico* model (Figure 2A, right panel), with thymocytes taking up on average 56±2% and TECs 44±2% of the volume in homeostasis. We wondered to what extent TEC morphology and density could impact thymocyte population size. This aspect is difficult to study experimentally and is, therefore, an ideal use case for the virtual thymus model. In a series of simulations, we combinatorically varied (i) the size of TECs, (ii) the number of their protrusions, and (iii) the TEC cell density (Figure 2B). The average homeostatic thymocyte population size in each simulation was then used as a readout. The results from 26 different tested conditions predicted that a higher density of larger TECs with more protrusions led to an almost two-fold increase in the number of thymocytes (200±13, N=3; scenario 26 in Figure 2C, Figure 2 – Supplement 1A bottom) compared to the reference condition (98±7, N=19; Reference in Figure 2C). Conversely, scenarios where these parameters were reduced, particularly a reduced TEC size, also diminished the homeostatic thymocyte population size (scenarios 1-14 in Figure 2C). The condition with the least amount of cells, scenario 1, had a reduced TEC size, but an unchanged number of protrusions and density. The condition diametrically opposite to scenario 26 was scenario 6, with a reduced TEC size, reduced number of protrusions, and reduced TEC density (Figure 2 – Supplement 1A top). In both scenario 1 (45±1, N=3) and scenario 6 (51±2, N=3), the thymic population size was only about half of that in the reference condition (98∓7, N=19). Statistical analysis confirmed that all three parameters and their interactions were significant predictors of total thymocyte population size (Figure 2 – Supplement 1B-C). The statistically significant interaction can be explained by the fact that higher TEC density increases the number of TECs, and thereby amplifies the effect of an increased TEC radius or number of protrusions. Thus, these three alterations of the thymic niche morphology act synergistically to modulate thymocyte population size.

**Figure 2.**
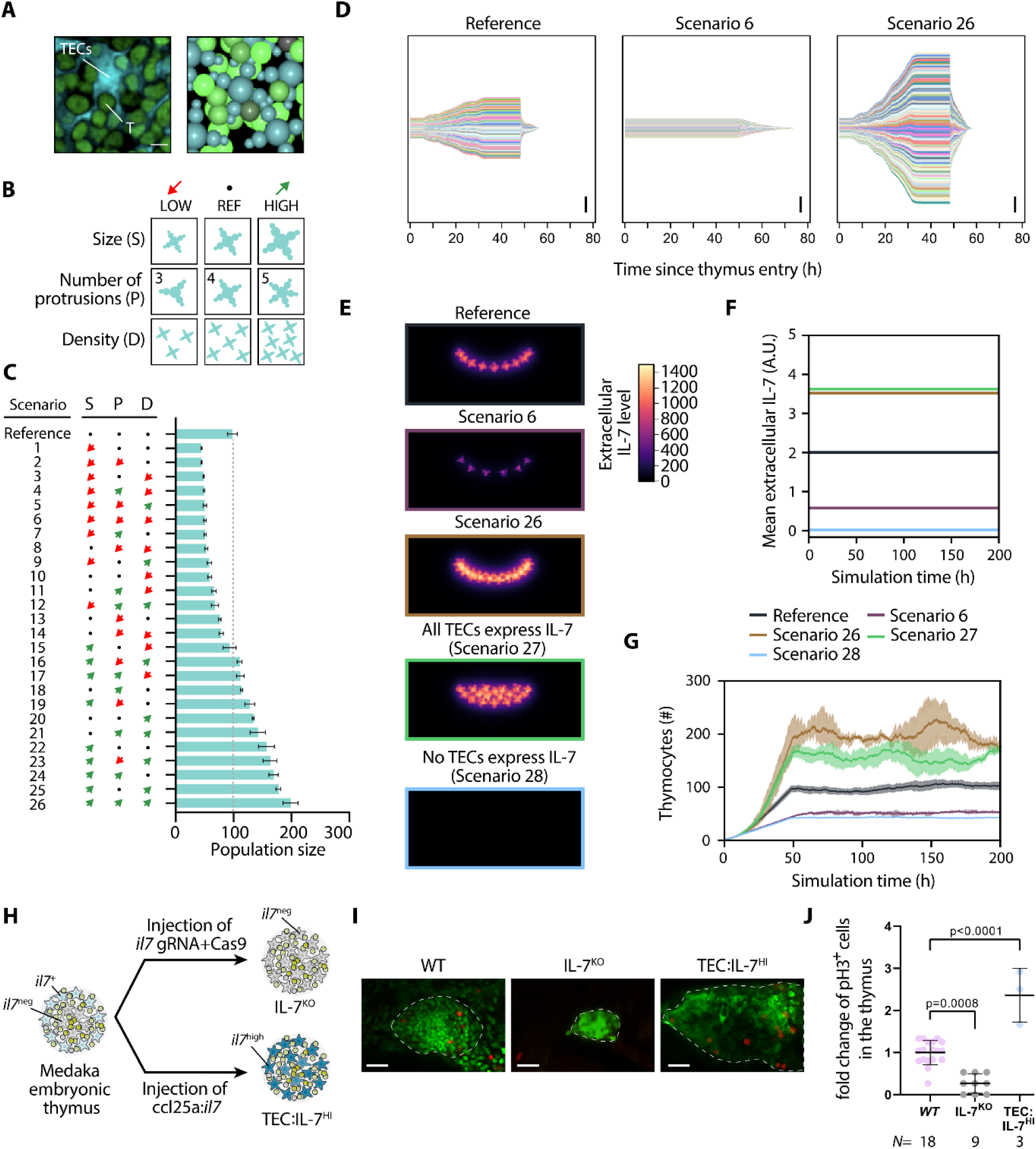
TEC density influences thymocyte proliferation via IL7R signaling. (A) Left: Detail of confocal section highlighting the tight spatial interaction between TECs and thymocytes (T). Scale bar 5 µm. Right: Detail from a three-dimensional render of a simulation snapshot; TECs cyan, thymocytes green. (B) Parameter permutations tested. (C) Impact of parameter permutations on average homeostatic cell population. Bars indicate means, error bars standard deviation. The abbreviations “S”, “P”, and “D” correspond to parameters “Size”, “Number of Protrusions”, and “Density”, respectively, as in panel (B). (D) Clonal lineages normalized to time of thymic entry for reference (left), a scenario with low average population (middle) and a scenario with high average population (right). Scale bar: 100 cells. (E) Extracellular IL-7 gradient in simulations; sum projection of all z-planes of a simulated confocal stack. Units are arbitrary. (F) Mean extracellular IL-7 concentration in the entire simulated volume. Note that these values are lower than in (E) because of averaging over the volume, including empty space (black areas in (E)). (G) Thymocyte population size over time averaged over several simulation runs. Shaded area indicates standard deviation. (H) Illustration of experimental setup for IL-7^KO^ and TEC:IL-7^HI^. At least 3 technical replicates were done for the injection of each construct. (I) Representative images of embryos stained with GFP (green) and pH3 (red) of WT, IL-7^KO^ and TEC:IL-7^HI^. Scale bar 20µm. (J) Numbers of pH3 positive cells per thymus normalized to mean of WT. *N* represents the number of individual fish. Data show means ± standard deviation.

There are two possibilities that could explain the variations in total thymocyte numbers in simulations. Either over- or under-proliferation of a few thymocyte lineages, or the cumulative impact of subtle changes across several thymocyte clones. To distinguish between these possibilities, we used our clonal analysis tool for scenarios 6 and 26 (Figure 2D). This analysis revealed that variations in total thymocyte numbers were a consequence of changes in proliferation across all clones to a similar degree. These results predict that the TEC architecture has a direct impact on thymocyte proliferation rate, which fits well with the fact that TECs act as the main source of ligands, growth factors, cytokines, and chemokines (Gameiro, Nagib, and Verinaud 2010). Any changes in the TEC architecture could therefore influence the amount of cytokine production. In our virtual thymus model, this affects the extracellular spatial distribution of IL-7 cytokine (Figure 2E). To further explore this, we tested two additional scenarios. In scenario 27, TECs maintained their reference architecture and density but all expressed IL-7, leading to a more uniform cytokine distribution in the thymic environment (Figure 2E). The extracellular level of IL-7 (Figure 2F) and thymocyte population size (155±11, N=5; Figure 2G) in scenario 27 was similarly high as in scenario 26, where the size, protrusion number, and density of TECs were increased. Conversely, when TECs did not express IL-7 (scenario 28), the thymocyte population size (Figure 2G) was low (42±1, N=5), akin to scenario 6, where the size, protrusion, and density of TECs were reduced. Interestingly, although the average extracellular IL-7 level is at its highest in scenario 27 with ubiquitous expression, the spatially restricted elevated IL-7 concentration in scenario 26 is more effective at driving thymic population growth (Figure 2E-G). Thymocytes first enter the organ at its periphery from below (at the ventrolateral site), where IL-7 levels are highest, and tend to migrate to the IL-7-depleted center of the thymus as they become non-proliferative (Aghaallaei et al. 2021; see also Figure 1 – Supplement Movie 1). Thus, in scenario 26, thymocytes in their proliferative phase immediately encounter elevated IL-7 at the organ periphery, stimulating cell proliferation. In contrast, in scenario 27, thymocytes that enter the organ periphery encounter comparable levels of IL-7 to the reference scenario, and only after entering deeper into the organ can the ubiquitous expression of IL-7 unfold its effects. This delay contingent to random cell migration (which is further exacerbated by cell crowding posing an obstacle), reduces the impact of elevated IL-7 in scenario 27. These results highlight how the spatial structure of the thymic niche can impact on thymocyte population dynamics.

Our simulations predict that the availability of IL-7 provided by TECs has a direct impact on thymocyte population size. To validate this prediction *in vivo*, we conducted two functional analyses using the medaka model organism (Figure 2H). In one experiment, employing the CRISPR-Cas9 technique, we knocked out the *il7* gene (Figure 2- Supplement 2A, B) in a medaka transgenic line, where thymocytes express green fluorescent protein (GFP). Consistent with our *in silico* outcomes, *il7* crispant embryos displayed fewer thymocytes and a smaller thymus size (Figure 2I, Figure 2 - Supplement 2C, D). To estimate the extent of cell proliferation, we counted mitotic cells throughout the entire thymus using the M-phase marker phospho-histone 3 (pH3) and normalized values to the mean of the control. This analysis further supported that a reduction in IL-7 availability in the thymus reduces thymocyte proliferation (Figure 2I, J). In a second experimental setup (Figure 2H), we artificially increased *il7* levels in the thymic niche by injecting the ccl25a:*il7* construct into embryos (hereafter called TEC:IL-7^HI^), resulting in a strong upregulation of the levels of *il7* produced by the TECs (Aghaallaei et al. 2021). Compared to WT, the thymus of TEC:IL-7^HI^ embryos appeared visibly larger, and the number of pH3+ cells was significantly increased (Figure 2I, J). Therefore, the *in vivo* results confirm the critical role of IL-7 for thymocyte proliferation in medaka, a mechanism that is evolutionarily conserved among vertebrates (Iwanami et al. 2011). Additionally, the alterations in thymic size suggest a regulatory crosstalk between thymocyte proliferation and TEC architecture.

### Cell-autonomous sensitivity to IL-7 signaling promotes thymocyte proliferation

Given the important contribution of extracellular IL-7 on thymocyte population size, we next evaluated the impact of (i) extracellular IL-7 depletion (e.g., via ligand internalization), (ii) the rate of signal transduction activation upon IL7R and IL-7 binding, and (iii) the rate of decay of signaling activity (Figure 3A, B). In our virtual thymus model, the IL-7 signaling activity (*σ*_IL7_) of a thymocyte over time (*t*) is modeled phenomenologically using a function involving the IL7R concentration ([IL7R]), the average extracellular IL-7 concentration (〈[IL-7_ex_]〉), a parameter (*a*_IL-7_) that scales the strength of signal transduction activation, and another parameter (*d*_IL-7_) that scales the rate at which the signaling activity diminishes over time (Figure 3B). As before, we combinatorically perturbed these three factors, simulating 17 different scenarios, to assay their impact on thymocyte population size (Figure 3C). As expected, we observed a positive correlation between the level of signal transduction activation *a*_IL-7_ and thymocyte numbers. Conversely, variations in signaling decay rate *d*_IL-7_ showed the opposite effect. The rate of signal transduction deactivation models cellular short-term memory, thus a lower signaling deactivation rate indicates that cells retain their IL-7 stimulus for a longer duration. Because IL-7 signaling activity *σ*_IL7_ promotes cell cycle entry (see Appendix section *X. Subcellular Scale: Cell proliferation model* for an in-depth explanation), we expect that parameter changes that increase stimulus duration should promote cell division. Indeed, the scenario with the highest number of thymocytes occurred when signaling deactivation was low and activation was high (274±11, N=4; scenario 45 in Figure 3C). The diametrically opposite combination had the least amount of cells (42±1, N=4; scenario 29 in Figure 3C). Statistical analysis confirms that signal transduction activation *a*_IL-7_ , and the signal transduction deactivation rate *d*_IL-7_ were both statistically significant predictors of thymocyte population size (Figure 3 - Supplement 1). There was no statistically significant effect of parameter interactions, likely because these parameters exert their effect independently. There was only a limited reduction in population size with IL-7 depletion (91±11, N=4; scenario 36 in Figure 3C) compared to no depletion in our reference setting (98±7, N=19). Overall, IL-7 depletion did not have a significant effect on thymocyte population (Figure 3 - Supplement 1). However, we noted that population size difference was more pronounced at higher cell numbers, e.g., comparing between scenario 44 (237±7, N=4) and scenario 45 (274±11, N=4). Similarly, the mean extracellular IL-7 content in the virtual thymus barely changed when comparing the reference to scenario 36 but was noticeably reduced in scenario 44 compared to scenario 45 (Figure 3D). Therefore, thymocyte proliferation could self-inhibit via depletion of extracellular IL-7, but we expect this effect to be small unless cell numbers are massively increased.

**Figure 3.**
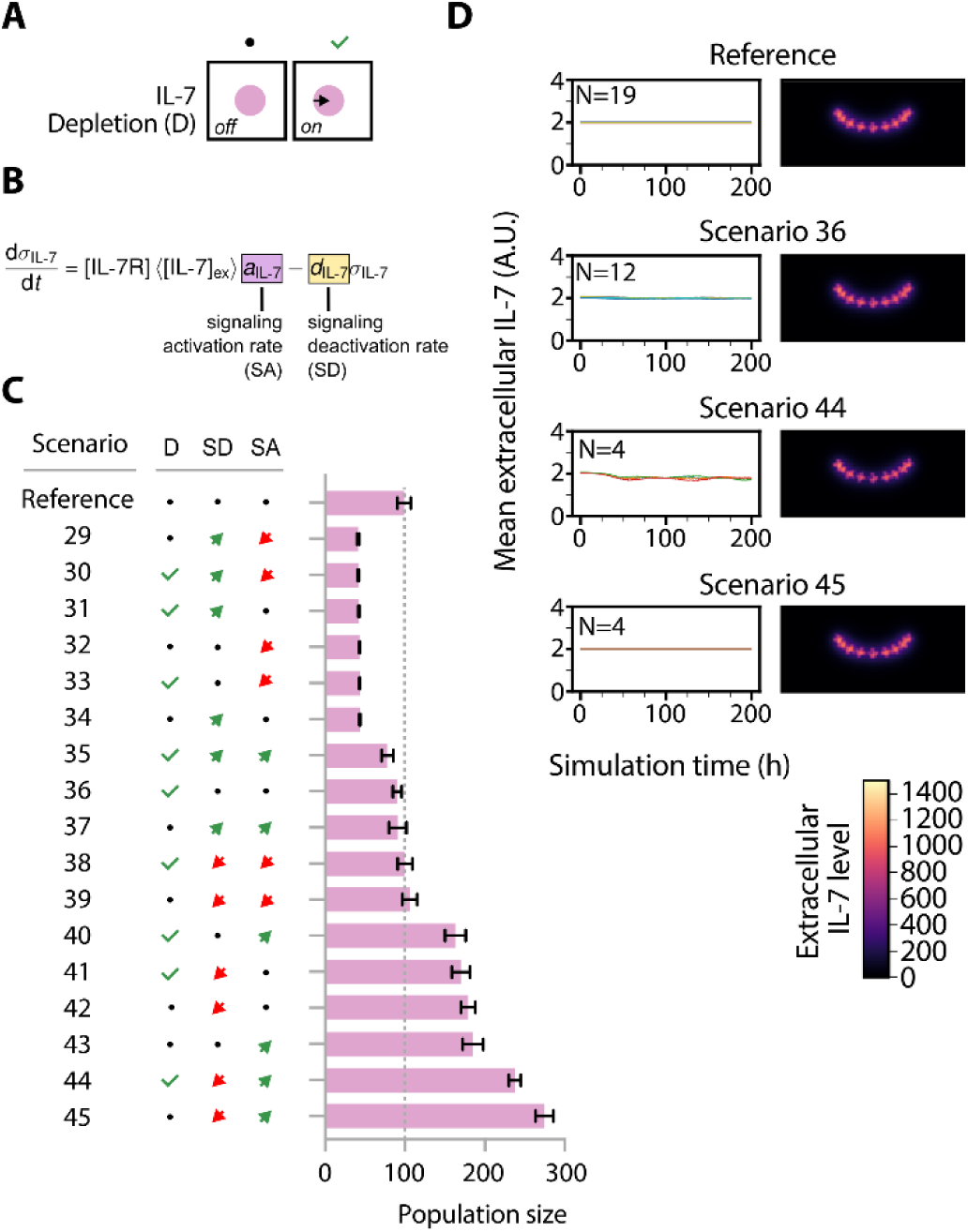
High IL7R signaling promotes proliferation. (A) Illustration of scenarios without extracellular IL-7 depletion (left) and with extracellular IL-7 depletion (right). (B) Ordinary differential equation used to model the IL7R signaling activity. Parameters that were modified are highlighted. (C) Effect of IL7R signaling-related parameter permutation. Bars indicate means and error bars standard deviation. The abbreviations “D”, “SD”, and “SA” correspond to “Depletion”, “Signaling Deactivation Rate”, and “Signaling Activation Rate”, respectively, as indicated in panels (A) and (B). (D) Left: Mean extracellular IL-7 concentration in the entire simulated volume. Right: Extracellular IL-7 gradient in simulations; sum projection of all z-planes of a simulated confocal stack. Units are arbitrary. The introduction of IL-7 depletion by thymocytes leads to a slight reduction in extracellular IL-7 availability.

Together, the outcomes of our simulations reveal that molecular and cellular changes, specifically those capable of increasing the extrinsic factor IL-7 in the niche — such as a higher density of TECs or an elevated expression level of IL-7 by individual TECs — directly contribute to an increase in thymocyte population size. Tuning thymocytes’ sensitivity to IL- 7 by manipulating IL7R signaling can further amplify proliferation. Finally, we expect that cytokine internalization by thymocytes has only a mild inhibitory effect on excessive population growth by limiting the availability of the pro-proliferative IL-7 in the extracellular space.

### A systematic approach identifies TEC architecture as a synergistic factor promoting clonal expansion of IL7R-lesioned clones

Our 45 tested scenarios thus far reached a maximum three-fold induction of thymocyte population size when parameter changes promoted increased IL-7 signaling. While the activation of IL-7 signaling is linked to T- ALL development, this pathology is associated with proliferation of clones carrying somatic mutations (Oliveira et al. 2019; 2022; Silva et al. 2021). We therefore modified the code of our virtual thymus model to enable the introduction of parameter changes exclusively in a single clone and all of its progeny (hereafter referred to as the lesioned clone). Inspired by observations that 10% of T-ALL patients exhibit dominant active mutations in the *IL7R* gene (Liu et al. 2017; Shochat et al. 2011; Zenatti et al. 2011), and a significant subset of T-ALL patients with active IL7R signaling display elevated *IL7R* expression levels (Silva et al. 2021), two types of lesions were introduced: (1) a dominant active IL7R lesion (hereafter called IL7R^DA^ lesion), (2) an IL7R overexpression lesion (hereafter called IL7R^HI^ lesion). For the IL7R^DA^ lesion, we set the IL7R level of the lesioned clone to the maximum WT level of 1 and allowed it to activate IL-7 signaling regardless of extracellular IL-7 cytokine, mimicking the dominant activation mutations in the *IL7R* gene (Figure 4A, middle panel). For the IL7R^HI^ lesion, we mimicked overexpression of the *IL7R* gene by assigning a value of 10 for the IL7R level in the lesioned clone, i.e. 10-fold higher than the maximum attainable in the WT population (Figure 4A, right panel). In both scenarios, non-lesioned thymocytes retained reference IL7R levels randomly ranging between 0 to 1 (Aghaallaei et al. 2021). In each simulation, a single lesioned clone entered the thymic niche after the 60th hour of a simulation, that is shortly after the establishment of a homeostatic population size (Figure 4B, Figure 4 - Supplement Movie 1).

**Figure 4.**
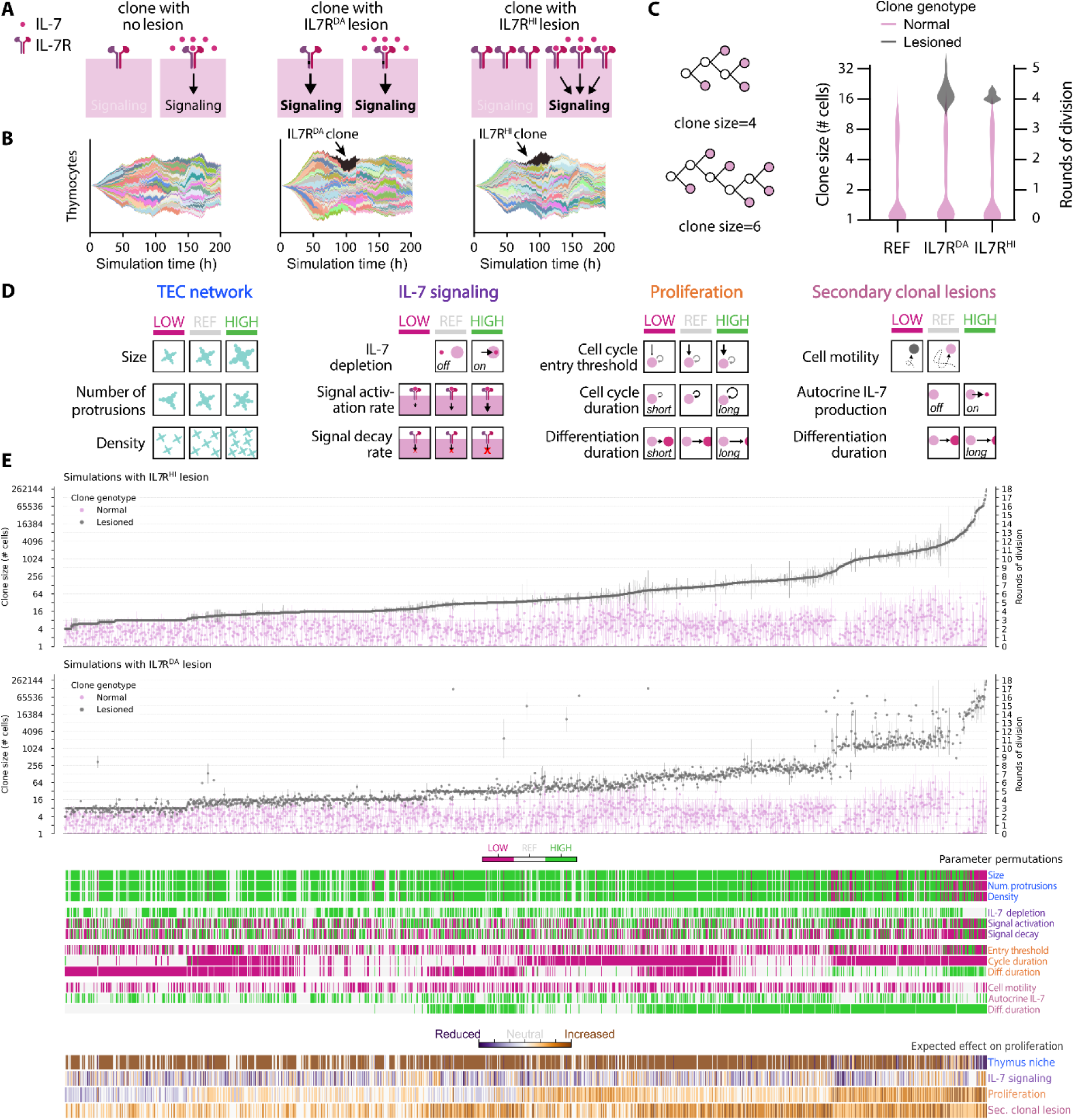
Systematic screen shows that lower TEC density promotes malignant expansion. (A) Schematic representation of the major lesions to IL7R that were implemented in the model. (B) Representative clonal plot highlighting in black the lesioned clone. (C) We defined clone size as the maximum number of terminal leaves of a lineage, regardless of eventual fate of each cell. Clone size distribution in the reference simulation (REF), compared to IL7R^DA^ and IL7R^HI^. REF: N=20 simulations, with 2088 clones total: IL7R^DA^: N=13 simulations with 1330 non-lesioned clones and 13 lesioned clones. IL7R^HI^: N=13 simulations with 1336 non-lesioned clones and 13 lesioned clones. (D) Schematic of tested scenarios shown in (E). (E) Simulated permutations ordered along increasing mean lesioned clone size for the IL7R^HI^ condition. Expected effect on proliferation is a qualitative estimate based on preliminary simulations. A total of 1580 permutations were tested with a varying amount of replicates per permutation. Error bars indicate standard deviation. Note that in (C) and (E) the axis is in base-2 logarithmic scale to better illustrate rounds of cell division.

First, we evaluated the impact of IL7R^HI^ and IL7R^DA^ lesions in the reference condition, meaning no additional modifications were made to our model. The clonal analysis of simulations revealed that introducing a lesioned clone did not markedly alter the division behavior of non-lesioned clones in the same simulation (scenario with no lesion: 1.04±1.26 rounds of cell division; scenario with IL7R^DA^: 0.98±1.22 rounds of cell division; scenario with IL7R^HI^: 1.04±1.23 rounds of cell division; Figure 4B-C). The large variance in non-lesioned clones is consistent with our previous work (Aghaallaei et al. 2021), which showed that clones expressing endogenously high levels of IL7R undergo more rounds of cell division in the virtual thymus, whereas clones with lower levels of IL7R proliferate very little or not at all. In contrast, lesioned clones averaged just above 4 rounds of cell division (4.20±0.34 for IL7R^DA^ and 4.10±0.13 for IL7R^HI^). Consequently, the size of IL7R^DA^ and IL7R^HI^ clones was nearly 8-fold larger than an average non-lesioned clone (Figure 4C). This result suggests that lesioned clones acquired a distinct proliferative advantage over non-lesioned clones in our virtual thymus model. Nevertheless, since the expansion of lesioned clones amounted to, at best, only one additional round of cell division compared to the upper range of the non-lesioned clone distribution (Figure 4C), our virtual thymus model predicts that a single lesion in the *IL7R* gene will not be clinically impactful.

Therefore, we next explored whether additional modifications in both cell-autonomous and non-autonomous factors might substantially enhance the clonal expansion. To identify these factors, we decided to undertake a systematic approach and tested combinations of scenarios affecting the TEC architecture and IL-7 signaling parameters, as tested in Figures 2 and 3, together with changing other parameters affecting proliferation and differentiation of all clones (Figure 4D). Furthermore, we considered scenarios in which only lesioned clones exhibited additional modifications, such as reduced cell motility, slower differentiation time, or autocrine IL-7 production. The inclusion of the latter was inspired by a study showing that certain malignant T cells derived from T-ALL patients possess the ability to ectopically express IL-7 cytokine (Buffière et al. 2019). In theory, there are 104976 possible permutations of these scenarios. To reduce the space of possibilities and thus the computational cost, we prioritized alterations expected to increase proliferative potential based on outcomes shown in Figures 2 and 3. We also decided to focus on simulating the extremes; e.g., low and high levels shown in Figure 4D. In addition, we simulated a small sample of scenarios expected to decrease proliferative potential, such as a reduction in TEC density. In the end, 1580 scenarios were simulated. The clonal analysis tool was used to compare clone size and the number of cell division rounds between lesioned and non-lesioned clones under the same conditions. Overall, we observed a similar proliferation pattern among IL7R^HI^ and IL7R^DA^ clones in most of the tested scenarios. Likewise, non-lesioned clones displayed similar trends as clones in control simulations that lacked a lesioned clone (Figure 4 - Supplement 1). The lesioned clones consistently showed a higher proliferation rate than their non-lesioned counterpart (Figure 4E; Figure 4 – Source Data 1). This trend persisted even when parameters known to generally increase cell division for all cells, such as mean cycle duration and proliferation duration, were modified (Figure 4 - Supplement 2, Figure 4 – Supplement Movie 2). In fact, lesioned clones disproportionally gained up to 6 rounds of cell division by modulating proliferative parameters, while normal clones only gained 2 rounds of cell division in the same scenarios. This difference results from cell crowding creating a disproportional disadvantage to non-lesioned clones: In the scenario with increased proliferation, the first clones to colonize the organ can massively proliferate and completely surround the TECs, preventing later arriving clones from accessing TEC-derived pro-proliferative signals (Figure 4 – Supplement Movie 2, right column). Thus, these few “lucky” early colonizers prevent late cells from attaining their full proliferative potential. Lesioned clones carry an intrinsic pro-proliferative advantage due to their lesions in the IL7R, enabling proliferation despite a lack of direct contact with TECs. In contrast, modifying parameters that affected IL-7 activity only enhanced the proliferation rate of non-lesioned clones but showed no impact on either IL7R^HI^ or IL7R^DA^ clones (Figure 4 - Supplement 3). While this outcome is expected for IL7R^DA^ clones, which are insensitive to IL-7, for IL7R^HI^ clones it indicates that 10-fold receptor overexpression is enough to activate downstream pro-proliferative effects of the pathway regardless of the parameter permutations that we tested. In contrast, proliferation of normal clones is strongly affected in these conditions, gaining up to 3 rounds of cell division between least and most proliferative conditions.

In our systematic approach, we did not detect an added effect on clonal expansion from combining reduced cell motility or autocrine IL-7 production with IL7R^HI^ or IL7R^DA^ lesions (Figure 4 - Supplement 4). Added IL- 7 secretion by lesioned clones with the autocrine lesion had only a small effect on proliferation of non-lesioned clones in the same simulation. The most critical lesion in conferring a proliferative advantage to lesioned clones was a delay in differentiation. This delay essentially doubled the duration of the proliferative phase, and indeed we observed up to 8 rounds of cell division, double the level of clones having only a single lesion in the IL7R. This result fits well with *in vivo* data suggesting that perturbations in differentiation have been implicated in contributing to the initiation and progression of cancer (Ruijtenberg and van den Heuvel 2016).

Surprisingly, the expansion of lesioned clones was amplified when the TEC niche was at its sparsest – a result opposite to the non-lesioned clones which instead profit from a dense TEC niche (Figure 4 - Supplement 5). While non-lesioned clones gained 2 rounds of division with an increase in TEC density, lesioned clones lost 2 rounds of division. Indeed, parameter combinations with reduced density and size of TECs were overrepresented in the scenarios that most increased proliferation (Figure 4 - Supplement 6A-C). For example, in the most extreme scenario observed in our systematic approach, IL7R^HI^ or IL7R^DA^ clones underwent up to 18 rounds of cell division – far above the 12 rounds predicted from modifying each group of parameters in isolation (+6 from proliferation, +4 from added lesions, +2 from reduced TEC density, assuming an additive effect; Figure 4 – Supplement 2-5). This result suggests that a defective niche amplifies the effect of other pro-proliferative modulations such as slower differentiation and shorter cell cycle, conferring a synergistic advantage to IL7R^HI^ and IL7R^DA^ clones compared to their normal counterparts. Note that at 18 rounds of division lesioned clones were so massive that the total lesioned cell volume was over a 1000-fold larger than the volume of the simulated organ slice (Figure 4 - Supplement 6D). In comparison, in the homeostatic condition, the total volume occupied by both thymocytes and TECs amounted to only 0.7-fold of the available tissue volume. Though our simulations by default include physical volume exclusion effects that prevent dense packing, we did not implement density-dependent feedback on proliferation, enabling cells to duplicate their volume at cell division and proliferate uncontrolled despite the lack of available space. Presumably, similar conditions *in vivo* would lead to thymus hyperplasia. In support of this view, our experimental manipulations of IL-7 levels in the thymus had a direct effect on organ size (Figure 2I, Figure 2 - Supplement 2C, D).

Identifying a sparse TEC network as a new factor that could influence the clonal expansion of lesioned clones was an unexpected outcome. This is because, in the WT situation, a sparser TEC network results in decreased extracellular IL-7, leading to reduced clonal size (Figure 2). While being disadvantageous for non-lesioned clones, this compromised thymic niche proved to be optimal for a massive expansion of lesioned clones. Conversely, we observed that a denser TEC network – characterized by a higher density of TECs with more protrusions and larger size – positively influenced the population size of non-lesioned cells but had a negative impact on the IL7R^HI^ and IL7R^DA^ lesioned clones (Figure 4 - Supplement 5). Similarly, in conditions of high cell crowding, lack of contact with TECs put non-lesioned clones at a disadvantage, while lesioned clones appeared to benefit (Figure 4 – Supplement 2, Figure 4 – Supplement Movie 2). This outcome might be explained by the signaling dynamics within our model implementation, where the engagement of the NOTCH1 receptor on thymocytes with the DLL4 ligand on TECs, on the one hand, promotes proliferation and, on the other hand, is a prerequisite for the differentiation process until commitment to a non-dividing fate. In the model, if the NOTCH1 receptor fails to engage with DLL4 on TECs due to their low density or cell crowding effects, cells will remain in an undifferentiated state for a longer time. Closer inspection of time-normalized clones in scenario 6 (very sparse TECs) and in scenario 26 (very dense TECs) indeed confirms that the thymocyte population has a longer turnover time when TECs are sparse (almost 80h, Figure 2D middle panel), and a shorter turnover when TECs are dense (less than 60h, Figure 2D right panel), indicative of the effect of Notch signaling. Together, cells in an environment with fewer TECs and thus lower Notch signaling will experience a prolonged proliferative phase but still require pro-proliferative signals to commit to the cell cycle. Owing to the higher levels of IL7R signal intrinsic to the lesion, lesioned clones will be more easily competent to proliferate even without Notch signaling and, therefore, benefit from a lower TEC density.

### The interplay between IL7R and NOTCH1 signals in the clonal expansion of thymocytes

In our virtual thymus model, ETPs simultaneously assess whether the combined sum of IL7R and NOTCH1 signals surpasses a threshold required for entry into the cell cycle ((Aghaallaei et al. 2021); Figure 5A). Besides its effect on proliferation, NOTCH1 signaling also promotes ETP differentiation (Aghaallaei et al. 2021; 2022), and we used this fact and the observations in *notch1b* medaka mutant phenotypes to include an accelerating effect of NOTCH1 on differentiation in our model (Aghaallaei et al. 2021; see also Appendix). Thus, NOTCH1 has a dual effect in thymocytes, which we implemented in our model as an incoherent feed-forward loop (Figure 5A): NOTCH1 signaling promotes proliferation and accelerates differentiation, while differentiation inhibits proliferation by preventing further entry into the cell cycle. IL7R signaling promotes proliferation independently of NOTCH1 and has no direct or indirect effect on the differentiation process. To further explore the interplay between these two factors, we compared scenarios where lesioned clones exhibited only constitutive activation of the NOTCH1 receptor (hereafter called NOTCH1^DA^) or in combination with either IL7R^DA^ or IL7R^HI^. The simulations predicted that lesioned clones with only NOTCH1^DA^ modification displayed a very small advantage compared to non-lesioned clones; this increase in proliferation was only marginally higher than expected for a clone expressing the highest endogenous levels of IL7R (hereafter IL7R^WT^; Figure 5B). In contrast, lesions that specifically delay differentiation led to higher proliferation. Indeed, the addition of NOTCH1^DA^ modification to IL7R^DA^ or IL7R^HI^ lesioned clones did not show any effect on clonal expansion but combining delayed differentiation with IL7R^DA^ or IL7R^HI^ doubled proliferation. This result indicates that modulating the effect of NOTCH1 globally does not affect proliferation in our model, which is likely due to feedback inhibition, but specifically targeting the differentiation process leads to clonal expansion.

**Figure 5.**
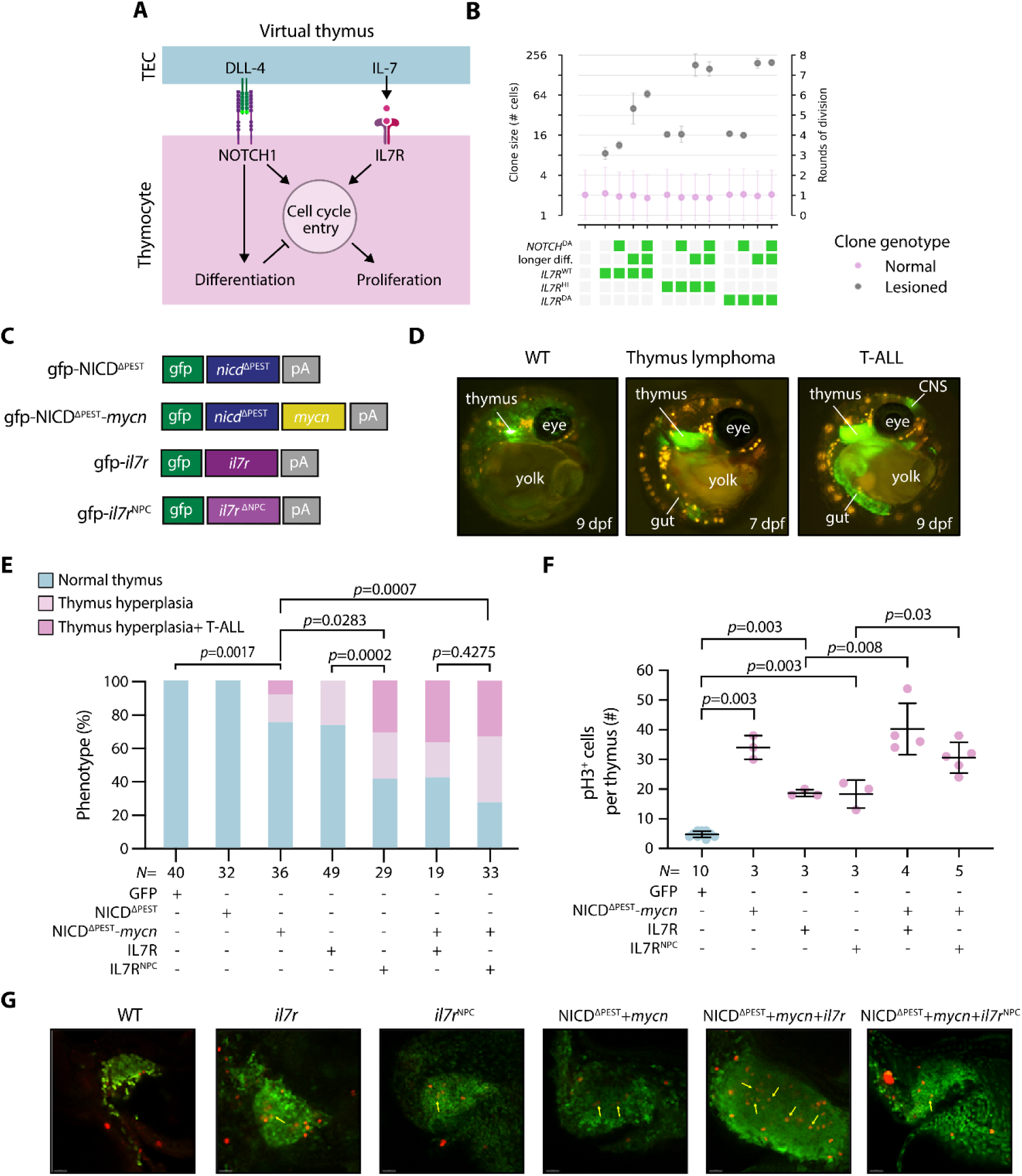
Interplay of IL7R and NOTCH1 potentiates clonal expansion *in vivo*. (A) Schematic of Notch1 and IL7R signaling and their regulation of thymocyte cell cycle entry, proliferation, and differentiation as implemented in the virtual thymus model. (B) Simulated data; rounds of cell division of normal and lesioned clones with different types of lesions in NOTCH1 or IL7R signaling. Note that the y-axis is base-2 logarithmic. Green boxes: Condition/lesion present. Gray boxes: Condition/lesion absent. (C) Illustration of plasmids for DNA micro-injection for transient overexpression of the different genes. Note that the expression of GFP was used to select successfully injected embryos. (D) Representative image: (left) of the WT control at 9 dpf where thymocytes are labeled in green with normal thymus size; (middle) of a thymus hyperplasia phenotype at 7 dpf; (right) same embryo developed T-ALL phenotype at 9 dpf, where infiltration in other organs was detected, namely in central nervous system (CNS) and gut. (E) Percentage of phenotypes: normal thymus, thymus hyperplasia and T-ALL at 11dpf. For each construct injection, we did at least three technical replicates. For detailed statistics see Figure 5 - Supplement Table 1. Statistical data rounded to 4 decimal places. (F) Number of pH3 positive cells per thymus counted during confocal imaging. (G) Representative confocal images of immunostaining against pH3 (red) and GFP (green). Scale bar is 20µm. *N* represents the number of biological samples.

To test the outcome of the virtual model, we performed a series of *in vivo* experiments to induce NOTCH1 and IL7R signaling in thymocytes. To constitutively activate NOTCH1 signaling in thymocytes, we took advantage of the previously cloned medaka *notch1b* intracellular domain (NICD) construct (Aghaallaei et al. 2021; 2022), which was driven by a thymocyte-specific promoter (Bajoghli et al. 2015). Furthermore, the destabilizing PEST domain from the NICD was removed in this construct (hereafter referred to as NICD^ΔPEST^; Figure 5C). This genetic modification was made due to the known impact of the PEST domain on protein stability, as nonsense mutations lacking the PEST domain of the human *NOTCH1* gene have been frequently observed in T-ALL patients (Breit et al. 2006; Ferrando 2009; Weng et al. 2004) (Figure 5 - Supplement 1A). To mimic the *in silico* IL7R^HI^ clones, we employed a previously developed construct wherein a thymocyte-specific promoter drives the medaka full length *il7r* cDNA (Aghaallaei et al. 2021). We then performed mutation in the extracellular juxtamembrane-transmembrane region of the medaka *il7r* cDNA (Figure 5 - Supplement 1D) to develop a dominant active IL7R form, akin to the NPC mutation found in the *IL7R* gene in some T-ALL patients (Zenatti et al. 2011; Oliveira et al. 2022), hereafter called *il7r*^NPC^. In our experimental setup, the promoter also co-expressed GFP, which allowed us to (i) identify thymocytes expressing the oncogenes, (ii) determine the clonal expansion of cells expressing the oncogene, and (iii) assess thymus hyperplasia and infiltration into other organs, a characteristic feature for T-ALL. DNA constructs were then injected into blastomeres of embryos at one-cell stage and they were observed during their development using live imaging, with a focus on two specific phenotypes: Firstly, we assessed whether the thymus exhibited enlargement beyond the normal size for its developmental stage. In particular, thymus lymphoma was defined as hyperplasia with an increase of more than twice the organ’s typical size. Secondly, we identified T-ALL by observing infiltration of GFP-co-expressing malignant cells in other organs, including the brain, intestine, or heart (Figure 5D). To align our *in vivo* experiments with the *in silico* conditions, we limited our observations to 11 dpf, concluding the experiments at the freshly hatched yolk-sac larval stage. Since thymopoiesis begins at 3 dpf in medaka embryos (Bajoghli et al. 2015; 2009), this means we monitored thymus growth for a period of 8 days.

None of the embryos injected with the NICD^ΔPEST^ construct (N=32) displayed thymus hyperplasia at 11 dpf (Figure 5E), supporting our simulation outcome showing that global activation of NOTCH1 signaling in thymocytes alone is insufficient to result in clonal expansion within a short time frame of 8 days. The MYC oncogene is an endogenous downstream target of NOTCH1 and plays a major role in NOTCH1-induced transformation (Sanchez-Martin and Ferrando 2017). Therefore, to attempt to shift the balance of NOTCH1 action towards proliferation, we decided to introduce the medaka *mycn* cDNA into our construct. Consequently, we observed that 25% of injected embryos with NICD^ΔPEST^ and *mycn* (N=36) displayed thymus hyperplasia at 11 dpf (Figure 5E). Among them, 8% also exhibited a massive infiltration of GFP-expressing cells in other organs such as the brain and gut. Further analysis of sorted GFP-expressing thymocytes revealed ectopic *il7* expression, while the expression level of endogenous *il7r* remained unchanged (Figure 5 - Supplement 1B). Whole-mount in situ hybridization (WISH) analysis further confirmed a robust upregulation of *il7* expression in the thymus of these embryos (Figure 5 - Supplement 1C). MYCN is a basic helix-loop-helix transcription factor that is downstream of several pro-proliferative signaling pathways (Ruiz-Pérez, Henley, and Arsenian-Henriksson 2017), including NOTCH1 in the context of thymocytes (Sanchez-Martin and Ferrando 2017)). Together with our experimental results, it is therefore likely that MYCN could enact the proliferative effect of NOTCH1, shifting the balance in the incoherent feed-forward loop towards proliferation. Interestingly, our results also indicate that transformed thymocytes acquire the ability to release IL- 7 (Figure 5 - Supplement 1B, C), which may further promote proliferation in an autocrine fashion.

To test whether constitutive IL7R activation could enhance the development of thymus hyperplasia and T-ALL, we injected various constructs to overexpress either *il7r* or the dominant active *il7r*^NPC^ alone, or in combination with NICD^ΔPEST^ and *mycn*. Thymus hyperplasia was found in a slightly higher frequency of 26% (N=49) or 59% (N=29) of embryos, respectively, when only *il7r* or *il7r*^NPC^ was overexpressed in thymocytes. Additionally, we FACS-sorted thymocytes and performed a qPCR for lck, since this gene was shown to be upregulated in zebrafish after IL7R^DA^ T-ALL development using RNASeq data (Oliveira et al. 2022). Indeed, we detected a significant upregulation of *lck* upon *il7r*^NPC^ overexpression when compared to the WT counterparts. However, upregulation was not significantly different when the WT was compared to *il7r*^HI^ condition (Figure 5 - Supplement 1D). Combining *il7r*^NPC^ and NICD^ΔPEST^ and *mycn* yielded a notably elevated frequency of ∼73% (N=33) for thymus hyperplasia (Figure 5E). Of this group, 33% also exhibited a T-ALL phenotype, suggesting an additive effect of these factors in the T-ALL development (Figure 5 - Supplement Table 1). GFP and pH3 double staining of embryos exhibiting the T-ALL phenotype further confirmed that the combination of NICD^ΔPEST^, *mycn* and *il7r* overexpression in thymocytes resulted in a higher number of mitotically active cells within the thymus (Figure 5F). Notably, we observed many pH3-stained cells in the inner zone of the medaka thymus (Figure 5G, arrows), an area where WT thymocytes are mitotically quiescent (Bajoghli et al. 2015; Aghaallaei et al. 2021).

The prediction that higher NOTCH1 signaling does not lead to increased proliferation is likely due to the incoherent feed-forward loop downstream of NOTCH1, which shuts down excessive proliferation by triggering differentiation to a non-dividing cell fate (Figure 5B). We observed a similar effect in our *in vivo* experiments, where activation of NOTCH1 alone did not lead to thymus hyperplasia within the observed time window of 8 days (Figure 5E). In contrast, combining NOTCH1 with its downstream pro-proliferative effector *mycn* led to thymus hyperplasia. Intriguingly, in other *in vivo* systems, long-term activation of NOTCH1 signaling in thymocytes is one of the main drivers of T cell leukemogenesis and T-ALL development (Weng et al. 2004; Lin et al. 2006; Chen et al. 2007; Liu et al. 2017; Neumann et al. 2014). Several *in vivo* studies have demonstrated that constitutive activation of NOTCH1 receptor leads to leukemia development after several weeks in zebrafish (Chen et al. 2007; Blackburn et al. 2012) or months in mice (Chiang et al. 2008; Hu et al. 2009; Sharma et al. 2006; Wendorff and Ferrando 2020). We did not observe thymus hyperplasia when upregulating NOTCH1 signaling alone, which may stem from the shorter time window of our *in vivo* experiments (i.e., 8 days). However, in the virtual thymus model, a sole upregulation of NOTCH1 cannot produce thymus hyperplasia regardless of the duration of pathway activation. Different possibilities could reconcile these discrepancies. Firstly, the balance between pro-proliferative and pro-differentiation effects of NOTCH1 *in vivo* could be different from what we modeled. Secondly, there could be additional negative feedbacks between proliferation and differentiation, which our model does not consider. Thirdly, proliferation *in vivo* could continue even past differentiation if NOTCH1 signaling is sufficiently stimulated. Finally, long-term pathway misregulation could act as a stressor that promotes additional compounding lesions.

Overall, our *in vivo* results support the outcomes of our simulations, showing that overexpression of IL7R alone, but not NOTCH1 alone, is sufficient to induce thymus hyperplasia in a very short time period. However, dual activation of IL7R and the NOTCH1-MYCN axis can promote clonal expansion *in vivo*, leading to the rapid development of thymus hyperplasia and T-ALL in the medaka model system within eight days.

### Oversupply of IL-7 by the thymic niche could accelerate the T-ALL development

One of the most unexpected outcomes of our simulations was that a defective TEC network provides lesioned clones with a substantial advantage in proliferation. Given the crucial role of IL-7 in proliferation of thymocytes, and considering that TECs, and not thymocytes, serve as the primary source of IL-7 cytokine in the normal thymic milieu, we next asked whether elevated *il7* expression in TECs alone is sufficient to induce thymus hyperplasia and to enhance T-ALL development in a thymus where thymocytes express high levels of *il7r*. This question was also driven by the results of scenario 27, which showed a twofold increase in thymocyte population size when all TECs express IL-7 (Figure 2E-G). In this thymic niche enriched with IL-7, lesioned clones either with IL-7^HI^ or extended differentiation times showed a significant advantage in clonal expansion compared to their WT counterparts (Figure 6A). Further, the impact of alterations in the thymic niche on T-ALL development has not been previously addressed. Therefore, we monitored the thymus development of the TEC:IL-7^HI^ embryos until 11 dpf to mimic scenario 27 *in vivo*. We found that 12% of these embryos (N=24) displayed signs of thymus hyperplasia (Figure 6B), however, we did not observe massive infiltration of GFP-expressing cells into other organs. The frequency of thymus hyperplasia was increased to 41% (N=32) when DNA constructs designed to overexpress *il7* in TECs were co-injected with the construct overexpressing *il7r* in thymocytes (Figure 6B). Comparably, thymus hyperplasia was only observed in 26% of embryos injected with the IL7R overexpression construct in thymocytes alone (N=49). A synergistic effect was observed when *il7* was overexpressed in TEC along with NICD and *mycn* in thymocytes (Figure 6B). Interestingly, the frequency of embryos with T-ALL development was increased to 25% (N=48) in this group. Taken together, our *in vivo* results reveal that an excess of IL-7 in the thymic environment, combined with alterations that affect the differentiation status of thymocytes, triggers the T-ALL development in a short period of time.

**Figure 6.**
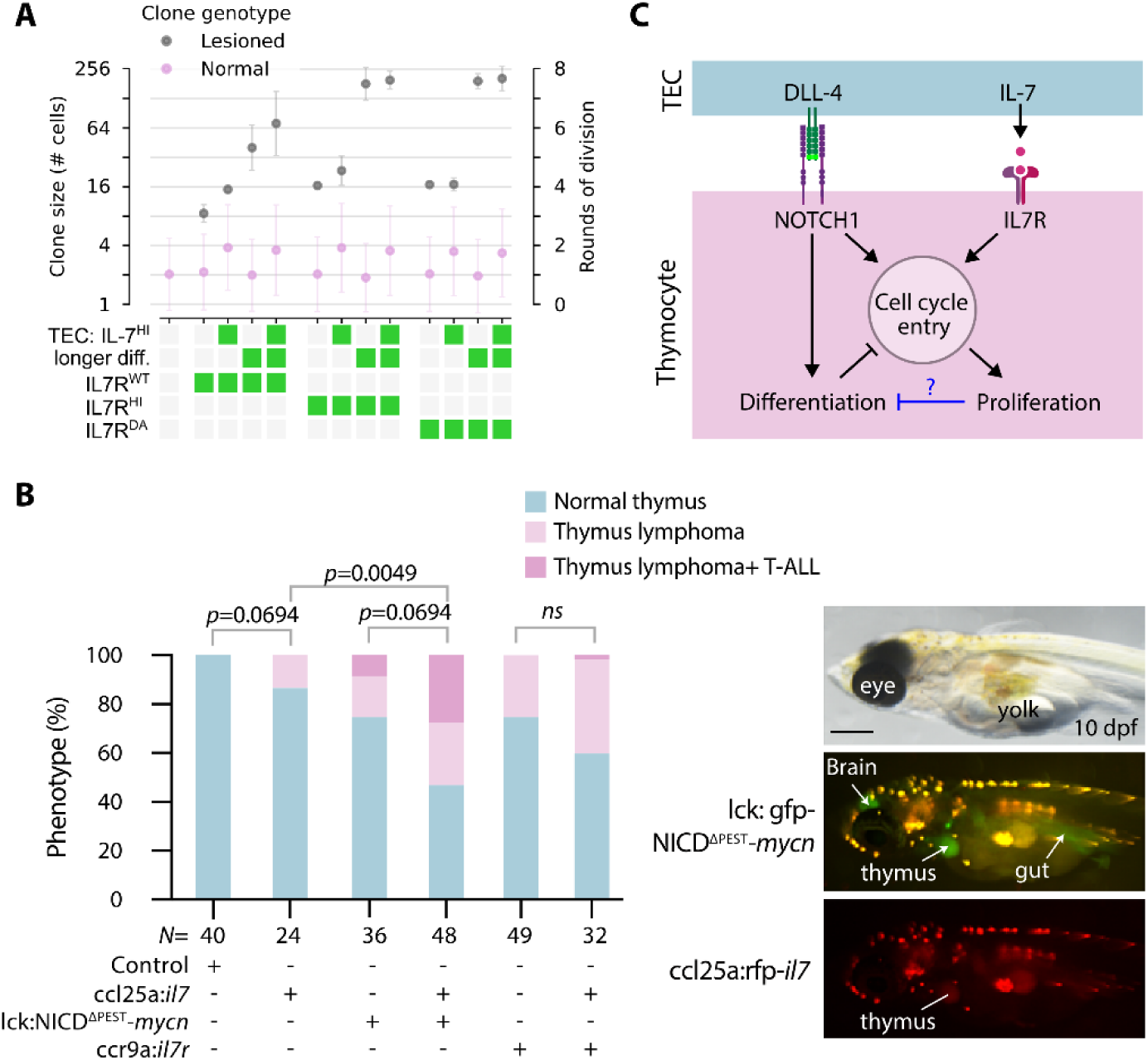
IL-7 supplied by the niche can lead to thymus hyperplasia. (A) Simulated data; rounds of cell division of normal and lesioned clones for different combinations of lesions and TECs expressing IL-7 ubiquitously. Note that some conditions are reproduced from Figure 5B for easier comparison. (B) Left panel: Frequency of thymus hyperplasia and T-ALL phenotype observed in the injected embryos at 11 dpf. For detailed statistics see Figure 5 - Supplement Table 1. Statistical data rounded to 4 decimal places. Note that some data from Figure 5E are repeated here to facilitate a better comparison. *N* represents the number of biological samples. Right panel: representative images at 10 dpf after co-injection of lck:gfp-NICD^ΔPEST^-*mycn* and ccl25a:tagRFP-il-7 constructs. Arrows indicate GFP-expressing malignant thymocytes in the thymus, brain, and gut. Scale bar is 400 µm (C) We propose a hypothetical negative feedback loop from proliferation to differentiation, which could explain discrepancy between *in silico* predictions and *in vivo* observations.

Overall, several of our experimental observations indicate that there might be additional negative feedback from pro-proliferative signaling to differentiation (Figure 6C). Most strikingly, manipulations in IL7R or in thymic IL-7 levels are sufficient to induce hyperplasia and T-ALL (Figure 5E and Figure 6B) – an outcome that is much more severe than the model predicts (Figure 5B and Figure 6A). In simulations, we could only obtain such massive overproliferation by modifying multiple parameters, including delaying differentiation, accelerating cell divisions, and reducing TEC density (Figure 4 – Supplement 5), as differentiation in our model is a hard stop to proliferation regardless of pro-proliferative stimuli. Similarly, the experimental observation that NOTCH1-MYCN upregulation leads to thymus hyperplasia and T-ALL development also suggests that proliferation can proceed unbridled by differentiation *in vivo* when sufficiently stimulated. A negative feedback loop from pathways regulating proliferation (or cell cycle entry) on differentiation pathways could explain these discrepancies between the virtual thymus model and *in vivo* observations. We therefore propose that continued thymocyte proliferation can downregulate differentiation pathways and maintain cells in a constant proliferative state.

## CONCLUSION

This study demonstrates the powerful synergy between computational modeling and *in vivo* experimental validation in dissecting the complex cellular and molecular interactions within an organ that drive disease initiation and progression. Further, we highlight a previously underexplored area – the role of TECs and the thymic niche in the development of T-ALL – thereby bridging a major gap in our understanding of the earliest stages of disease development.

By simulating over 1500 scenarios, we identified potential drivers such as alterations in the shape, density, and spatial organization of the TEC network – factors that are inherently difficult to manipulate in experimental systems. Consistent with these computational predictions, our *in vivo* results reveal that an enriched thymic IL-7 milieu can accelerate disease progression, particularly when thymocytes display enhanced IL7R levels or express constitutively active NOTCH1 and MYCN overexpression. These results challenge the traditional gene-centric paradigm of T-ALL and emphasize the critical influence of the spatial structure of the thymic microenvironment.

However, it is worth noting several limitations of our current virtual thymus model. The cell-based computational framework was calibrated to mimic the medaka embryonic thymus (Bajoghli et al. 2015; 2009; Aghaallaei et al. 2021), and as such, it may not fully capture the cellular dynamics or thymic architecture of other species, particularly human. On the technical side, the simulation lacks the ability to represent changes in TEC shape over time and TEC proliferation, as all particles are treated as loose objects and hence movement of a TEC would lead to detachment of its protrusion particles. This limitation could be overcome with a more advanced biomechanical framework that includes bonded interactions between particles such as Tissue Forge [Sego et al. 2023], which would ensure that particles belonging to the same cell stay together as one part of the cell moves. Due to this limitation, the dynamic remodeling of the thymic niche is not fully accounted for in our current model. Furthermore, the model simplifies certain T-cell developmental stages as it does not consider cross-regulatory feedback loops between IL-7 and Notch pathways, and does not include thymic growth over time. These simplifications are due to the lack of species-specific knowledge and the tools to investigate these questions in medaka. Nonetheless, we anticipate that adapting the framework for use in murine or human thymic contexts, where stage-specific markers and niche components are well characterized and where detailed morphometric studies of TEC architecture are emerging [Lagou et al. 2024], would enable us to explore these questions.

Despite its current simplifications, our integrative approach offers valuable insights into how microenvironmental factors influence leukemia expansion and provides a framework for exploring their role in chemoresistance and relapse. Ultimately, these findings open new avenues for developing therapeutic strategies aimed at disrupting the leukemic- supportive niche, with the potential to improve treatment outcomes in T- ALL.

## MATERIALS AND METHODS

### In silico model

To develop, implement, and simulate the virtual thymus model, we used the modeling and simulation software EPISIM (Sütterlin et al. 2017; 2013). This multiscale simulation software uses an agent-based paradigm to represent cells as spheres or ellipsoids in three-dimensional space, enabling each of the cells to individually perform internal processes based on flow diagrams and logic rules (e.g. if/else statements) or differential equations, and also implements a partial differential equation solver to simulate diffusion of chemicals in the extracellular space. The virtual thymus model has been comprehensively described in the supplementary material to our previous study (Aghaallaei et al. 2021). In this work, we used this model as a baseline to introduce a small number of additions, explained in the following. A summary of the full model implementation is provided in the Appendix.

#### Rate of IL-7 pathway signal transduction

The equation describing the IL-7 signaling rate in each cell was rescaled to introduce the parameter *a*_IL-7_ for the IL-7 signal transduction activation rate. The default value of *a*_IL-7_ was chosen such that the model behavior did not change. The rescaled equation for IL-7 signal transduction reads:

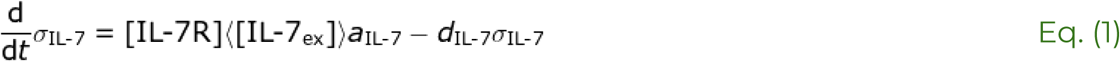

where *σ*_IL-7_ is the IL-7 pathway signal transduction activity, [IL7R] is the IL7R concentration in the given cell,〈[IL-7_ex_]〉is the mean extracellular IL-7 concentration in the cell’s microenvironment, and *d*_IL-7_ is the signal transduction deactivation rate.

In the scenario where we introduced a lesioned clone with dominant-active IL7R, IL-7 signal transduction activity was calculated as

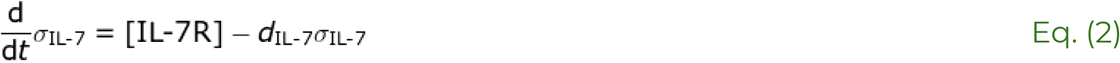

Thus, in IL7R^DA^ clones, neither extracellular IL-7 nor the signal transduction activation rate *a*_IL-7_ had an impact on cells’ IL-7 signaling pathway activity.

#### Depletion of extracellular IL-7

In our original model (Aghaallaei et al. 2021), IL-7 ligand is released back to the extracellular space after binding to IL7R. Thus, thymocytes do not affect the extracellular IL-7 gradient, which is given by the partial differential equation

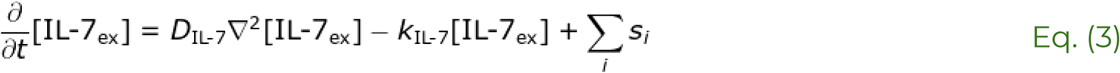

where *D*_IL-7_ is the IL-7 diffusion constant, *k*_IL-7_ is a baseline level of extracellular IL-7 degradation, and *si* is a source term accounting for IL-7 secretion, e.g. by a subset of TECs. In the modeling platform we use, secretion occurs throughout a cell’s volume; more specifically, for solving the partial differential equation, space is discretized into voxels, and all voxels *i* that intersect with the ellipsoids that are used to represent a cell contribute to the source term.

In this work, we introduce the option, controlled via a Boolean flag *DEPL* that is set by the user before the simulation starts, to simulate a scenario where thymocytes internalize IL-7 ligand after it is bound to IL7R. In this scenario, the extracellular IL-7 concentration is given by:

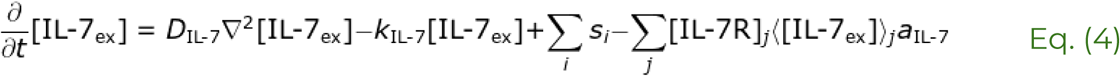

Note that the additional sink term in Equation 4 represents extracellular IL- 7 depletion and is simply the IL-7 signal transduction activity from Equation 1 summed over all voxels *j* that intersect with thymocytes. We assume that all internalized IL-7 is permanently removed from the extracellular pool.

For lesioned IL7R^DA^ clones, we used the alternative sink term

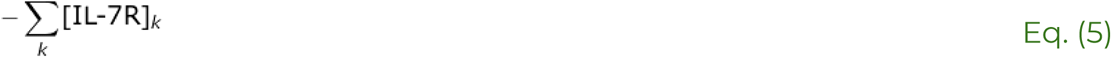

which is summed over all voxels *k* that intersect with thymocytes of the lesioned clone. This term corresponds to the IL-7 signal transduction activity as shown in Equation 2. Reduction of extracellular IL-7 below zero was prevented by a computational check.

#### Determining clonal lineages

The EPISIM software we used assigns to each cell a unique cell identifier. We made use of this unique cell identifier to generate unique clonal identifiers as follows. For each thymocyte that entered the simulation via migration from outside the thymus, we used the value of their unique cell identifier to assign the value of a new clonal identifier variable. This cell-specific value of the clonal identifier variable was passed on to any daughter cells produced via cell division. Thus, using the clonal identifier, we essentially generated an *in silico* clonal label which we used to determine clonally related cells.

#### Lesioned clone

To generate a lesioned clone in a given simulation, we created a computational label to mark a single cell selected randomly among new thymic immigrants once the thymic cell population reached its homeostatic size (t >= 60h). This label marked this cell and its progeny as a “lesioned clone”, such that we could specifically modulate parameters and introduce new rules only for this single clonal lineage. We tested the effect of several alterations specific to the lesioned clone (Figure 4 - Supplement Table 1).

#### Parameter variations

Please refer to the supplementary material of (Aghaallaei et al. 2021) for a comprehensive explanation of all parameters and the choice of the reference values. Altered parameters used in this work are listed in Figure 1 - Supplement Table 1.

### Calculation of simulated clone size

Clone size was defined as the number of terminal leaves in a cell lineage tree (see scheme in Figure 4C); terminal leaves were counted regardless of the ultimate fate of those cells (i.e. positive or negative selection). All clones of the same genotype (wildtype or lesioned) belonging to the same simulated scenario were averaged across all replicate simulations of that scenario. This calculation gives the average number of cells per clone in a given scenario. The number of cells was log2-transformed to represent rounds of cell division.

### In vivo model

Medaka (*Oryzias latipes*) husbandry was performed in accordance with the German animal welfare standards (Tierschutzgesetz §11, Abs. 1, Nr. 1, husbandry permit no. 35/9185.46/Uni TÜ.) The transgenic line (tg) lck:gfp was described previously (Bajoghli et al. 2015). All experiments conducted in medaka embryos were performed prior to the legal onset of animal life stages under protection, utilizing both males and females of the wildtype line Cab (Loosli et al. 2000).

### Cloning of DNA constructs

To overexpress *il7* in TECs, we used the construct ccl25a:*il7*, which was described previously (Aghaallaei et al. 2021). To overexpress *il7r* in thymocytes, the full-length medaka *il7r* cDNA (Aghaallaei et al. 2021) was cloned into vectors containing a medaka thymocyte-specific promoter (Bajoghli et al. 2015) that drives a fluorescent protein (sfGFP or TagRFP). To generate a dominant active form of *il7r*, we introduced three amino acids, Asparagine (N), Proline (P), and Cysteine (C) (known as NPC mutation, as shown in (Zenatti et al. 2011) and (Oliveira et al. 2022)), in the medaka *il7r* extracellular juxtamembrane-transmembrane interface region after position 266 using site-directed mutagenesis (Figure 5 - Supplement Figure 1D), this corresponds to the human NPC mutation in the position 242 of *IL7R* gene. To activate Notch signaling in thymocytes, we utilized the medaka *notch1b* intracellular domain (NICD), as described previously (Aghaallaei et al. 2021), and removed its PEST domain to enhance protein stability, mimicking the situation reported in most T-ALL patients harboring gain-of-function mutations in the NOTCH1 gene (Weng et al. 2004). For overexpressing the *mycn* oncogene, we first isolated the full-length medaka *mycn* cDNA (accession number: ENSORLG00000022362) and introduced a mutation at position 44 from Proline to Leucine (P44L) using site-directed mutagenesis, as identified in T-ALL patients (Liu et al. 2017). The DNA fragments were then cloned into vectors containing a medaka thymocyte-specific promoter, either alone or in combination (Bajoghli et al. 2015).

### DNA micro-injection

Plasmids at concentration 10-25 ng/µl together with I-*Sce*I meganuclease and NEB buffer (NewEngland BioLabs), were co-injected into the blastomere at one-cell stage embryos. Fluorescent signals, based on the constructs either sfGFP, tagRFP or mTurquoise, were used to select positive embryos.

### Generation and genotyping of medaka *il7* mutant

The CRISPR-Cas9 approach was employed to generate medaka *il7* crispant. Crispr RNA (crRNA AGTAGACTGATGCAAAGAAG) was designed using the CCTop website (Stemmer et al. 2015; 2017) and ordered from IDT. RNP complexes were prepared as described before (Hoshijima et al. 2019) using AltR tracrRNA (IDT) and Alt-R S.p. Cas9 nuclease (IDT). The injection mixture contained a total of 25 µM crRNA:tracrRNA duplex and 25 µM Cas9. Injection was performed into the blastomere at the one-cell stage transgenic embryos carrying the lck:gfp reporter. Injected embryos were raised until 8 dpf and then fixed in 4% PFA/2xPBS + 0.1% Tween20 for whole-mount immunostaining. To correlate the phenotype with the genotype, each embryo was genotyped using PCR and Sanger sequencing

### Phenotype assessment

The development of injected embryos was monitored using a NIKON SMZ18 stereo-fluorescent microscope. Only embryos with fluorescent signals in the thymus region were selected for further analysis. Thymuses of embryos before hatching or freshly hatched yolk sac larvae were examined. Thymuses with at least a two-fold increase in size compared to transgenic embryos carrying the lck:gfp reporter construct (Bajoghli et al. 2015) were considered as having thymic hyperplasia. The identification of cells expressing fluorescent proteins and oncogenes outside the thymus, such as in the brain, gut, and heart, was considered indicative of the T-ALL phenotype.

### Whole-mount immunostaining

Immunostaining procedures followed established protocols (Inoue and Wittbrodt 2011; Aghaallaei et al. 2021). Briefly, mitotically active cells were identified using a rabbit anti-phosphohistone-3 antibody (Ser10, Millipore 06–570, 1:500 dilution), with a Cy3-donkey anti-rabbit immunoglobulin G secondary antibody (the Jackson laboratories, 711-165-152; 1:500 dilution). GFP expression was detected using a goat Anti-GFP antibody (Abcam, ab5450, 1:500 dilution), with Alexa488-donkey anti-goat IgG (Abcam, ab15129, 1:600 dilution) as the secondary antibody. To quantify the number of pH3-expressing cells, the entire thymus region was imaged using a LSM710 (Zeiss) confocal microscope with z-stacks (z=1 µm).

### Whole-mount *in situ* hybridization (WISH)

Whole-mount *in situ* hybridization in medaka embryos was carried out using digoxigenin-labeled RNA probes, as previously described (Aghaallaei et al. 2005). Probes used in this study for *il7r* and *il7* were previously described (Aghaallaei et al. 2021).

### Cell sorting and quantitative RT-PCR

Injected embryos were first smashed through a 40 µM strainer (Greiner) and cells were collected in 0.9x PBS with 500 U.I. Heparinum Natricum (Liquemin). Sorting was then performed using the Cell Sorter MA900 (Sony Biotechnology). WT embryos were used as negative sorting control. Only GFP positive cells were collected and used for RNA isolation. RNA of sorted cells was isolated by using NucleoSpin RNA XS (Macherey-Nagel) and 20 ng carrier RNA, following the manufacturer’s protocol. RNA was treated with rDNase and eluted in 14 µl dAqua. The first-strand cDNA synthesis was carried out with random hexamer primers and SuperScript III Reverse Transcriptase (Thermo Fisher Scientific) by following the manufacturer’s protocol. SYBR Green Kit (Applied Biosystems) was used for Quantitative PCR on the LightCycler 480 (Roche). The data was evaluated in Microsoft Excel using the ΔCt method and normalized to the housekeeping gene ef1a. The primers used for RT-PCR have been described previously (Aghaallaei et al. 2021).

### Statistical analysis

Statistical analysis was performed using RStudio version 2024.4.2.764 (Posit team 2024) using R version 4.4.0 (R Core Team 2024), the R library jtools (Long 2022) and GraphPad Prism version 8.0.2. The two-tailed Fisher’s exact t-test was used to compare the thymus hyperplasia phenotype with normal thymus. Welch’s t-test was used for thymus volume size comparison in WT and *il7* crispants, multiple t-test was used for comparison of normal thymus with thymus hyperplasia and T-ALL phenotype. Mann-Whitney test was used to evaluate the qPCRs. p-values <0.05 were considered as statistically significant. The number of biological samples for each experiment (*N*) is indicated in the figures, and bar graphs present the absolute numbers with mean ± SD, ± SEM or the percentage as indicated in the figure legends.

## Supporting information

Figure_1_SuppMov_1

Figure_1_SuppMov_2

Figure_4_SuppMov_1

Figure_4_SuppMov_2

## Acknowledgment

The authors thank Anna-Sophia Hellmuth for assistance with *il7* gRNA injection; the institute of Medical Virology and Microbiology for continuous support in confocal microscopy; Larissa Doll, Julia Skokowa and Karl Welte for support and encouragement. Initial computational work presented here was performed using the ALICE computer resources provided by Leiden University, later work was performed using the computer lab in the Theoretical Biology and Bioinformatics department of Utrecht University. This work was supported by the Madeleine Schickedanz Kinderkrebsstiftung (grant number D.30.28666), and Deutsche Forschungsgemeinschaft (BA 5766/5-1). ET is funded by the Dutch Research Council (NWO) in the NWO Talent Programme with project number VI.Veni.222.323. We thank members of the Theoretical Biology and Bioinformatics division for their constructive feedback on the manuscript.

We acknowledge support from the Open Access Publishing Funds of the University of Tübingen and the Faculty of Science of Utrecht University for supporting this research.

## Author contributions

E.T. designed and developed the computational model. A.M.D. performed the *in vivo* experiments and analyzed their results. B.B. designed and supervised the study. All authors jointly interpreted the data and wrote the manuscript.

## Data and materials availability

All data needed to evaluate the conclusion in the paper are present in the paper and/or the Supplementary Materials. The computational model is available on Zenodo: doi:10.5281/zenodo.11656320 (Tsingos 2024). Additional materials related to this paper such as plasmids or DNA constructs may be requested from the authors. Some graphs were created with BioRender.com.

---

**Figure 1 - Supplement 1.**
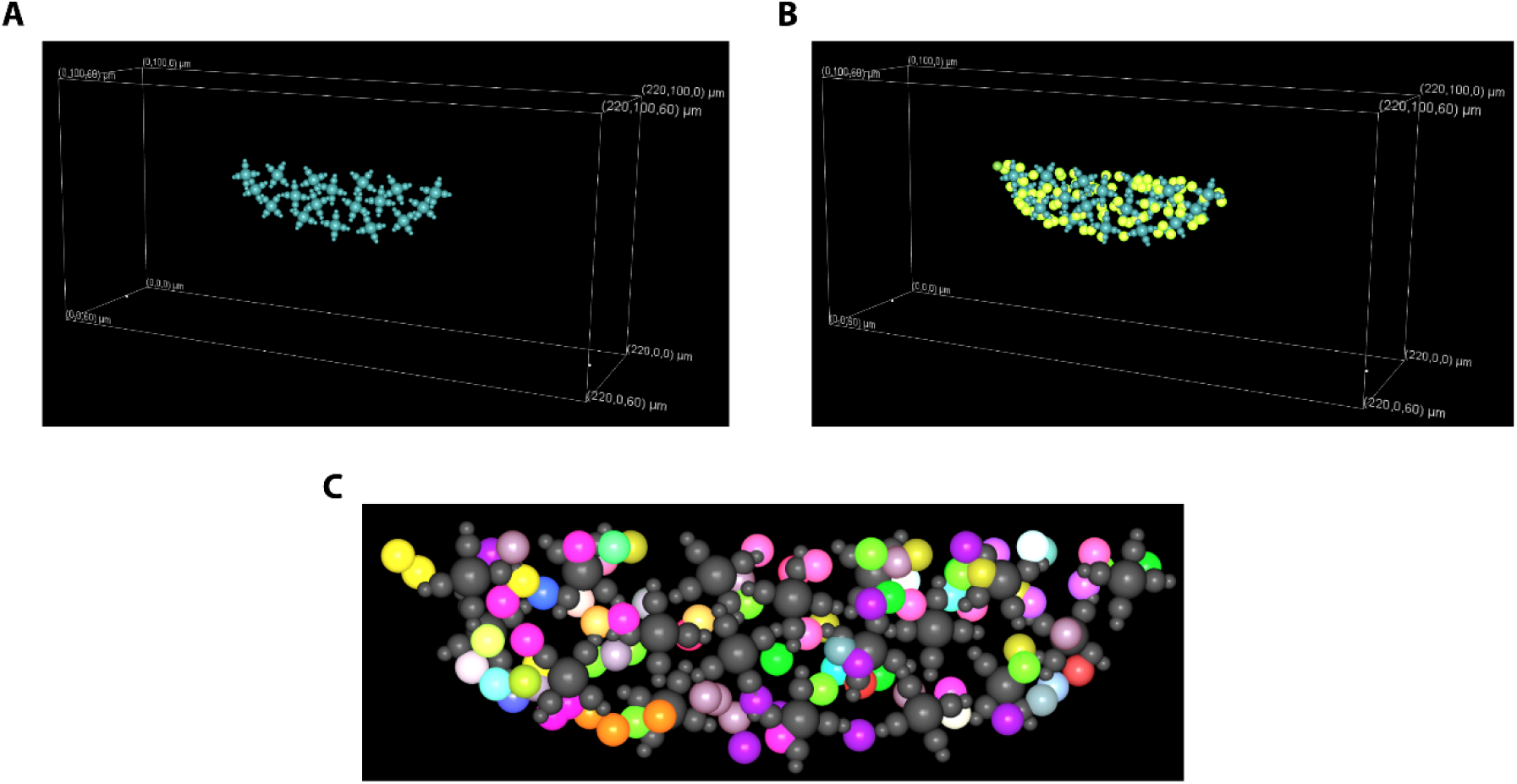
The virtual thymus represents a slice of the organ. Three-dimensional renders of the simulation. (A) Showing only TECs in cyan. (B) Showing both thymocytes (bright green) and TECs (cyan). (C) Detail showing clonal thymocyte lineages. Each lineage was assigned a randomly selected color, TECs in dark gray. In homeostasis, most clones consist of 1-8 cells.

**Figure 1 - Supplement Movie 1.**
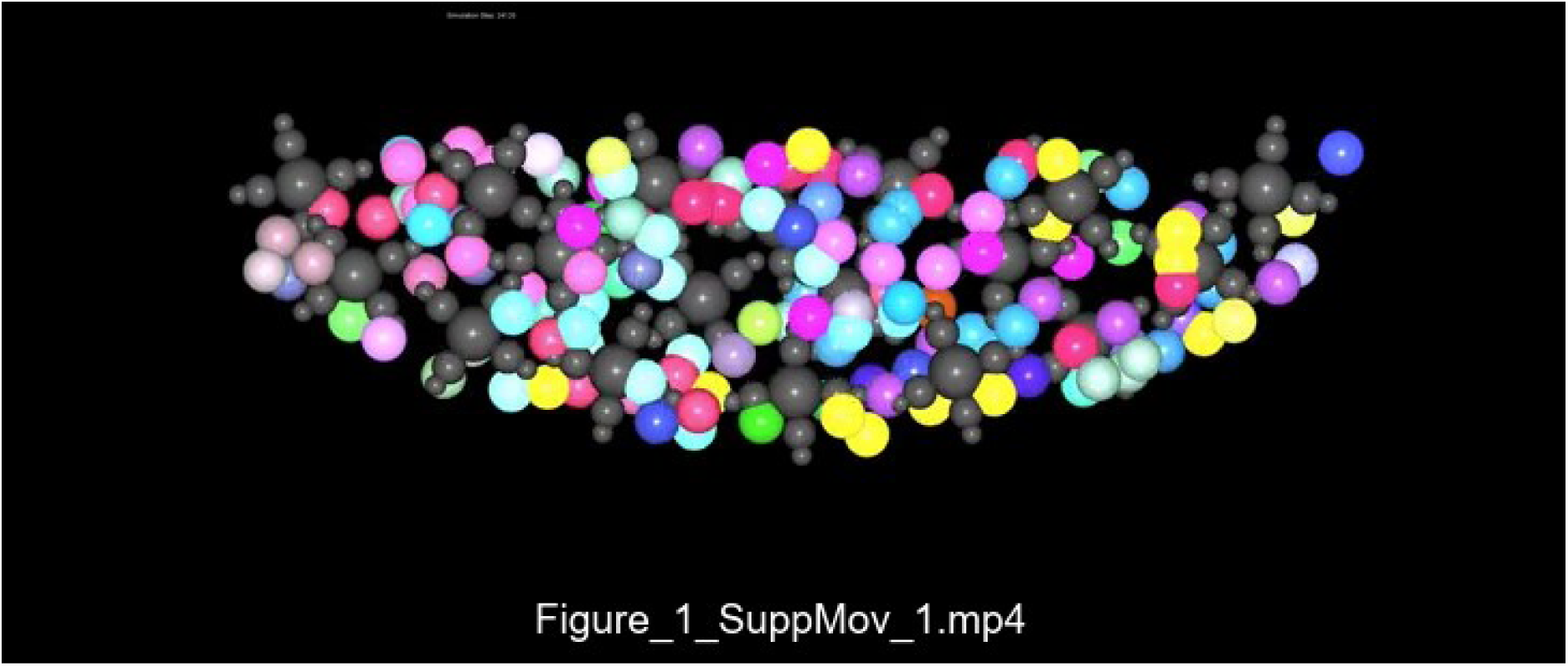
Simulation time-lapse. Three-dimensional renders of a typical simulation where each thymocyte lineage was assigned a randomly selected color. TECs are shown in dark gray. Simulation runs for 48000 steps, where 1 step is 15 seconds. Note how thymocytes can crawl over and under TECs.

**Figure 1 - Supplement Movie 2.**
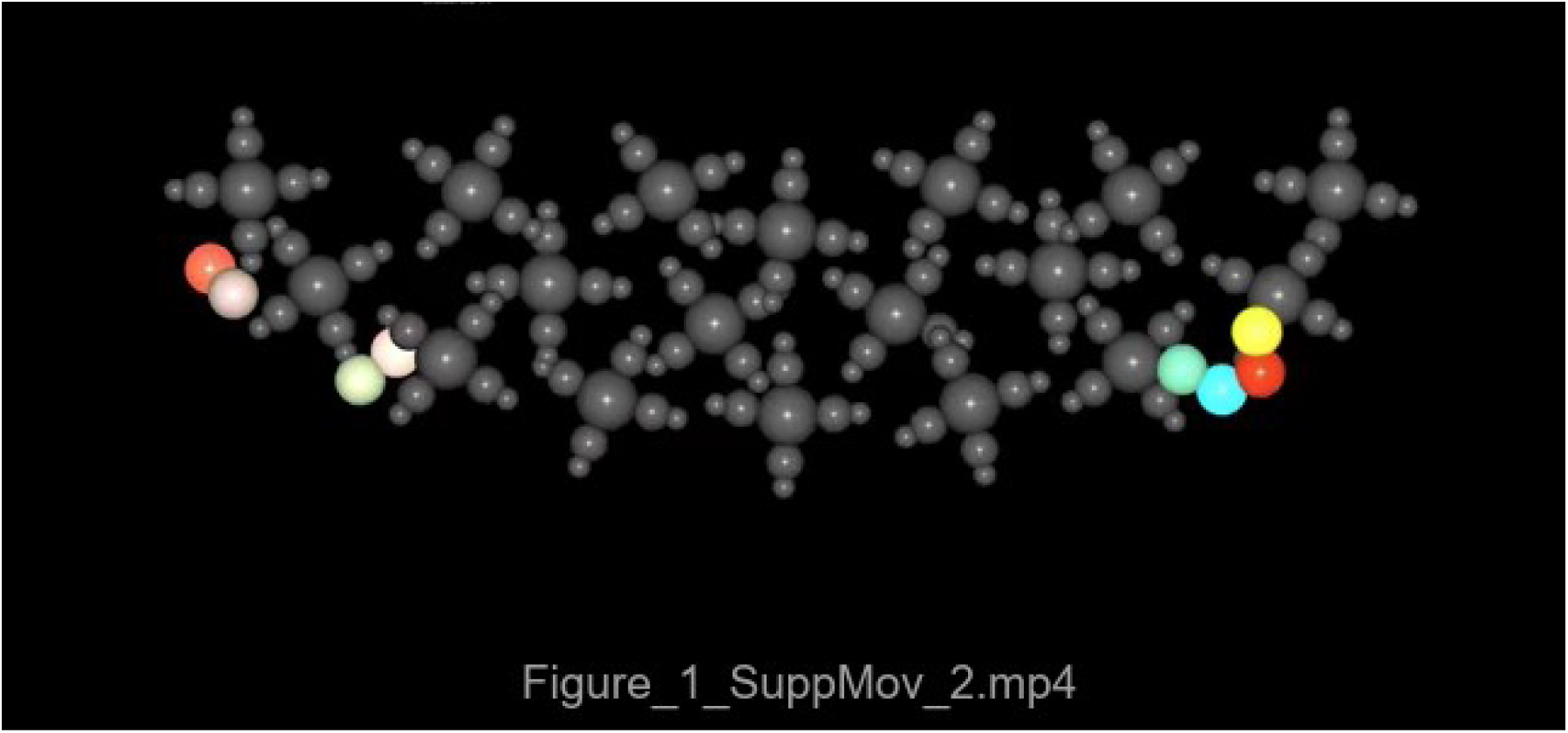
Simulation time-lapse with fewer cells to highlight clonal dynamics. In this simulation, a total of eight ETPs were introduced into the simulation at the same time to highlight clonal expansion and motility within the organ (00:10-02:00s), and later cell exit and cell death (starting at 02:08s). Three-dimensional renders of a simulation where each thymocyte lineage was assigned a randomly selected color. TECs are shown in dark gray. Each simulation step is 15 seconds.

**Figure 1 - Supplement Table 1 -.**
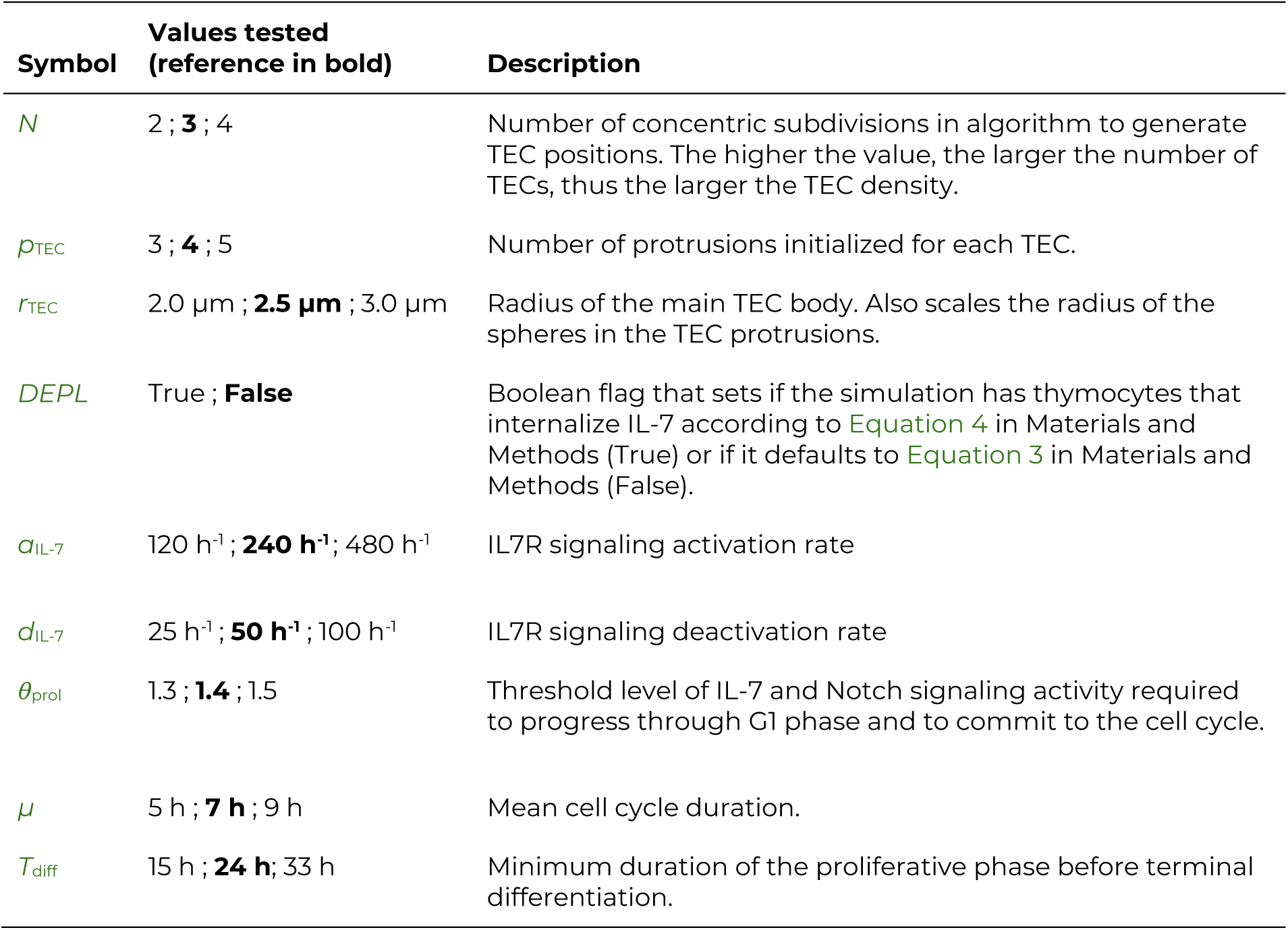
Parameters varied in this work. For a full list of parameters and their values, please refer to Aghaallaei et al. 2021.

**Figure 2 - Supplement 1.**
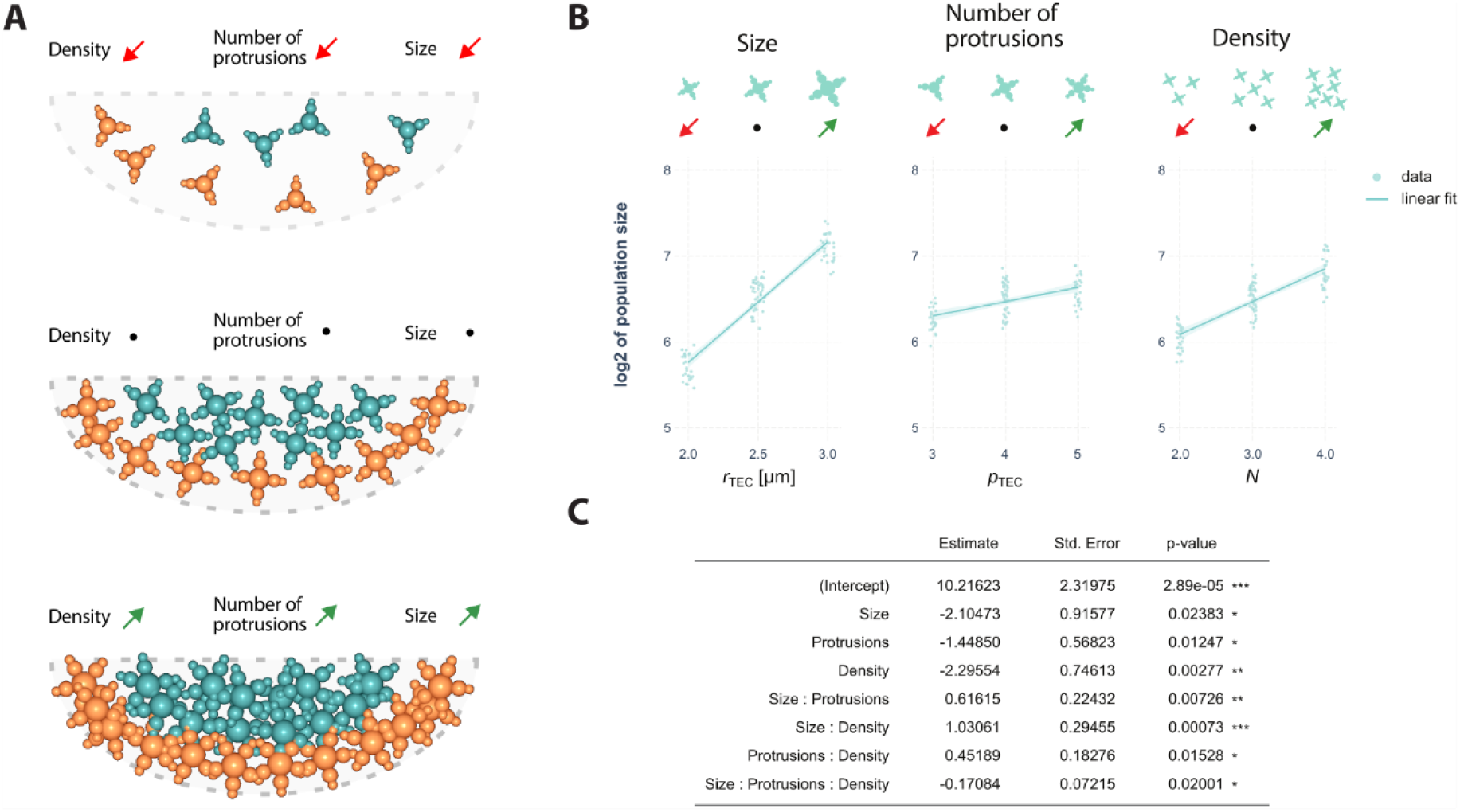
Statistical analysis of simulated data. (A) Three-dimensional rendering of TECs in the simulation with the indicated parameter alterations. Orange: IL-7-secreting TECs. Cyan: Non-secreting TECs. Dotted outline indicates thymus outline in the simulation. (B) Partial residual effect plot showing linear model fit on log2-transformed thymic population size. Shaded area is 95% confidence interval. (C) Estimated coefficients and p-values of the linear model. Significance codes: <0.001 “***”; <0.01 “**”; <0.05 “*”. All parameters have a statistically significant effect on cell number. Parameter interactions are also statistically significant, which indicates that changing combinations has a greater effect than the sum of individual changes.

**Figure 2 - Supplement 2.**
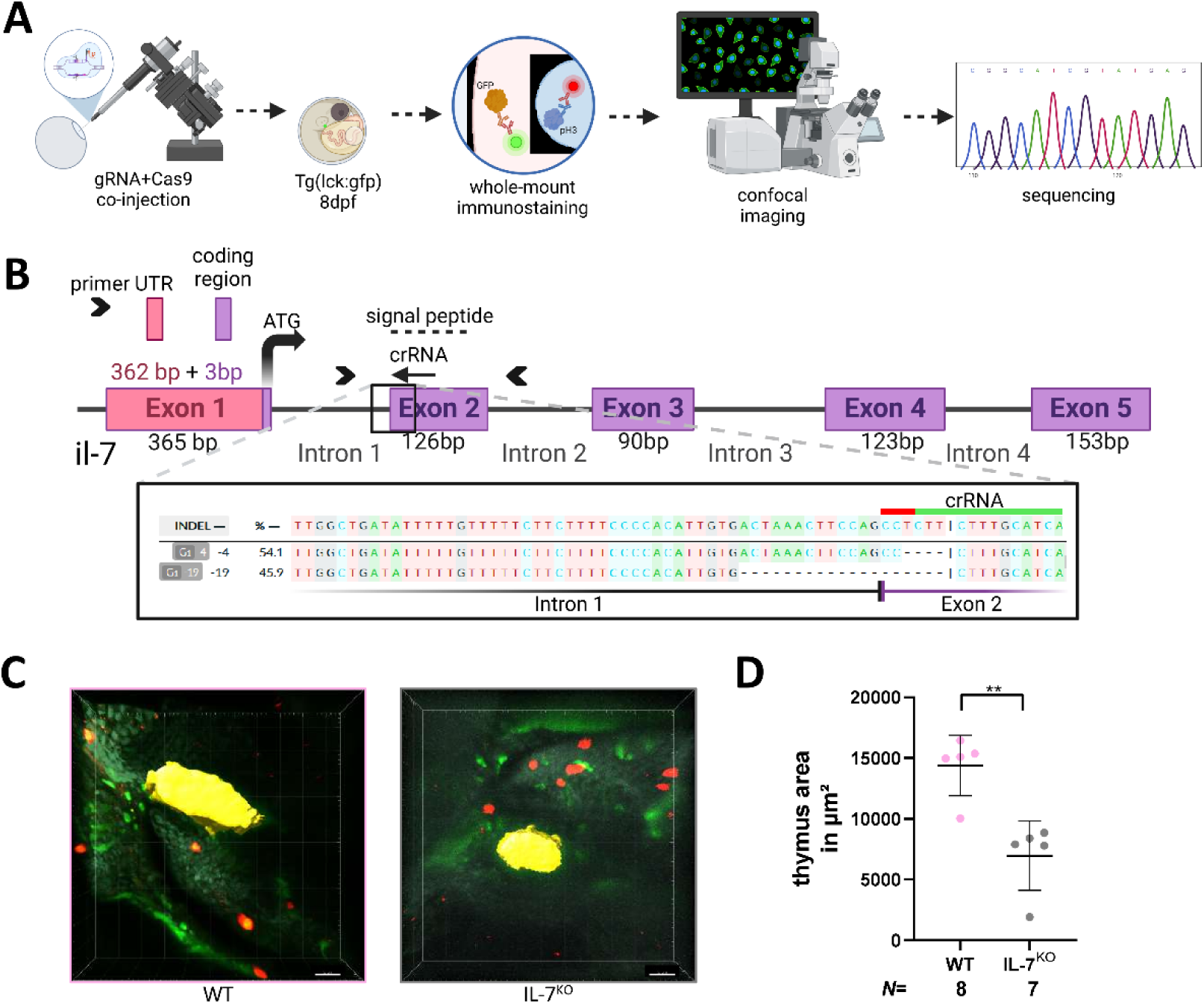
Reduced thymus volume upon IL-7^KO^. (A) Experimental design: gRNA targeting the signal peptide of il-7 was co-injected with Cas9 into the 1-cell-stage blastomere of Tg(lck:gfp) medaka. The embryo was raised until 8 dpf. Whole-mount immunostaining was performed using anti-GFP and M-phase marker phospho-histone 3 (pH3). The thymus was imaged using confocal microscopy, and embryos were genotyped by sequencing. (B) Schematic of the il-7 gene structure and binding site for the crRNA for CRISPR-Cas9 gene editing in the signal peptide region of exon 2. Zoom in: Representative sequencing data analyzed with decodr.org (Bloh et al. 2021) software. (C) Representative image of immunostaining of Tg(lck:gfp) in WT and IL-7^KO^ and evaluation of the thymus size using the area measurement tool in Imaris software (yellow). Scale bar 20 µm. (D) Quantification of thymus size: Thymus area in µm^2^ in WT and IL-7^KO^. *N* represents the number of biological samples. For statistics, unpaired t-test with Welch’s correction was used; Data show means ± standard deviation. Significance threshold was set to p= 0.05. ** means p < 0.01.

**Figure 3 - Supplement 1.**
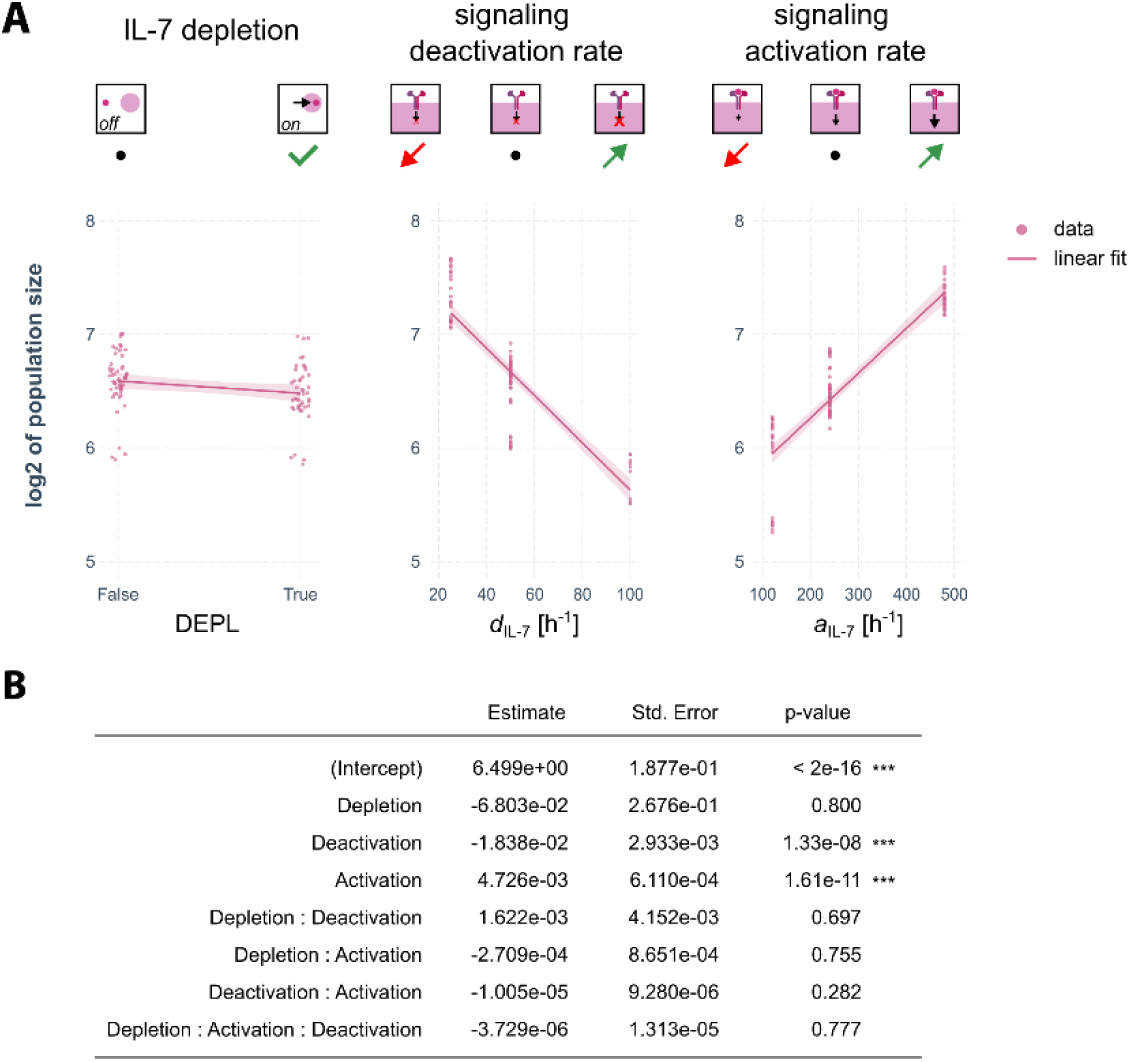
Statistical analysis of simulated data. (A) Partial residual effect plot showing linear model fit on log2-transformed thymic population size. Shaded area is 95% confidence interval. (B) Estimated coefficients and p-values of the linear model. Significance codes: <0.001 “***”; <0.01 “**”; <0.05 “*”. Although the population size is slightly reduced for IL-7 depletion, the effect is not statistically significant. In contrast, IL7R signaling activation and deactivation have a statistically significant effect. None of the interactions have statistically significant effects.

**Figure 4 - Supplement Table 1 -.** Clonal lesions and their effects.

| Lesion | Explanation |
| --- | --- |
| IL7R<sup>WT</sup> | The concentration of IL7R receptor in the lesioned clone was set to the maximum attainable in wildtype thymocytes. Thus, this lesion is representative of an extreme example among the non-lesioned population and was included as a control. |
| IL7R<sup>DA</sup> | This concentration of IL7R receptor in the lesioned clone was set to the maximum attainable in wildtype thymocytes. Moreover, the receptor was constitutively active, irrespective of the presence of IL-7 ligand. |
| IL7R<sup>HI</sup> | The concentration of IL7R receptor in the lesioned clone was set to 10-fold higher than the maximum attainable in wildtype thymocytes. |
| IL-7 autocrine | Lesioned clones secrete IL-7 at the same rate as TECs, acting as additional sources in Equation 4. |
| NOTCH1<sup>DA</sup> | Lesioned clones had constitutively active Notch signaling irrespective of the presence of Delta ligand. The levels of Notch signaling were set to the maximum attainable in non-lesioned thymocytes and could not be reduced. |
| Delayed differentiation | The lesioned clone had its rate of differentiation halved, effectively doubling the duration of the proliferative phase. |
| Slower cell speed | The lesioned clone's maximum speed was half of the non-lesioned population's maximum speed. |

**Figure 4 - Supplement 1.**
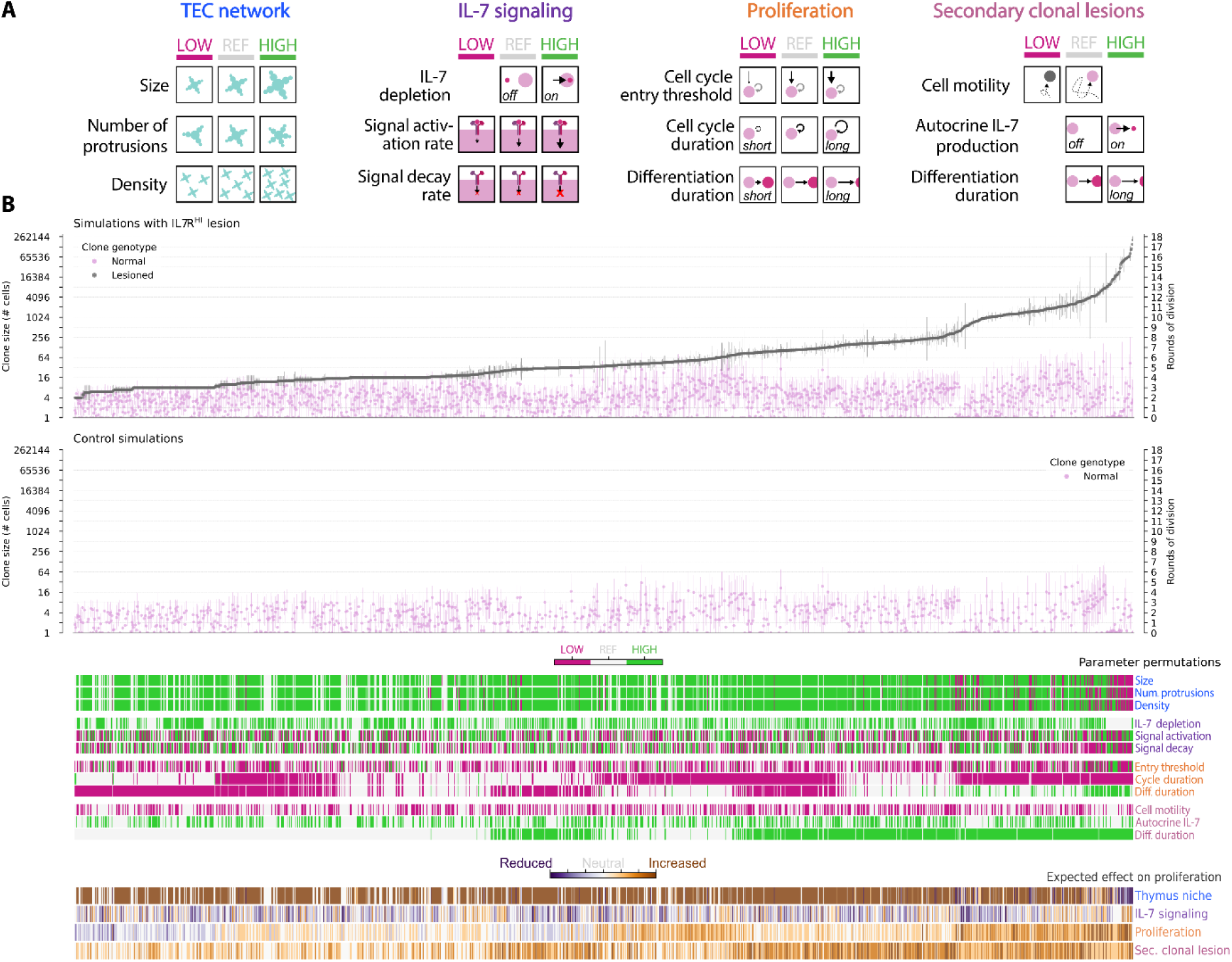
Clones in control simulations show similar sensitivity to parameter alterations as non-lesioned clones. (A) Schematic of tested scenarios shown in (B). (B) Simulated permutations ordered along increasing mean lesioned clone size for the IL7RHI condition compared to control simulations without a lesioned clone. The top panel is identical to the top panel in Figure 4 (E) and is replicated here for easier visual comparison. Expected effect on proliferation is a qualitative estimate based on preliminary simulations. A total of 1580 permutations were tested with a varying amount of replicates per permutation. Note that control simulations were run only for a subset of parameter configurations. Error bars indicate standard deviation. Note that in (B) the axis is in base-2 logarithmic scale to better illustrate rounds of cell division.

**Figure 4 – Source Data 1.**
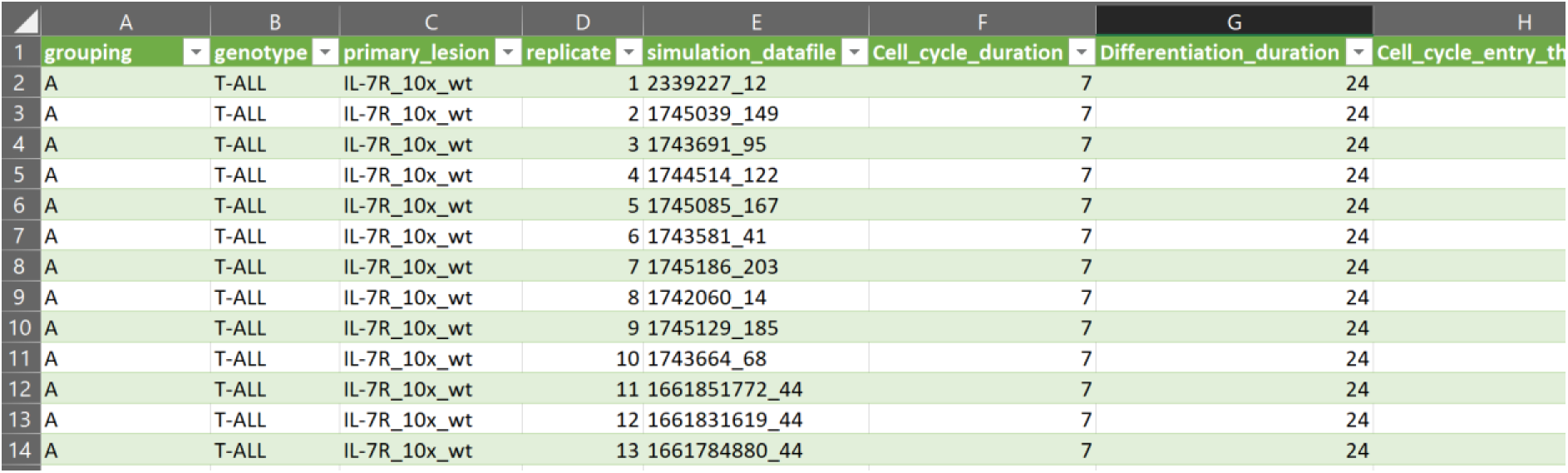
Source data file for visualization in Figure 4 E and Figure 4 Supplement 1 B. In the data file, each row corresponds to the log2-transformed average clone size for one cell genotype (normal or lesioned) from one single independent replicate simulation. Additionally, the parameter values used in that run are listed in separate columns. The “grouping” column corresponds to the simulated scenarios (parameter permutation). The “simulation_datafile” column is the unique identifier of a given simulation run.

**Figure 4 - Supplement Movie 1.**
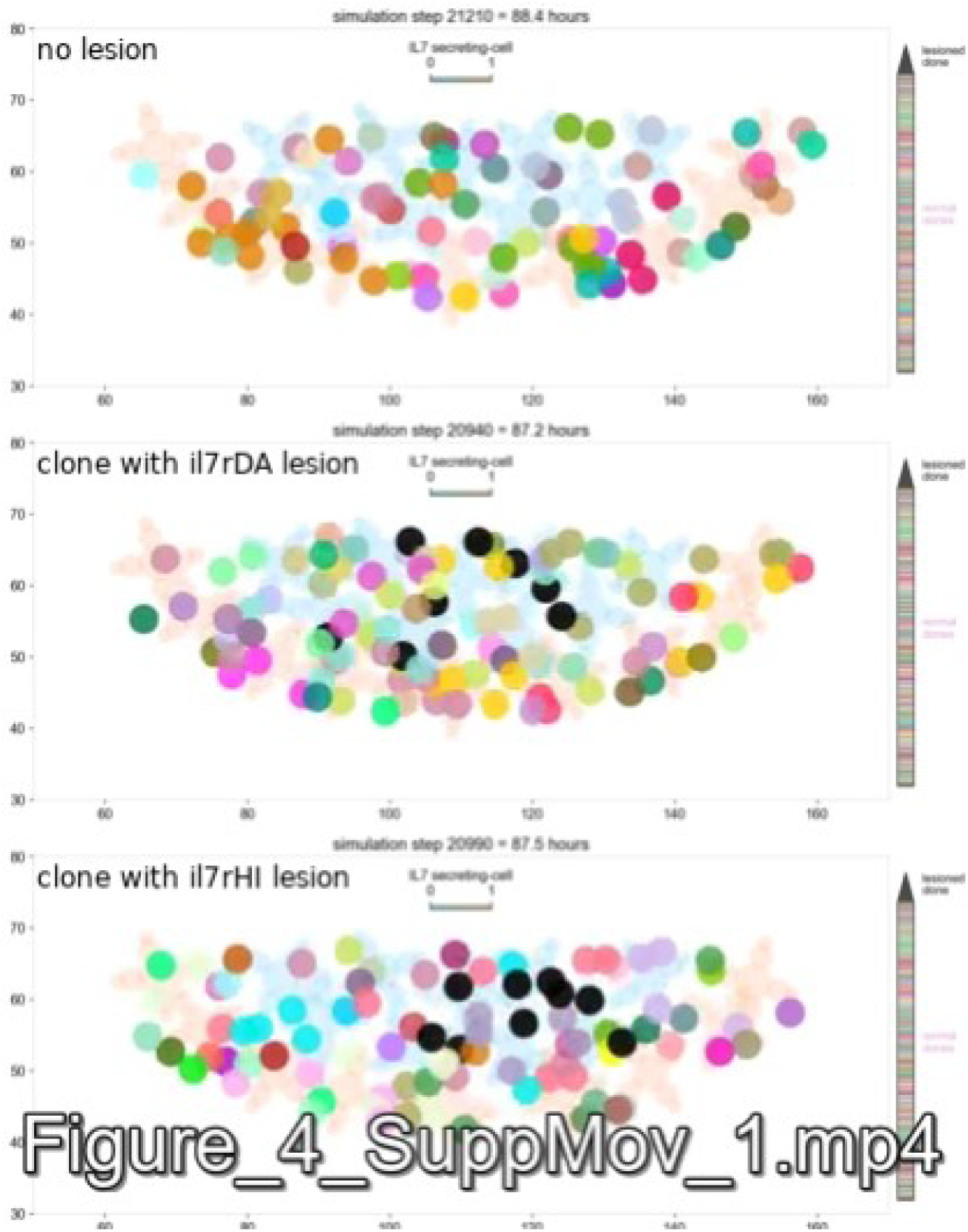
Simulation time-lapse with lesioned clone. Two-dimensional projections of the virtual thymus volume showing time-lapse of simulations representing conditions in Figure 4B. Each thymocyte lineage was assigned a unique color; the lesioned clone was colored in black. Representative position of TECs shown in light blue (non-secreting) and light orange (IL-7-secreting). Movies were synchronized to the time of entry of the first thymocyte.

**Figure 4 - Supplement 2.**
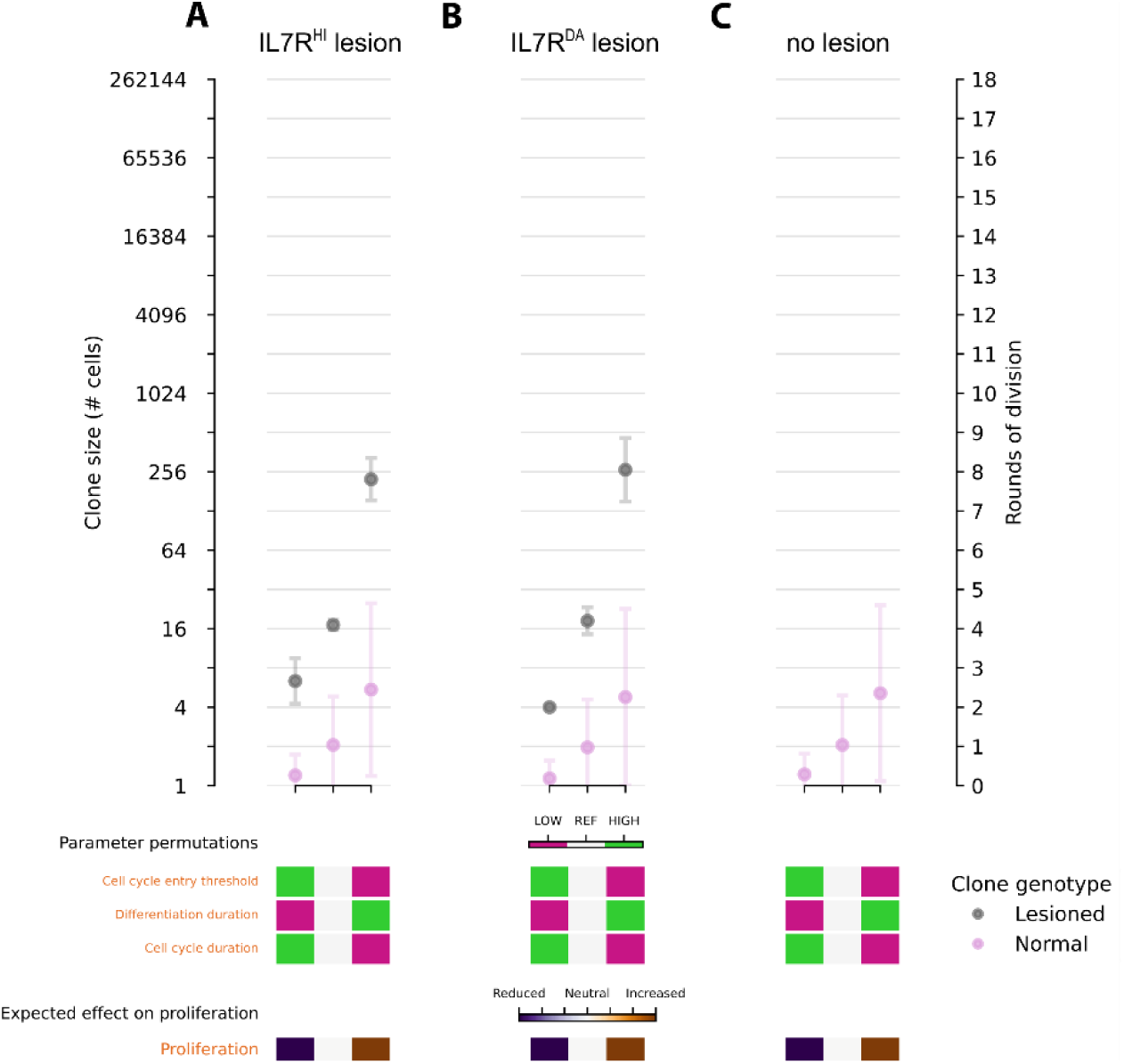
Modulation of proliferation-related parameters affects lesioned thymocytes more strongly. (A) Clones with IL7R^HI^ lesion. (B) Clones with IL7R^DA^ lesion. (C) Control simulations without lesioned clones. Note that the y-axis of all plots is base-2 logarithmic to better illustrate rounds of cell division. Error bars indicate standard deviation. By modulating proliferation parameters, lesioned clones gain up to 6 rounds of division, while non-lesioned clones gain only 2 rounds of division. Expected effect on proliferation is a qualitative estimate based on preliminary simulations.

**Figure 4 - Supplement Movie 2.**
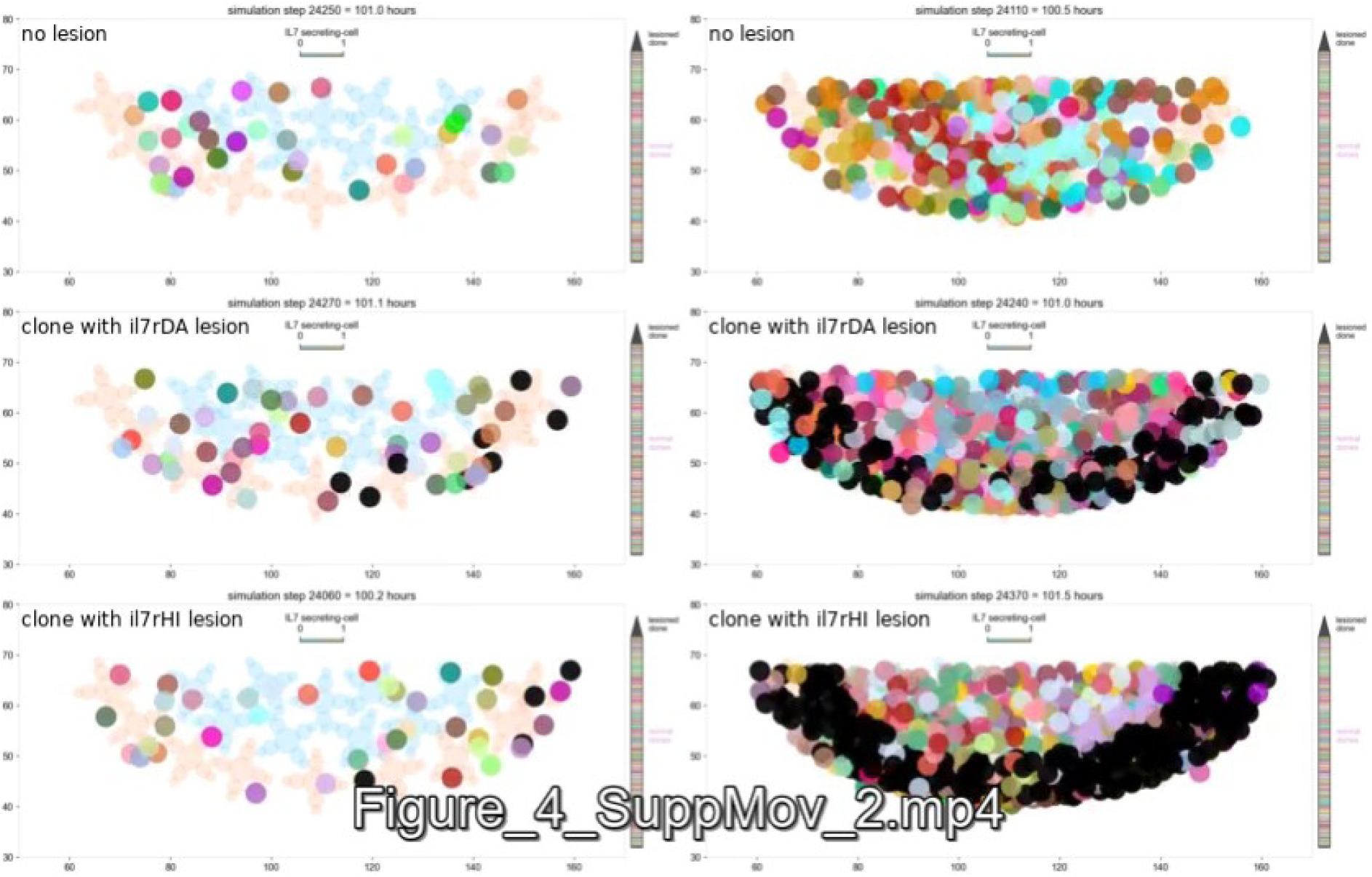
Simulation time-lapse. Two-dimensional projections of the virtual thymus volume showing time-lapse of simulations representing conditions in Figure 4 – Supplement 1A-C. Specifically, left column represents the condition with reduced proliferation, and the right column the condition with increased proliferation. Note how, with increased proliferation, newcomer cells struggle to enter deeper into the organ due to cell crowding. Each thymocyte lineage was assigned a unique color; the lesioned clone was colored in black. Representative position of TECs shown in light blue (non-secreting) and light orange (IL-7-secreting). Movies were synchronized to the time of entry of the first thymocyte.

**Figure 4 - Supplement 3.**
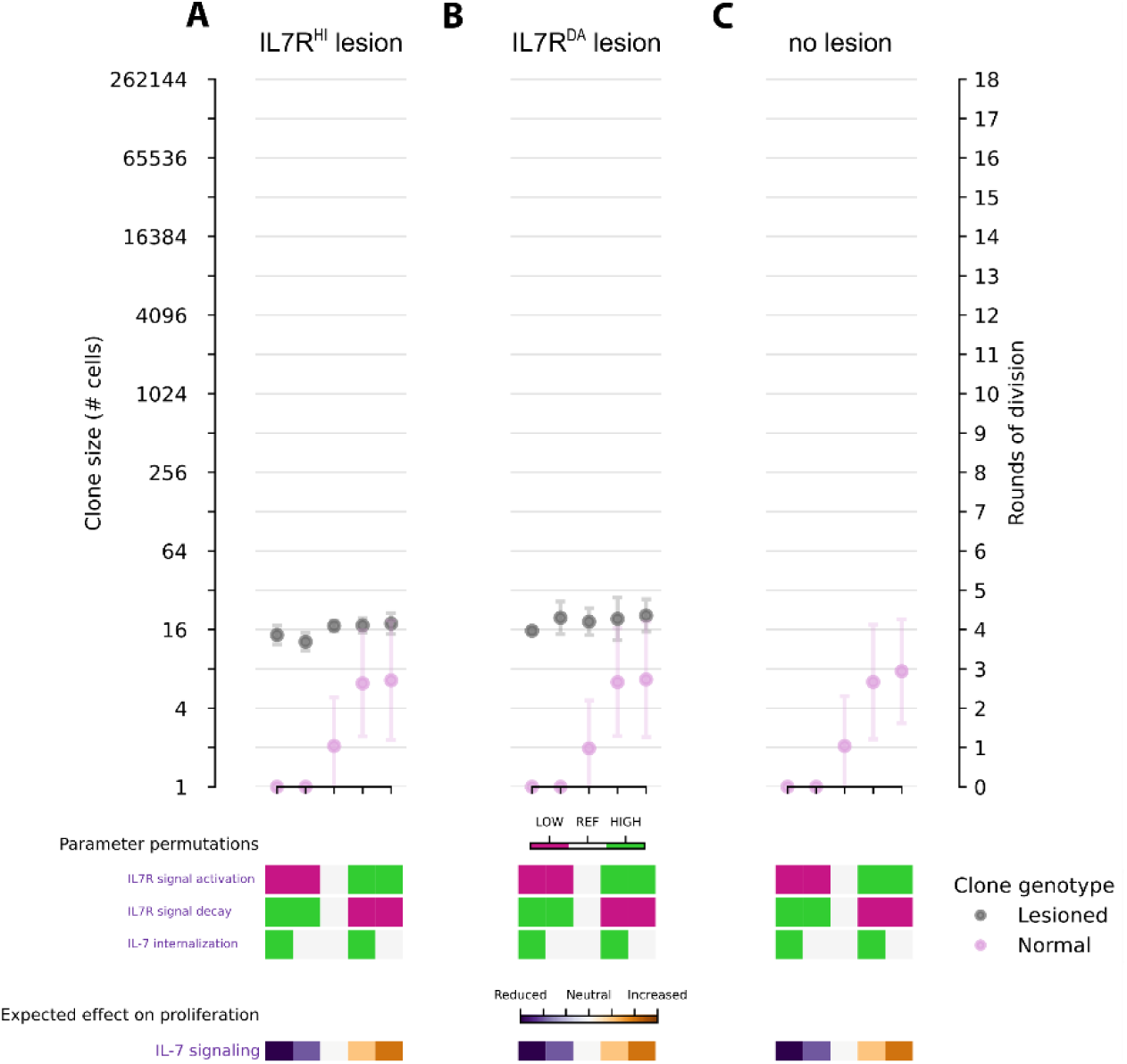
Modulation of parameters affecting IL7-signaling have almost no effect on lesioned clones. (A) Clones with IL7R^HI^ lesion. (B) Clones with IL7R^DA^ lesion. (C) Control simulations without lesioned clones. Note that the y-axis of all plots is base-2 logarithmic to better illustrate rounds of cell division. Error bars indicate standard deviation. By modulating IL-7 signaling parameters, lesioned clones do not gain extra rounds of division, while non-lesioned clones gain up to 3 rounds of division. Expected effect on proliferation is a qualitative estimate based on preliminary simulations.

**Figure 4 - Supplement 4.**
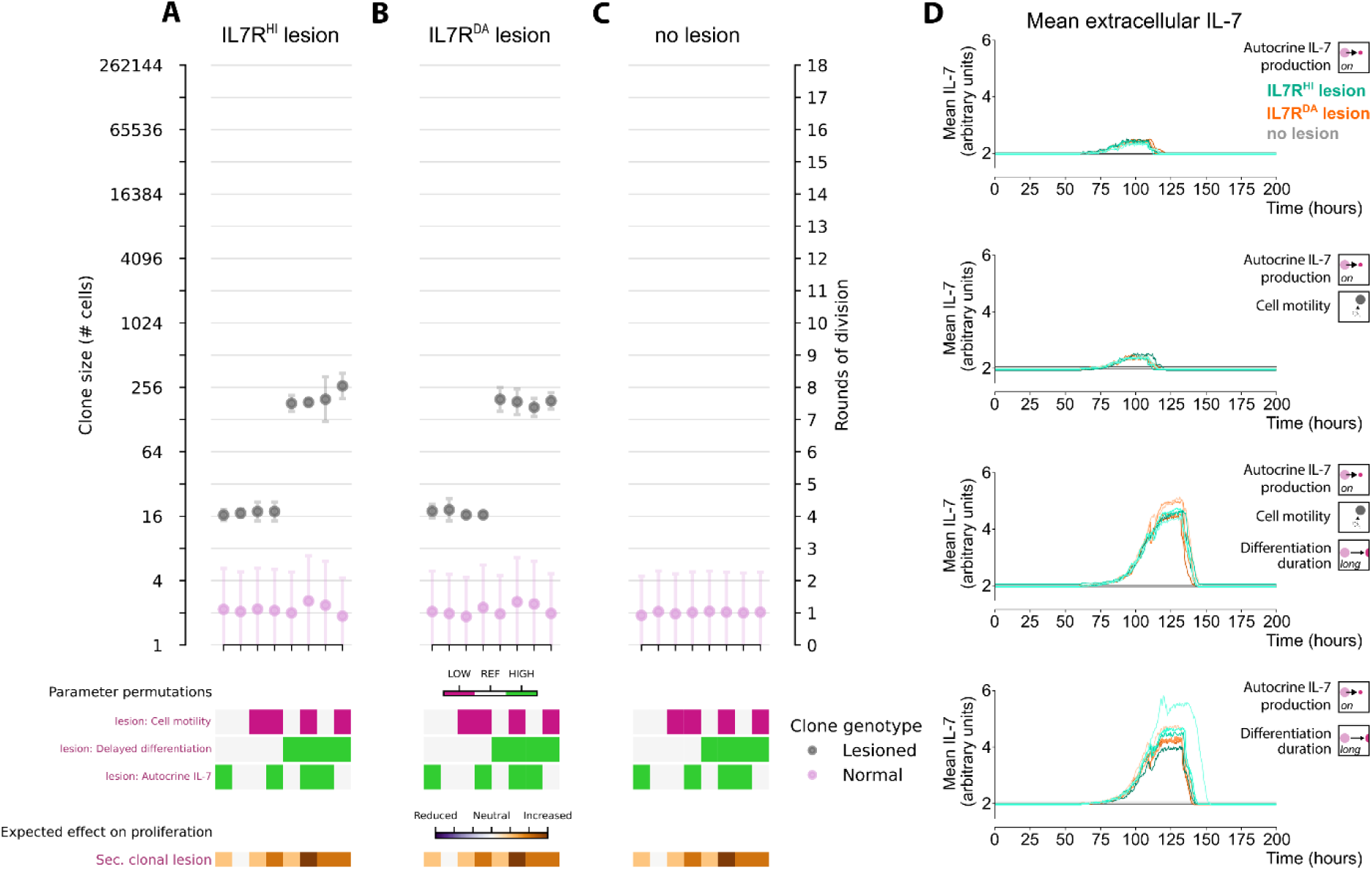
A secondary lesion delaying differentiation doubles proliferative potential. (A) Clones with IL7R^HI^ lesion. (B) Clones with IL7R^DA^ lesion. (C) Control simulations without lesioned clones. Note that the y-axis of all plots is base-2 logarithmic to better illustrate rounds of cell division. Error bars indicate standard deviation. Secondary clonal lesions lead to a gain of 4 rounds of division in the lesioned clones, with no effect on non-lesioned clones. Expected effect on proliferation is a qualitative estimate based on preliminary simulations. (D) Mean extracellular IL-7 concentration over time for selected simulations shown in (A)-(C). The increase in extracellular IL-7 by autocrine activity of lesioned clones is not enough to promote substantial proliferation of non-lesioned cells.

**Figure 4 - Supplement 5.**
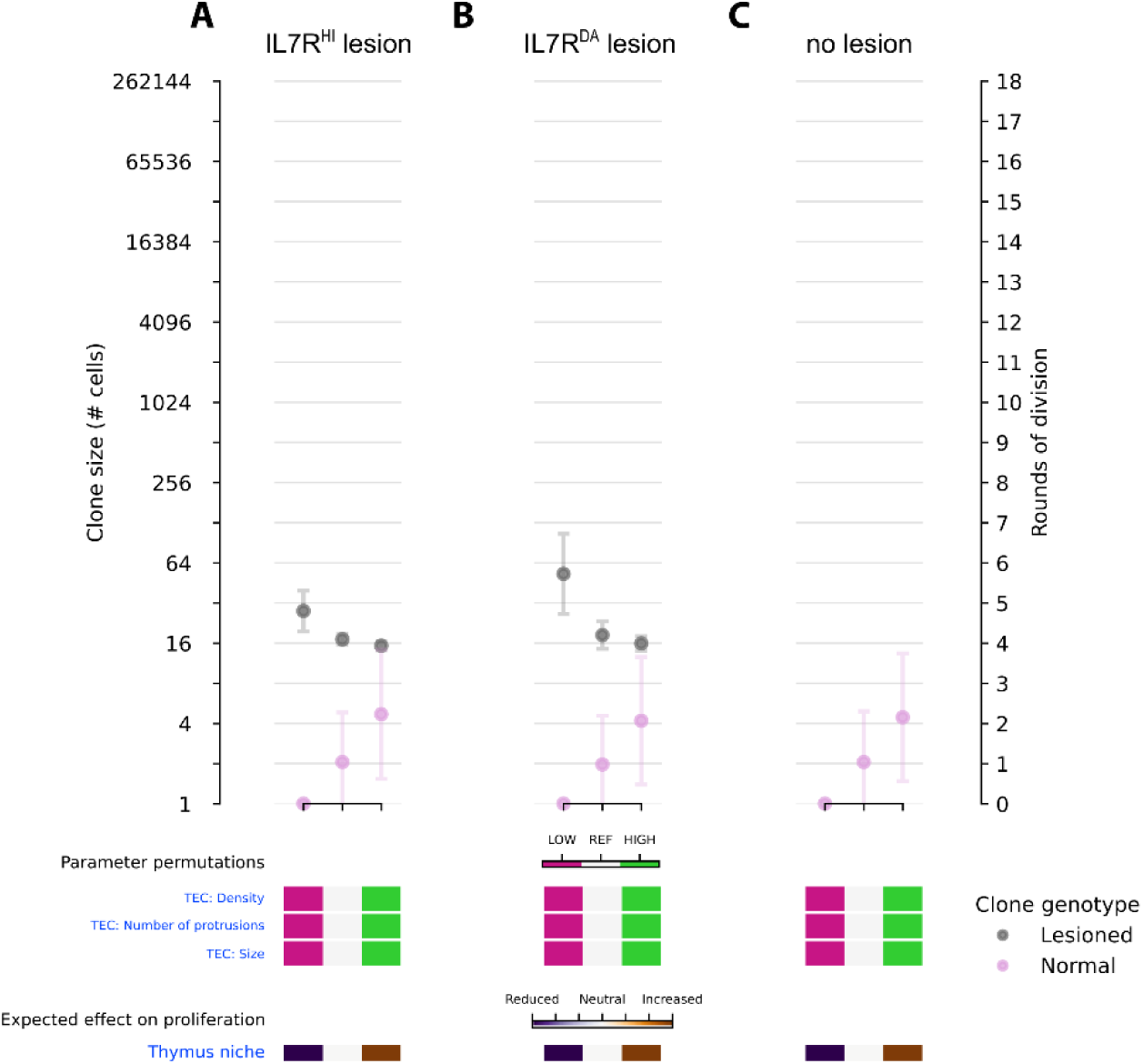
Parameters affecting TEC distribution have opposing effects on lesioned and non-lesioned clones. (A) Clones with IL7R^HI^ lesion. (B) Clones with IL7R^DA^ lesion. (C) Control simulations without lesioned clones. Note that the y-axis of all plots is base-2 logarithmic to better illustrate rounds of cell division. As TEC architecture is made denser, lesioned clones lose up to 2 rounds of cell division, while non-lesioned clones gain 2 rounds of cell division. Expected effect on proliferation is a qualitative estimate based on preliminary simulations.

**Figure 4 - Supplement 6.**
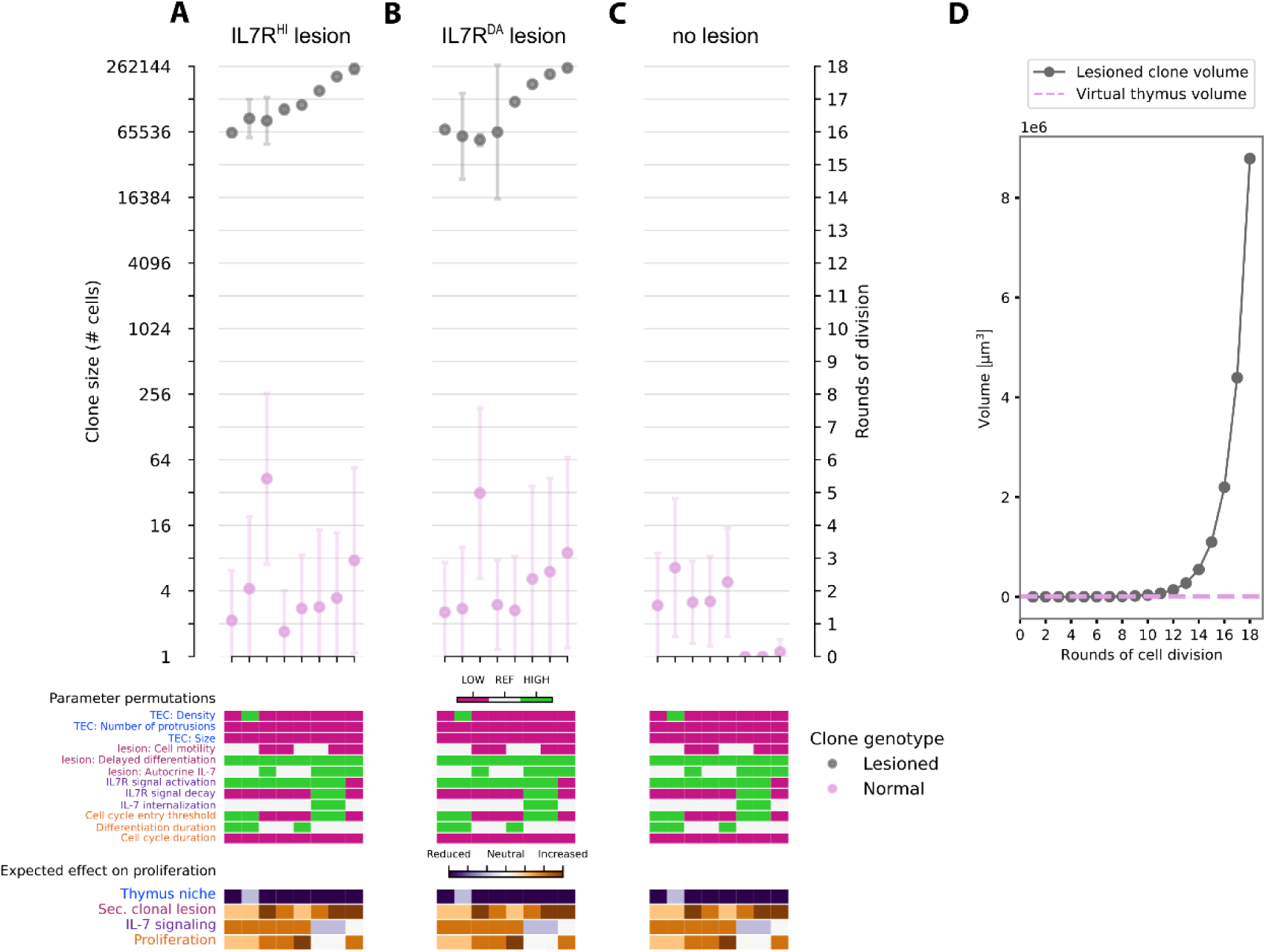
Sparse TEC distribution is among permutations that most enabled lesioned clone expansion. Shown are top 8 conditions from Figure 4 (E). (A) Clones with IL7R^HI^ lesion. (B) Clones with IL7R^DA^ lesion. (C) Control simulations without lesioned clones. Note how lesioned clones with autocrine IL-7 production favor expansion of non-lesioned clones as well. The y-axis of plots in (A)-(C) is base-2 logarithmic to better illustrate rounds of cell division. Points are mean values, error bars indicate standard deviation. Expected effect on proliferation is a qualitative estimate based on preliminary simulations. (D) Volume occupied by thymocytes (1 thymocyte ≅ 33.5 μm^3^) of a hypothetical lesioned clone compared to the volume of the virtual thymus (≅ 7854 μm^3^). The y-axis is scaled in millions of cubic micrometers. The volumes strongly diverge after 12 rounds of cell division indicating extremely dense cell packing that would result in hyperplasia or metastasis *in vivo*.

**Figure 5 - Supplement 1.**
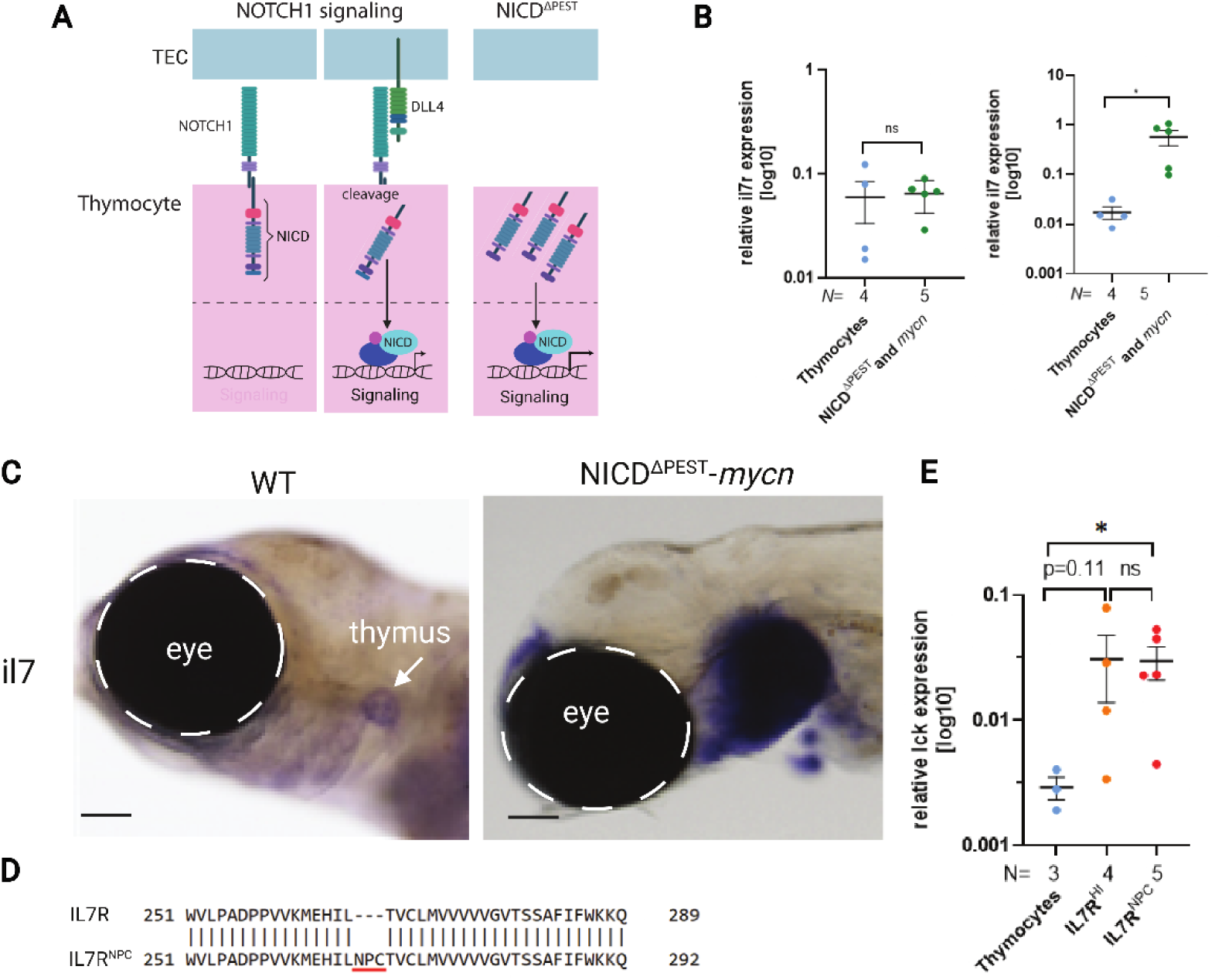
Increased *il7* expression upon NICD^ΔPEST^-mycn overexpression. (A) Schematic depiction of NOTCH1 signaling (left) and overexpression of NICD (right). The engagement of NOTCH1 receptor with its ligand (DLL4) leads to its cleavage, and release and translocation of the NICD in the nucleus. NICD through interaction with other co-factors and triggers the expression of the target genes. Overexpression of NICD leads to activation of NOTCH signaling independent of the ligand. (B) Relative expression of *il7r* and *il7* of isolated thymocytes from WT compared to those expressing both NICD^ΔPEST^ and *mycn*. Each dot represents data from an individual sorted embryo. Data are log10 means ± standard error of the mean. Statistics: unpaired Mann-Whitney test, significance threshold was set to 0.05. (C) WISH with the il7 probe of freshly hatched yolk sac WT and NICD^ΔPEST^ and *mycn* larvae. Scale bare is 100µm. (D) Pairwise sequence alignment was done using EMBOSS Needle and the Needleman-Wunsch algorithm (Madeira et al. 2022), highlighting the partial medaka IL7R protein sequence and IL7R^NPC^ insertion mutation of amino acids NPC at position 266 in the extracellular juxtamembrane-transmembrane interface. (E) Relative *lck* expression quantified by qPCR of sorted WT thymocytes compared to those in IL7R^HI^ and IL7R^NPC^. The *lck* expression was normalized to the housekeeping gene *ef1a*. Each dot represents an individual sorted embryo. Data are log10 means ± standard error of the mean. Statistics: unpaired Mann-Whitney test. Significance threshold was set to p=0.05. * means p < 0.05.

**Figure 5 - Supplement Table 1:** Detailed statistics for phenotype comparison using Fisher’s exact test and pairwise comparisons; adjusted p values were calculated with the Benjamini & Hochberg method. Values reported in figures are the adjusted p values. Significance threshold was set to 0.05. Data rounded to 4 decimal places.

| Group 1 | Group 2 | p.value | significant | p.value adjusted | significant adjusted |
| --- | --- | --- | --- | --- | --- |
| WT | lck:NICD | 1 | FALSE | 1 | FALSE |
| WT | ccl25a:il7 | 0.0486 | TRUE | 0.0694 | FALSE |
| WT | lck:NICD-MycN | 0.0007 | TRUE | 0.0017 | TRUE |
| WT | ccr9a:il7r | 0.0002 | TRUE | 0.0006 | TRUE |
| WT | ccr9a:il7r + ccl25a:il7 | 4.90E-06 | TRUE | 2.20E-05 | TRUE |
| WT | lck:NICD-MycN + ccl25a:il7 | 4.15E-08 | TRUE | 4.67E-07 | TRUE |
| WT | ccr9a:il7r<sup>NPC</sup> | 8.70E-09 | TRUE | 1.31E-07 | TRUE |
| WT | lck:NICD-MycN + ccr9a:il7r | 2.70E-07 | TRUE | 2.03E-06 | TRUE |
| WT | lck:NICD-MycN + ccr9a:il7r<sup>NPC</sup> | 3.00E-12 | TRUE | 1.47E-10 | TRUE |
| lck:NICD | ccl25a:il7 | 0.0730 | FALSE | 0.0996 | FALSE |
| lck:NICD | lck:NICD-MycN | 0.0040 | TRUE | 0.0076 | TRUE |
| lck:NICD | ccr9a:il7r | 0.0011 | TRUE | 0.0024 | TRUE |
| lck:NICD | ccr9a:il7r + ccl25a:il7 | 5.2885E-05 | TRUE | 0.0002 | TRUE |
| lck:NICD | lck:NICD-MycN + ccl25a:il7 | 6.92E-07 | TRUE | 4.42E-06 | TRUE |
| lck:NICD | ccr9a:il7r<sup>NPC</sup> | 9.67E-08 | TRUE | 8.70E-07 | TRUE |
| lck:NICD | lck:NICD-MycN + ccr9a:il7r | 1.59E-06 | TRUE | 7.93E-06 | TRUE |
| lck:NICD | lck:NICD-MycN + ccr9a:il7r<sup>NPC</sup> | 1.24E-10 | TRUE | 2.78E-09 | TRUE |
| ccl25a:il7 | lck:NICD-MycN | 0.4027 | FALSE | 0.4531 | FALSE |
| ccl25a:il7 | ccr9a:il7r | 0.2344 | FALSE | 0.2776 | FALSE |
| ccl25a:il7 | ccr9a:il7r + ccl25a:il7 | 0.0351 | TRUE | 0.0526 | FALSE |
| ccl25a:il7 | lck:NICD-MycN + ccl25a:il7 | 0.0025 | TRUE | 0.0049 | TRUE |
| ccl25a:il7 | ccr9a:il7r<sup>NPC</sup> | 0.0008 | TRUE | 0.0018 | TRUE |
| ccl25a:il7 | lck:NICD-MycN + ccr9a:il7r | 0.0007 | TRUE | 0.0017 | TRUE |
| ccl25a:il7 | lck:NICD-MycN + ccr9a:il7r<sup>NPC</sup> | 8.65E-06 | TRUE | 3.54E-05 | TRUE |
| lck:NICD-MycN | ccr9a:il7r | 0.0872 | FALSE | 0.1155 | FALSE |
| lck:NICD-MycN | ccr9a:il7r + ccl25a:il7 | 0.1314 | FALSE | 0.1642 | FALSE |
| lck:NICD-MycN | lck:NICD-MycN + ccl25a:il7 | 0.0493 | TRUE | 0.0694 | FALSE |
| lck:NICD-MycN | ccr9a:il7r<sup>NPC</sup> | 0.0170 | TRUE | 0.0283 | TRUE |
| lck:NICD-MycN | lck:NICD-MycN + ccr9a:il7r | 0.0281 | TRUE | 0.0436 | TRUE |
| lck:NICD-MycN | lck:NICD-MycN + ccr9a:il7r<sup>NPC</sup> | 0.0003 | TRUE | 0.0007 | TRUE |
| ccr9a:il7r | ccr9a:il7r + ccl25a:il7 | 0.2187 | FALSE | 0.2660 | FALSE |
| ccr9a:il7r | lck:NICD-MycN + ccl25a:il7 | 0.0003 | TRUE | 0.0007 | TRUE |
| ccr9a:il7r | ccr9a:il7r<sup>NPC</sup> | 6.7619E-05 | TRUE | 0.0002 | TRUE |
| ccr9a:il7r | lck:NICD-MycN + ccr9a:il7r | 8.0909E-05 | TRUE | 0.0003 | TRUE |
| ccr9a:il7r | lck:NICD-MycN + ccr9a:il7r<sup>NPC</sup> | 7.85E-07 | TRUE | 4.42E-06 | TRUE |
| ccr9a:il7r + ccl25a:il7 | lck:NICD-MycN + ccl25a:il7 | 0.0258 | TRUE | 0.0415 | TRUE |
| ccr9a:il7r + ccl25a:il7 | ccr9a:il7r<sup>NPC</sup> | 0.0135 | TRUE | 0.0234 | TRUE |
| ccr9a:il7r + ccl25a:il7 | lck:NICD-MycN + ccr9a:il7r | 0.0078 | TRUE | 0.0141 | TRUE |
| ccr9a:il7r + ccl25a:il7 | lck:NICD-MycN + ccr9a:il7r<sup>NPC</sup> | 0.0016 | TRUE | 0.0033 | TRUE |
| lck:NICD-MycN + ccl25a:il7 | ccr9a:il7r<sup>NPC</sup> | 0.7902 | FALSE | 0.8269 | FALSE |
| lck:NICD-MycN + ccl25a:il7 | lck:NICD-MycN + ccr9a:il7r | 0.6492 | FALSE | 0.6955 | FALSE |
| lck:NICD-MycN + ccl25a:il7 | lck:NICD-MycN + ccr9a:il7r<sup>NPC</sup> | 0.1243 | FALSE | 0.1598 | FALSE |
| ccr9a:il7r<sup>NPC</sup> | lck:NICD-MycN + ccr9a:il7r | 0.9308 | FALSE | 0.9519 | FALSE |
| ccr9a:il7r<sup>NPC</sup> | lck:NICD-MycN + ccr9a:il7r<sup>NPC</sup> | 0.4821 | FALSE | 0.5291 | FALSE |
| lck:NICD-MycN + ccr9a:il7r | lck:NICD-MycN + ccr9a:il7r<sup>NPC</sup> | 0.3705 | FALSE | 0.4275 | FALSE |

