## Appendix for "Altered thymic niche synergistically drives the massive proliferation of malignant thymocytes"

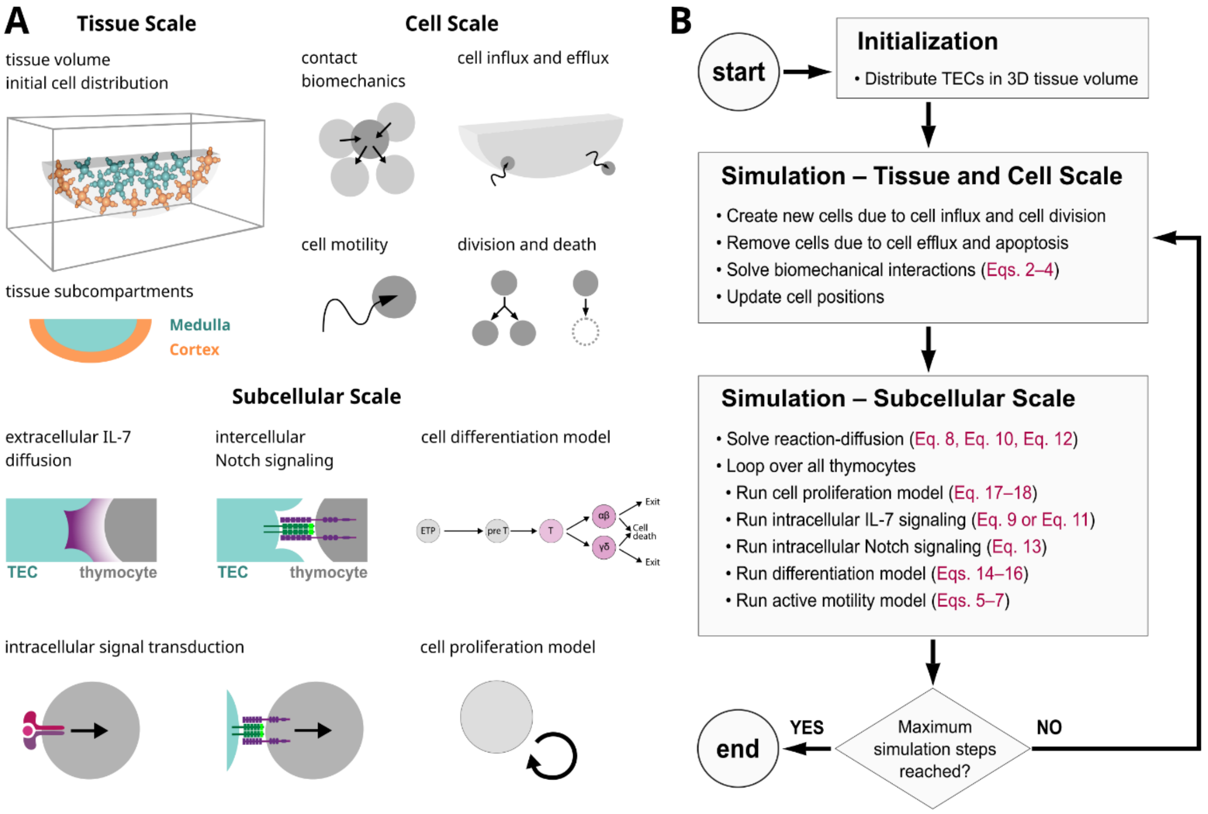


**Appendix Figure 1. Overview of simulation components.** A) Schematic representation of model components at the tissue, cell, and subcellular scale. B) Flowchart illustrating the model operation with references to equations shown in the text.

### Appendix

#### The virtual thymus model

The cell-based model of the virtual thymus was implemented in the platform EPISIM, which simulates center-based biomechanical interactions for spherical or ellipsoid cells [Sütterlin et al. 2013, Sütterlin et al. 2017].

A detailed description of the mathematical underpinning and biological inspiration of the model and its parametrization was published previously [Aghaallaei et al. 2021]. Parameters were adapted from this previous work; a few additional scaling parameters were added to enable differential behavior in lesioned thymocytes (Appendix Table 1, Appendix Table 2). The code and model files of the most recent version associated with this publication are freely available online on a Zenodo repository [Tsingos 2024]. In the following, we recapitulate the main features of the model implementation for the reader's convenience.

The virtual thymus model represents processes in the thymus at the tissue, cellular, and subcellular scales (Appendix Figure A1 A–B). At the tissue scale, we model a thin three-dimensional (3D) slice of the thymus of medaka hatchlings.

At the cellular scale we model thymic epithelial cells (TECs) and thymocytes, and both cell types are represented as one or more spherical particles. Biomechanical interactions modelled at the cellular scale are cell motility and cell-cell contact mechanics. This scale also includes processes that change the number of cells such as cell influx into and efflux out of the tissue volume, cell proliferation, and cell death.

At the subcellular scale, we model biochemical interactions such as extracellular diffusion of cytokines, signaling between cells, intracellular signal transduction, and include phenomenological models of the thymocyte cell differentiation and proliferation programs. In the following sections, we provide an overview of each of these processes, and refer to original publications for further details where appropriate.

**Appendix Table 1 - Parameters defining initial condition**

All parameters were adapted from Aghaallaei et al. 2021 unless noted.

| Symbol | Reference value | Description |
|---|---|---|
| $a$ | 50 µm | Long hemi-axis of thymus cylindrical ellipse. |
| $b$ | 25 µm | Short hemi-axis of thymus cylindrical ellipse. |
| $h_c$ | 5 µm | Height of thymus cylindrical ellipse. |
| $g$ | 2 µm | Spatial discretization for solving IL-7 reaction-diffusion equation. |
| $t$ | 15 s | Duration of one simulation step. |
| $r_{\mathrm{TEC}}$ | 2.5 µm | Radius of the main TEC body. Also scales the radius of the spheres in the TEC protrusions. |
| $r_{\mathrm{C}}$ | 2 µm | Radius of thymocytes. |
| $N$ | 3 | Number of concentric subdivisions in algorithm to generate TEC positions. The higher the value, the larger the number of TECs, thus the larger the TEC density. |
| $p_{\mathrm{TEC}}$ | 4 | Number of protrusions initialized for each TEC. |
| *DEPL* | False | New parameter introduced in this work. Boolean flag that sets if the simulation operates with thymocytes as IL-7 sinks according to Eqs. 10 and 12 (True) or if it defaults to having no sink terms (False). |

**Appendix Table 2 - Simulation parameters**

All parameters were adapted from Aghaallaei et al. 2021 unless noted.

| Symbol | Reference value | Description |
|---|---|---|
| $\gamma$ | $4 \cdot 10^{-7}$ N·s·µm$^{-1}$ | Friction of the environment. |
| $k_{spawn}$ | 0.45 h$^{-1}$ | Rate of thymocyte entry into the simulation from two separate positions in the simulation box, resulting in a net influx of 0.9 thymocytes per hour. |
| $\delta_{adh}$ | 1.3 | Scaling factor to define neighborhood and adhesive interaction distances. |
| $\delta_{ol\_max}$ | 0.5 | Scaling factor to define the inner repulsive interaction zone. |
| $\delta_{ol}$ | 0.85 | Scaling factor to define the outer repulsive interaction zone. |
| $d_{ol\_min}$ | 0.1 µm | Defines a neutral zone of no adhesive or repulsive forces. |
| $\mu_{S,\ imm}$ | 720 µm·h$^{-1}$ | Mean speed of cells immigrating into or emigrating out of the thymus. |
| $\mu_{S,\ res}$ | 150 µm·h$^{-1}$ | Mean speed of cells resident in the thymus |
| $\sigma_S$ | 0.5 | Scaling factor for cell speed variance. |
| $\tau$ | 1 (wildtype, non-resident)<br>$\tau_{diff}$ (wildtype, resident)<br>0.5 (speed lesion) | Scales the cell speed. In this work, the possibility of including a speed lesion was introduced by setting the parameter to 0.5 for lesioned clones only. |
| $D_{IL\text{-}7}$ | $1 \cdot 10^{-12}$ m$^2$·s$^{-1}$ | Extracellular IL-7 diffusion coefficient. |
| $k_{IL\text{-}7}$ | 0.0167 h$^{-1}$ | Extracellular IL-7 degradation rate. |
| $a_{IL\text{-}7}$ | 240 h$^{-1}$ | New parameter introduced in this work. IL7R signaling activation rate (reference value equivalent to previous work). |
| $d_{IL\text{-}7}$ | 50 h$^{-1}$ | IL7R signaling deactivation rate. |
| $k_{\sigma Notch}$ | 0.029 h$^{-1}$ | Notch signaling deactivation rate. |
| $\kappa$ | 0.3 | Notch-independent differentiation rate. |
| $T_{diff}$ | 24 h | Minimum duration of the proliferative phase before terminal differentiation. |
| $d_{diff\_lesion}$ | 1 (wildtype)<br>0.5 (differentiation lesion) | New parameter introduced in this work. Scales the rate of differentiation. |
| $\theta_{diff}$ | 0.4 | Threshold level of IL-7 signaling activity required to differentiate into $\gamma\delta^+$ T cell subtype. |
| $T_{mat}$ | 24 h | Duration of the maturation phase before thymic selection |
| $\mu$ | 7 h | Mean cell cycle duration. |
| $k$ | 50 | Shape parameter scaling the variance of the cell cycle distribution function. |
| $t_M$ | 0.5 h | Duration of the M phase of the cell cycle. |
| $\kappa_S$ | 0.5 | Fraction of the cell cycle duration allocated to the S phase. |
| $\theta_{prol}$ | 1.4 | Threshold level of IL-7 and Notch signaling activity required to progress in G1 phase and to commit to the cell cycle. |

#### I. Tissue scale

We define a simulation box of dimensions 220 x 100 x 60 micrometres. At the center of this box we define a subregion that represents a slice of the thymus volume as a cylindrical hemi-ellipse with

$$\begin{aligned} a &= 50 \\ b &= 25 \\ h_C &= 5 \end{aligned} \qquad \text{(Eq. 1)}$$

where $a$ and $b$ are the elliptical hemi-axes, and $h_C$ is the height of the cylinder representing a slice of the organ; all values are given in micrometres. Furthermore, we subdivide the thymus into two concentric tissue compartments: the inner medulla and the outer cortex (Figure A1 A). The cortex is defined as the region within the outermost 20% of elliptical hemi-axes $a$ and $b$. Since the thymus is a symmetric ellipsoid in young medaka fish, we chose to represent only a slice of the organ to speed up computation and facilitate analysis. The dimensions of the modelled slice were chosen based on measurements of the medaka thymus at 11 days post fertilization [Bajoghli et al. 2015].

#### II. Tissue Scale: Initial condition

We initialize the model by distributing TECs in the 3D thymus volume defined by the cylindrical hemi-ellipse in (Eq. 1). The algorithm has been described previously [Aghaallaei et al. 2021]. Briefly: We subdivide the thymus volume into a number of sub-domains, and in each subdomain we select a random point with uniform probability; the selected point is then the position of the central sphere representing a TEC's main body. The protrusions of TECs are initialized in four predefined directions starting from the main body with an added small random variation in angle to create irregularity. The radii of TEC protrusion particles are scaled down to generate a gradually tapered protrusion.

#### III. Cellular Scale: Biomechanical interactions

We phenomenologically model passive forces arising from biomechanical contact mechanics, which include intercellular adhesion due to cell-cell adhesion proteins and pressure due to volume exclusion. In addition, we include active forces resulting from cell motility. As the length scale of cells is on the order of 10 micrometers, we assume low Reynolds numbers and an overdamped environment [Purcell 1977, Berg 1993]. Thus, the force-balance equation for a particle $i$ at position $\mathbf{c}_i$ is given by

$$\gamma \frac{\mathrm{d}}{\mathrm{d}t}\mathbf{c}_i = \mathbf{F}_{\mathrm{contact}} + \mathbf{F}_{\mathrm{act}} \qquad \text{(Eq. 2)}$$

where $\gamma$ is the friction or damping constant of the environment, $\mathbf{F}_{\mathrm{contact}}$ is the sum of passive contact forces acting on particle $i$ due neighbouring particles in its proximity and $\mathbf{F}_{\mathrm{act}}$ is the active force exerted by the particle $i$ due to its intrinsic motility. The force balance equation is integrated with an explicit forward Euler scheme.

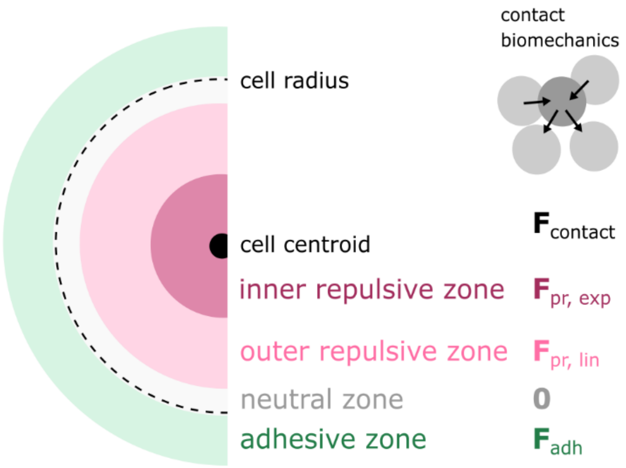


**Appendix Figure 2. Biomechanical interaction zones.** Schematic representation of the different interaction zones that are used for calculating biomechanical contact forces. The radii of the interaction zones in the image have been scaled assuming two spherical particles of identical radius.

To simplify the computational implementation, the particles that make up each TEC are fixed in space. In other words, TEC particles are included in the force balance calculation (Eq. 2), but their position in 3D space is not updated. This enables TECs to act as obstacles to thymocytes, but ignores dynamic shape changes and motility of TECs themselves. This simplification is a limitation of the current model.

##### Contact forces

The details of the mathematical implementation of contact forces have been published elsewhere [Sütterlin et al. 2017]. In essence, contact forces are made up of repulsive pressure forces that push particles apart and attractive adhesion forces that pull neighbouring particles toward each other. Two particles *i* and *j* are considered neighbours if

$$d_{ij} \leq d_{\mathrm{opt}}(i,j) \cdot \delta_{\mathrm{adh}} \qquad \text{(Eq. 3)}$$

where $d_{ij}$ is the Euclidean distance of the particle centroids, $d_{opt}(i,j)$ is an optimal distance obtained from the respective dimensions of particle *i* and its neighbour particle *j* (for the spherical particle interaction radii that we use in this work the calculation of $d_{opt}(i,j)$ simplifies to the sum of radii of *i* and *j*), and $\delta_{adh} > 1$ is a parameter that defines the neighbourhood interaction distance.
The contact forces for each particle-neighbour pair are given by a piecewise function with multiple distance thresholds based on $d_{ij}$ and $d_{opt}(i,j)$, which results in concentric interaction zones (illustrated in Appendix Figure 2):

$$\mathbf{F}_{\mathrm{contact}} = \begin{cases} \mathbf{F}_{\mathrm{pr,\ exp}} & \text{if} \quad 0 < d_{ij} < d_{\mathrm{opt}}(i,j) \cdot \delta_{\mathrm{ol}} \cdot \delta_{\mathrm{ol\_max}} \\ \mathbf{F}_{\mathrm{pr,\ lin}} & \text{if} \quad d_{\mathrm{opt}}(i,j) \cdot \delta_{\mathrm{ol}} \cdot \delta_{\mathrm{ol\_max}} \leq d_{ij} < d_{\mathrm{opt}}(i,j) \cdot \delta_{\mathrm{ol}} - d_{\mathrm{ol\_min}} \\ \mathbf{0} & \text{if} \quad d_{\mathrm{opt}}(i,j) \cdot \delta_{\mathrm{ol}} - d_{\mathrm{ol\_min}} \leq d_{ij} < d_{\mathrm{opt}}(i,j) + d_{\mathrm{ol\_min}} \\ \mathbf{F}_{\mathrm{adh}} & \text{if} \quad d_{\mathrm{opt}}(i,j) + d_{\mathrm{ol\_min}} \leq d_{ij} < d_{\mathrm{opt}}(i,j) \cdot \delta_{\mathrm{adh}} \\ \mathbf{0} & \text{if} \quad d_{\mathrm{opt}}(i,j) \cdot \delta_{\mathrm{adh}} \leq d_{ij} \end{cases} \qquad \text{(Eq. 4)}$$

Here, $\delta_{ol\_max}$, $\delta_{ol}$, $d_{ol\_min}$ are parameters that scale the interaction distances. The parameter $\delta_{ol\_max} \in (0, 1)$ defines the inner repulsive interaction zone where the repulsion force $\mathbf{F}_{pr,exp}$ exponentially increases as $d_{ij}$ decreases; $\delta_{ol} \in (0, 1)$ defines the outer repulsive interaction zone where the repulsion force $\mathbf{F}_{pr,lin}$ linearly increases as $d_{ij}$ decreases; $d_{ol\_min} \in (0, 1)$ defines a neutral zone with no net force; $\delta_{adh}$ which is used to determine neighbourhood in (Eq. 3) additionally defines the interaction zone where adhesive forces $\mathbf{F}_{adh}$ dominate (Appendix Figure 2). For the functional form chosen for the force terms we refer to Sütterlin et al. 2017. Based on previous work, the values $\delta_{ol\_max}$ = 0.5; $\delta_{ol}$ = 0.85; $d_{ol\_min}$ = 0.1 µm; $\delta_{adh}$ = 1.3 were chosen as they result in appropriate spacing of particles without mechanical instability [Sütterlin et al. 2017, Tsingos et al. 2019, Aghaallaei et al. 2021]. Note that our approach is consistent with state-of-the-art models of force-based interactions between particles to simulate cells [Pathmanathan et al. 2009, Osborne et al. 2017].

##### Active forces

During the simulation, each cell determines an active instantaneous displacement $\mathbf{d}_{act}$ from its instantaneous speed *s* and a unit vector **v** representing cell orientation

$$\mathbf{d}_{\mathrm{act}} = s \cdot \mathbf{v} \cdot \mathrm{d}t \qquad \text{(Eq. 5)}$$

where *t* is time. Both *s* and **v** are functions of the cell's internal state and its position in the tissue and are dynamically evaluated during the simulation. For an in-depth explanation see the supplementary material of Aghaallaei et al. 2021. In brief, the functional form for the speed *s* was chosen as

$$s = \mu_S \cdot (1 + \sigma_S \mathrm{X}) \cdot \tau \qquad \text{(Eq. 6)}$$

where $\mu_S$ and $\sigma_S$ are model parameters, X ∈ (-1, 1] is a random uniform number, and $\tau$ is either set to $\tau$ = 1 for thymocytes entering or exiting the thymus, or set to $\tau = \tau_{diff}$ for thymocytes currently residing in the thymus (for an explanation on $\tau_{diff}$ see section *IX. Subcellular Scale: Differentiation model*). Essentially, setting $\tau = \tau_{diff}$ enables speed to be linearly increasing with developmental stage, which is an assumption we made based on experimental measurements of thymocyte motility *in vivo* [Bajoghli et al. 2015]. For thymocytes of lesioned clones with reduced cell speed, we set $\tau$ = 0.5. Coefficients $\mu_S$ and $\sigma_S$ were fit to the speed measured in confocal imaging data of thymocytes in medaka; different coefficients for $\mu_S$ were used for thymic immigrant/emigrants and thymic resident cells [Aghaallaei et al. 2021]. Cells from a lesioned clone with slower cell speed had the value of their $\mu_S$ coefficient halved compared to wildtype cells.

The orientation vector **v** is a unit vector obtained by combining a directional vector **u** pointing from the current cell position towards a target location and a random vector **w** that introduces a random bias. The directional **u** components and random **w** components are weighted differently depending on cell location, developmental stage, and each cell's intrinsic expression levels of the cytokine receptor *Ccr9b* (schematically summarized in Appendix Figure 3). For a detailed explanation of the implementation, choice of parameter values, and a sensitivity analysis of tissue homeostasis with respect to parameters we refer to our previous publication [Aghaallaei et al. 2021].

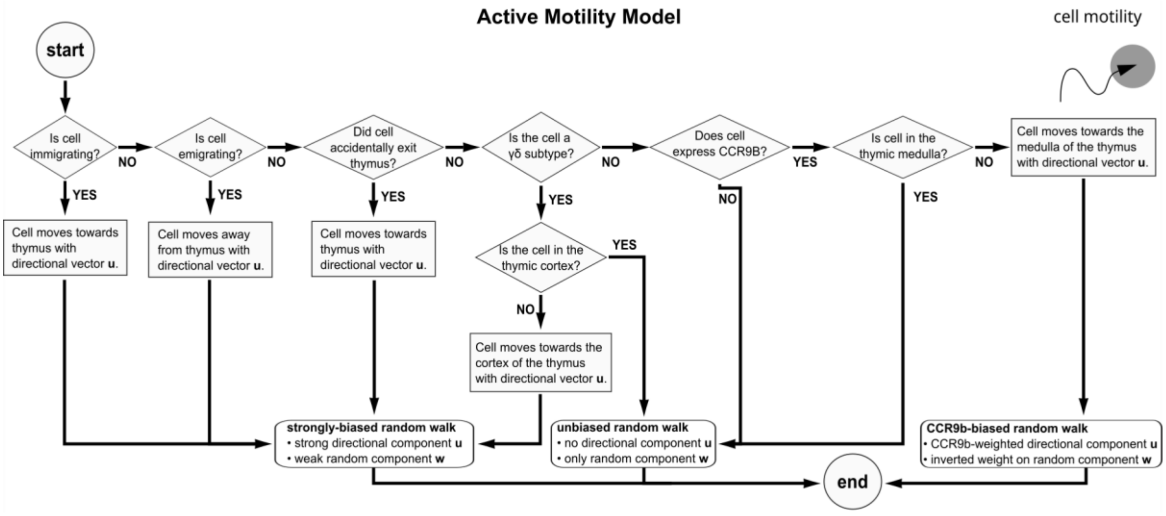


**Appendix Figure 3. Active motility model.** Schematic flowchart illustrating how cell location and state affect cell motility. Early thymic progenitors (ETPs) entering the thymus for the first time ("immigrating"), mature thymocytes leaving the thymus after positive selection ("emigrating"), and thymocytes at various stages of development that accidentally exit the thymus volume ("accidentally exit") have a strong directed component and weak random component to their migration. Otherwise, resident thymocytes at all stages of development do a random walk if they are in the correct thymic compartment, or a biased random walk if they are not in the correct thymic compartment.

The active displacement $\mathbf{d}_{act}$ is used to obtain the cell motility force $\mathbf{F}_{act}$

$$\mathbf{F}_{\text{act}} = \mathbf{d}_{\text{act}} \cdot \frac{\gamma}{\mathrm{d}t} \qquad \text{(Eq. 7)}$$

where $\gamma$ is the friction or damping constant of the environment. In the implementation, the active force $\mathbf{F}_{act}$ obtained in the previous simulation step with (Eq. 7) is used in the equation of motion (Eq. 2) at the beginning of each new simulation step. Because contact forces $\mathbf{F}_{contact}$ to neighbouring cells may hinder cell motility, the effective displacement of cells that results from (Eq. 2) may differ from the displacement calculated in (Eq. 5).

#### IV. Cellular Scale: Cell addition and removal

Cell addition can occur via influx or cell proliferation, and cell removal by efflux or cell death. Note that these mechanisms apply exclusively to thymocytes. The number of TECs is immutable after initialization in the simulation. This simplification is a limitation of the current model.

##### Cell addition

The model includes two mechanisms for adding new cells to the simulated volume: (1) Influx of early undifferentiated thymocytes from areas elsewhere in the body, and (2) increase in cell number due to thymocyte proliferation.

In medaka, thymocytes enter the organ from the ventral side at an estimated influx rate of 9 thymocytes per hour [Bajoghli et al. 2015]. The model implements cell influx by instantiating new cells at two positions at the bottom of the simulation box with rate $k_{spawn}$. Since the model represents about one-tenth of the full organ's volume, we scaled down the influx rate accordingly to $k_{spawn}$ = 0.45 thymocytes per hour, which adds up to 0.9 thymocytes per hour.

Cell proliferation follows a model of the cell division cycle, where progression through G1 phase and commitment to cell division depends on intracellular signaling at the subcellular level (see section *X. Subcellular Scale: Cell proliferation model* for an in-depth explanation). Once the subcellular program of the cell division cycle is complete, the implementation instantiates a new cell adjacent to the cell that completed the cycle at a random orientation; this simulates the close positioning of daughter cells just after cytokinesis with a random spindle orientation.

##### Cell removal

Two processes account for cell removal from the simulation: (1) Efflux of fully differentiated thymocytes that are positively selected, (2) death of fully differentiated thymocytes that are negatively selected. To our knowledge, there is no evidence of substantial cell death at earlier stages of differentiation in medaka [Bajoghli et al. 2015], hence we do not include other mechanisms of cell removal.

Once fully differentiated each thymocyte undergoes thymic selection (see subsection *IX. Subcellular Scale: Differentiation model* for details). Positively selected thymocytes migrate out of the thymus and eventually leave the simulation box, while negatively selected thymocytes stay in position and slowly shrink until their radius reaches 25% of the original value, at which point they are removed from the simulation.

#### V. Subcellular Scale: Extracellular diffusion of IL-7

The extracellular IL-7 concentration $[\text{IL-7}]_{\text{ex}}$ follows the reaction-diffusion equation

$$\frac{\partial}{\partial t}[\text{IL-7}_{\text{ex}}] = D_{\text{IL-7}}\nabla^2[\text{IL-7}_{\text{ex}}] - k_{\text{IL-7}}[\text{IL-7}_{\text{ex}}] + \sum_i s_{\text{SOURCE},i} - \sum_i s_{\text{SINK},i} \qquad \text{(Eq. 8)}$$

where $D_{\text{IL-7}}$ is the diffusion coefficient of extracellular IL-7, $k_{\text{IL-7}}$ is the baseline extracellular degradation of IL-7, and $s_{\text{SOURCE}}$ is a source term summed over the volume of cells that secrete IL-7 and $s_{\text{SINK}}$ is a sink term summed over the volume of cells that consume extracellular IL-7. The diffusion coefficient $D_{\text{IL-7}} = 1 \cdot 10^{-12}$ was chosen based on reported diffusion coefficients for cytokines [Moghe et al. 1995] and the baseline degradation $k_{\text{IL-7}} = 0.0167$ per hour was chosen by parameter scan to produce a gradient that decays within a few cell diameters as reported for other cytokine gradients [Thurley et al. 2015]. In the computational implementation, the parameter value for the secretion rate $s_{\text{SOURCE}}$ of IL-7-secreting TECs is set to 1 concentration unit per voxel of the numerical discretization. This choice ensures that the extracellular IL-7 gradient stays within the interval range of [0, 1] in unscaled concentration units. IL-7-secreting TECs include all TECs whose main cell body is located in the cortex tissue subcompartment, which results in an intra-thymic gradient of IL-7. Thymocytes with a lesion inducing autocrine IL-7 also acted as sources. In the baseline model of Aghaallaei et al. 2021 the sink term in (Eq. 8) is set to zero, while in the present work we compared zero to non-zero sink terms for all thymocytes.

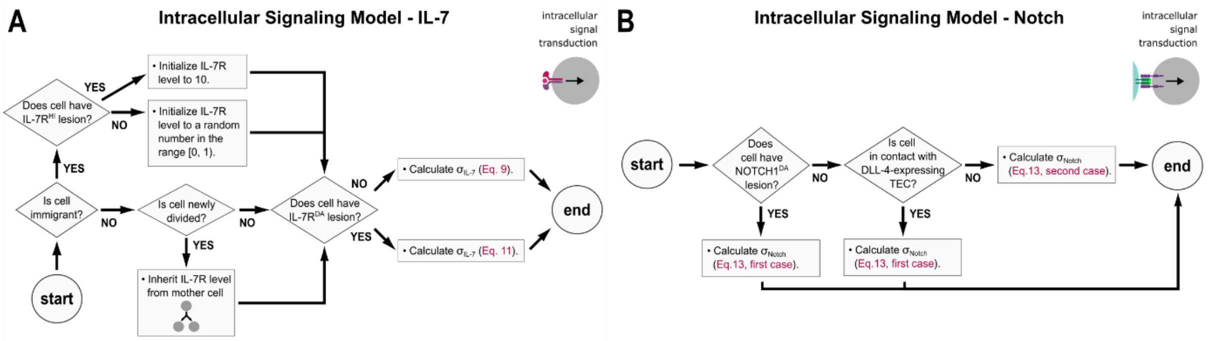


**Appendix Figure 4. Intracellular signal transduction model.** A) Schematic flowchart illustrating the intracellular IL-7 signal transduction model. Early thymic progenitors (ETPs) entering the thymus for the first time are labelled as "immigrants" and undergo an initialization of variables. B) Schematic flowchart illustrating the intracellular Notch signaling model.

The initial condition used to solve the partial differential equation in (Eq. 8) is chosen as uniform zero concentration, and the boundary conditions at the edges of the simulation box are set to a constant zero concentration. In test simulations we verified that the gradient achieves its numerical steady state within as few as 10 simulation steps, which corresponds to 150 seconds, and that the simulation box is sufficiently large to prevent boundary artifacts [Aghaallaei et al. 2021].

#### VI. Subcellular Scale: Intracellular IL-7 signal transduction

The model for intracellular IL-7 signal transduction is summarized in Appendix Figure 4 A. For wildtype thymocytes, the intracellular IL-7 pathway signal transduction level $\sigma_{\text{IL-7}}$ is governed by

$$\frac{\mathrm{d}}{\mathrm{d}t}\sigma_{\text{IL-7}} = [\text{IL-7R}]\langle[\text{IL-7}_{\text{ex}}]\rangle a_{\text{IL-7}} - d_{\text{IL-7}}\sigma_{\text{IL-7}} \qquad \text{(Eq. 9)}$$

where [IL-7R] is the concentration of IL-7 receptor of the cell, 〈[IL-7ex]〉 is the average extracellular IL-7 concentration in the cell's microenvironment (defined as the voxels of the numerical discretization that overlap with the cell's radius), $a_{\text{IL-7}}$ is the pathway activation rate and $d_{\text{IL-7}}$ is the pathway deactivation rate. In simulations where we used a sink term in the reaction-diffusion equation for extracellular IL-7 (Eq. 8), the sink term for wildtype cells was given by the first term of (Eq. 9):

$$s_{\text{SINK}} = [\text{IL-7R}]\langle[\text{IL-7}_{\text{ex}}]\rangle a_{\text{IL-7}} \qquad \text{(Eq. 10)}$$

In wildtype cells, the concentration of IL-7 receptor [IL-7R] was set to a random value in the range [0, 1) in unscaled concentration units. This amount of IL-7 receptor was set at the moment the cell entered the simulation via cell influx, and was inherited clonally to daughter cells. In previous work we examined the impact of this modelling choice on proliferation and differentiation, both of which depend on IL-7 signaling. In short, clones with higher IL-7 tended to proliferate more and differentiate into $\gamma\delta^{+}$ T-cell subtypes, while clones with lower IL-7 tended to proliferate less and differentiate into $\alpha\beta^{+}$ T-cell subtypes [Aghaallaei et al. 2021]. The model does not include any dynamic feedback mechanisms acting on the receptor concentration, which is a limitation that could be addressed in future work.

In this work, we introduced lesioned cells with impaired IL-7 signaling. If the lesion involved expressing a dominant-active IL-7R receptor, the following alternative functional form for the pathway activation $\sigma_{\text{IL-7}}$ was used

$$\frac{\mathrm{d}}{\mathrm{d}t}\sigma_{\text{IL-7}} = [\text{IL-7R}] - d_{\text{IL-7}}\sigma_{\text{IL-7}} \qquad \text{(Eq. 11)}$$

Correspondingly, the sink term for these lesioned cells was formulated as

$$s_{\text{SINK}} = [\text{IL-7R}] \qquad \text{(Eq. 12)}$$

The level of IL-7 receptor expression of cells with a dominant-active receptor lesion was set to 1 unscaled concentration unit, which is the maximum attainable in the wildtype population.

Cells with an overexpression lesion used (Eq. 9) and (Eq. 10) like wildtype cells, but their IL-7 receptor level was set to 10 unscaled concentration units, which is 10-fold the maximum attainable in the wildtype population.

#### VII. Subcellular Scale: Intercellular Notch signaling

Based on expression patterns in medaka fish thymus [Bajoghli et al. 2009] all TECs in the model express the Notch ligand Dll-4a, while all thymocytes express the Notch receptor Notch1b. Because concentrations of ligand and receptor on the cell membrane are not known, we set the concentration of Dll-4a on TECs to [DLL4a] = 1 unscaled concentration units and the level of Notch1b receptor on thymocytes to [Notch1b] = 1 unscaled concentration units. We do not include feedback mechanisms that dynamically regulate the expression levels of the ligand or the receptor; this is a limitation of the model that could be explored in future work.

Since Notch signaling is mediated by direct cell-cell contact, we used the cell-neighbour pair information obtained from the cellular scale biomechanical interaction model (Eq. 3) to determine if a ligand-expressing TEC was in contact with a receptor-expressing thymocyte.

#### VIII. Subcellular Scale: Intracellular Notch signal transduction

The model for intracellular Notch signal transduction is summarized in Figure A4 B. The activity of the intracellular Notch signaling pathway is modelled as follows:

$$\frac{\mathrm{d}}{\mathrm{d}t}\sigma_{\text{Notch}} = \begin{cases} 1 & \text{if at least one neighbour is a DLL-4-expressing TEC} \\ -k_{\sigma_{\text{Notch}}}\sigma_{\text{Notch}} & \text{otherwise} \end{cases} \qquad \text{(Eq. 13)}$$

where $k_{\sigma\text{Notch}}$ is a parameter for the pathway deactivation rate which we fitted to experimental data of pharmacological inhibition of the pathway in medaka [Pérez Saturnino et al. 2018]. The pathway's maximum activity level is set to 1 unscaled unit. Thymocytes that have a dominant-active lesion of the Notch pathway always have their pathway activity set to 1.

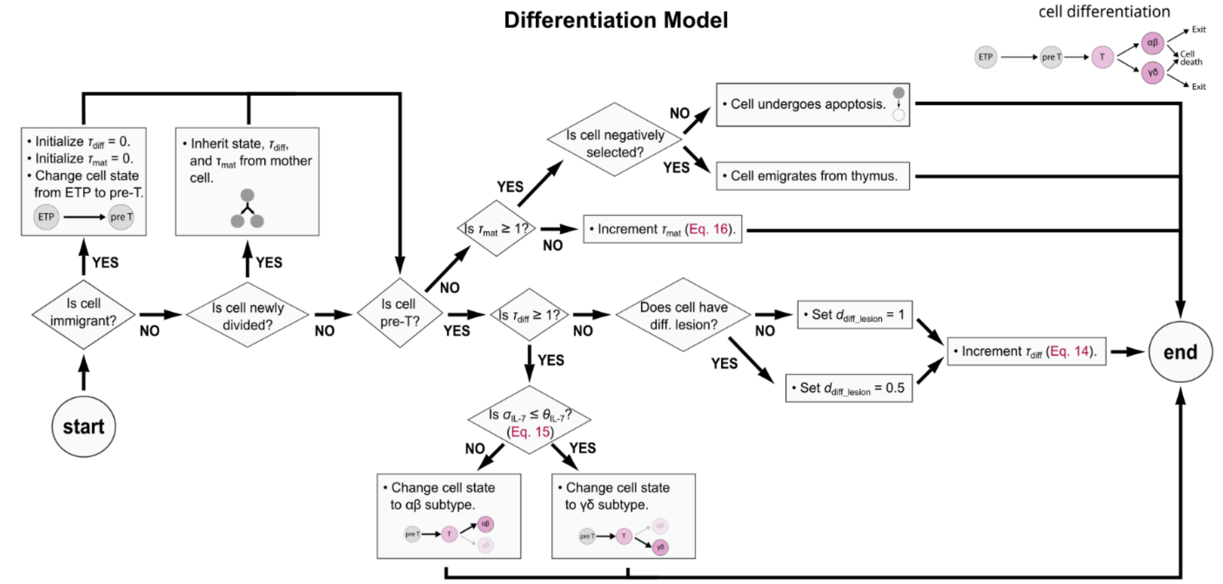


**Appendix Figure 5. Differentiation model.** Schematic flowchart illustrating how cell differentiation stage is updated at every simulation step. Early thymic progenitors (ETPs) entering the thymus for the first time are labelled as "immigrants" and undergo an initialization of variables.

#### IX. Subcellular Scale: Differentiation model

Thymocytes entering the thymus start out as early thymic progenitors in an undifferentiated state; after some time, they can differentiate into either $\alpha\beta^+$ or $\gamma\delta^+$ T-cell subtypes (Appendix Figure 5). To model this process, each cell has an internal variable τdiff that tracks its differentiation progress. A new cell that immigrates into the thymus starts with $\tau_{diff} = 0$ ; the value of $\tau_{diff}$ is then increased at every simulation step, and when $\tau_{diff} \geq 1$ for an individual cell, then this cell's differentiation is considered complete. When a cell divides, both daughter cells inherit the mother cell's value of $\tau_{diff}$. Experimental observations showed that Notch signaling stimulates differentiation [Aghaallaei et al. 2021]. To account for these observations, we model the rate of differentiation (i.e. the rate of increase of $\tau_{diff}$) with

$$\frac{\mathrm{d}}{\mathrm{d}t}\tau_{\mathrm{diff}} = \frac{\kappa+\sigma_{\mathrm{Notch}}(1-\kappa)}{T_{\mathrm{diff}}} \cdot d_{\mathrm{diff\_lesion}} \qquad \text{(Eq. 14)}$$

where $\sigma_{Notch}$ is the Notch pathway activity given by Eq. 13, $\kappa \in [0, 1]$ is a parameter for the rate of Notch-independent differentiation, which we previously estimated at 0.3, and $T_{diff}$ is a parameter for the minimum duration of differentiation, which we previously estimated at 24 hours [Aghaallaei et al. 2021]. The parameter $d_{diff\_lesion}$ was set to 1 for unlesioned wildtype thymocytes, and to 0.5 for thymocytes with a lesion leading to slower differentiation.

The fate selection between $\alpha\beta^+$ or $\gamma\delta^+$ T-cell subtypes depends on a threshold level of IL-7 signaling at the moment of differentiation (i.e. when $\tau_{diff}$ first exceeds 1):

$$\begin{array}{ll} \text{if } \sigma_{\mathrm{IL-7}} \leq \theta_{\mathrm{diff}} & \text{differentiate into } \alpha\beta^+ \\ \text{else} & \text{differentiate into } \gamma\delta^+ \end{array} \qquad \text{(Eq. 15)}$$

In previous work, we studied the effect of altering this threshold level, and identified $\theta_{diff} = 0.4$ as giving a good fit to the wildtype situation [Aghaallaei et al. 2021].

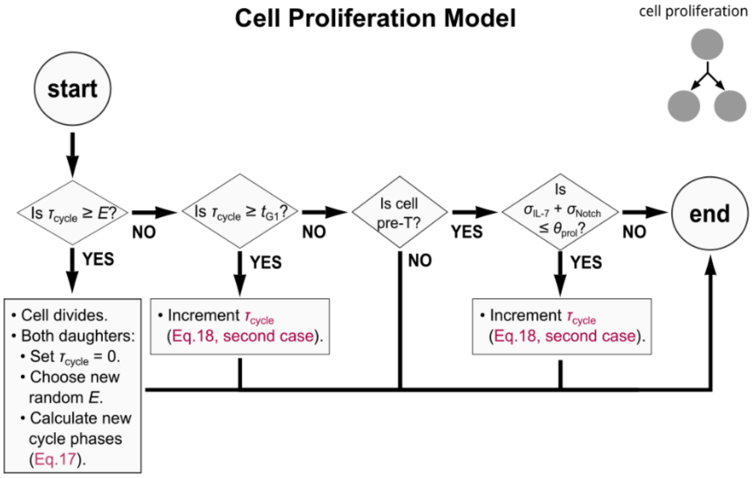


**Appendix Figure 6. Proliferation model.** Schematic flowchart illustrating the cell proliferation model. At each simulation step, cells first check if they completed the cell cycle and divide if yes. Next, if a cell is already past G1-phase, it is committed to finish the cell cycle. Note that this condition is checked before the cell state, hence a differentiated cell could divide once if it differentiated while already committed to the cell cycle. Only undifferentiated cells ("pre-T") can start a new cell cycle, which is conditional on sufficient pro-proliferative IL-7 and Notch signals.

Differentiated $\alpha\beta^+$ and $\gamma\delta^+$ cells linger in the thymus until they mature. Similar to differentiation, we model the maturation process phenomenologically with

$$\frac{\mathrm{d}}{\mathrm{d}t}\tau_{\mathrm{mat}} = \frac{1}{T_{\mathrm{mat}}} \qquad \text{(Eq. 16)}$$

where $\tau_{mat}$ is initially set to 0 and tracks the maturation progress and $T_{mat}$ is a parameter for the minimum duration of maturation, which we previously estimated at 24 hours [Aghaallaei et al. 2021]. When $\tau_{mat} \geq 1$ maturation is considered complete. Once mature, cells are either negatively selected and hence undergo apoptosis, or positively selected and leave the thymus, exiting the simulation.

##### X. Subcellular Scale: Cell proliferation model

The cell proliferation model is summarized in Appendix Figure 6. Thymocytes in the model are only competent for proliferation if they are undifferentiated ($\tau_{diff} < 1$). Each cell draws a random Erlang-distributed random number $E$ with mean $\mu$ = 7 hours and shape value $k$ = 50. This random number is then set as the minimum duration of the next cell cycle – that is, the fastest possible cell cycle if sufficient pro-proliferative signals are present. We subdivide the randomly chosen cell cycle interval $E$ into four phases using the assumption that a cell spends $\kappa_S$ = 50% of the cell cycle duration in the S-phase and $t_M$ = 30 minutes in the M-phase, as follows:

$$\begin{aligned} t_{\mathrm{G1}} &= \frac{E}{2}\left(1 - \frac{\kappa_S}{2}\right) \\ t_{\mathrm{S}} &= E\kappa_S \\ t_{\mathrm{G2}} &= \frac{E}{2}\left(1 - \frac{\kappa_S}{2}\right) - t_{\mathrm{M}} \\ t_{\mathrm{M}} &= t_{\mathrm{M}} \end{aligned} \qquad \text{(Eq. 17)}$$

where $t_{G1}$ is the time spent in G1-phase, $t_S$ the time spent in S-phase, $t_{G2}$ the time spent in G2-phase. The model also includes a G0-like quiescence due to the absence of sufficient pro-proliferative signals. At the beginning of a new cycle, each cell sets the value of its internal variable $\tau_{cycle}$ = 0, which tracks the progression through the cell cycle. This variable is incremented as

$$\frac{\mathrm{d}}{\mathrm{d}t}\tau_{\mathrm{cycle}} = \begin{cases} 0 & \text{if } \tau_{\mathrm{cycle}} \leq t_{\mathrm{G1}} \text{ and } \sigma_{\text{IL-7}} + \sigma_{\mathrm{Notch}} > \theta_{\mathrm{prol}} \\ \frac{1}{E} & \text{otherwise} \end{cases} \qquad \text{(Eq. 18)}$$

where $\sigma_{\text{IL-7}}$ is the IL-7 pathway signaling given by Eq. 9 (or Eq. 11 for clones with dominant-active receptor lesion), $\sigma_{\text{Notch}}$ is the Notch pathway signaling given by Eq. 13, and $\theta_{\text{prol}} = 1.4$ is a threshold parameter.

When $\tau_{\text{cycle}} \geq E$, the cell divides into two daughter cells and for both daughters the value is reset to $\tau_{\text{cycle}} = 0$. Essentially, Eq. 18 models how the IL-7 and Notch pathways act as independent permissive pro-proliferative signals on G1 phase progression; if these signals are absent or too weak, then the cell cycle is delayed as $\tau_{\text{cycle}}$ does not increase and the cell remains in a G0-like quiescent state.

During the initial creation of the model, the sensitivity of the model with respect to the parameter values in Eqs. 17-18 was studied by parameter scan [Aghaallaei et al. 2021]. To choose appropriate values for the mean cell cycle duration $\mu$ and the threshold proliferation-permissive level $\theta_{\text{prol}}$, the cell population size and the percentage of simulated mitotic events at homeostasis was compared to experimental measurements of phospho-histone 3 staining from confocal slices [Aghaallaei et al. 2021].
